# Knowledge-guided analysis of ‘omics’ data using the KnowEnG cloud platform

**DOI:** 10.1101/642124

**Authors:** Charles Blatti, Amin Emad, Matthew J. Berry, Lisa Gatzke, Milt Epstein, Daniel Lanier, Pramod Rizal, Jing Ge, Xiaoxia Liao, Omar Sobh, Mike Lambert, Corey S. Post, Jinfeng Xiao, Peter Groves, Aidan T. Epstein, Xi Chen, Subhashini Srinivasan, Erik Lehnert, Krishna R. Kalari, Liewei Wang, Richard M. Weinshilboum, Jun S. Song, C. Victor Jongeneel, Jiawei Han, Umberto Ravaioli, Nahil Sobh, Colleen B. Bushell, Saurabh Sinha

**Affiliations:** Carl R. Woese Institute for Genomic Biology, University of Illinois at Urbana-Champaign, Urbana, Illinois, 61801; Department of Electrical and Computer Engineering, McGill University, Montreal, Canada H3A 0E9; National Center for Supercomputing Applications, University of Illinois at Urbana-Champaign, Urbana, Illinois, 61801; Department of Computer Science, University of Illinois at Urbana-Champaign, Urbana, Illinois, 61801; Seven Bridges Genomics, Charlestown, Massachusetts, 02129; Division of Biomedical Statistics and Informatics, Department of Health Sciences Research, Mayo Clinic, Rochester, Minnesota, 55902; Division of Clinical Pharmacology, Department of Molecular Pharmacology and Experimental Therapeutics, 55902; Department of Physics, University of Illinois at Urbana-Champaign, Urbana, Illinois, 61801; Cancer Center at Illinois, University of Illinois at Urbana-Champaign, Urbana, Illinois, 61801; Department of Electrical and Computer Engineering, University of Illinois at Urbana-Champaign, Urbana, Illinois, 61801

## Abstract

We present KnowEnG, a free-to-use computational system for analysis of genomics data sets, designed to accelerate biomedical discovery. It includes tools for popular bioinformatics tasks such as gene prioritization, sample clustering, gene set analysis and expression signature analysis. The system offers ‘knowledge-guided’ data-mining and machine learning algorithms, where user-provided data are analyzed in light of prior information about genes, aggregated from numerous knowledge-bases and encoded in a massive ‘Knowledge Network’. KnowEnG adheres to ‘FAIR’ principles: its tools are easily portable to diverse computing environments, run on the cloud for scalable and cost-effective execution of compute-intensive and data-intensive algorithms, and are interoperable with other computing platforms. They are made available through multiple access modes including a web-portal, and include specialized visualization modules. We present use cases and re-analysis of published cancer data sets using KnowEnG tools and demonstrate its potential value in democratization of advanced tools for the modern genomics era.

## Introduction

The rapid growth of genomics data sets^1^ and efforts to consolidate diverse data sets into common portals^2^ have created an urgent need today for software frameworks that can be easily applied to these genomic ‘big data’ to extract biological and medical insights from them^3^. Here, we present ‘KnowEnG’ (Knowledge Engine for Genomics, pronounced ‘knowing’), a cloud-based engine that provides a suite of powerful and easy-to-use machine-learning tools for analysis of genomics data sets. These tools, also referred to as ‘pipelines’, are geared towards data sets represented as spreadsheets or tables (genes × samples) that record typical genomic profiles such as gene expression, mutation counts, etc. for a collection of samples, at the resolution of individual genes. The pipelines help identify biologically meaningful patterns in the provided spreadsheet data, through *ab initio* analysis as well as by contextualizing with prior knowledge. Here, we demonstrate the capabilities of KnowEnG by using it for common bioinformatics analyses such as patient stratification, gene prioritization, gene set characterization and signature analysis on two major data sets in cancer genomics^4,5^, and reproducing key results of the original studies as well as gleaning new biological insights. In doing so, we hope to highlight both the sophisticated level of analysis possible and the ease-of-use with which multiple pipelines can be invoked, individually as well as in combination, to generate a multi-faceted narrative of the insights that the data have to offer.

### Diverse computing environments for KnowEnG

The genomics computing infrastructure of the future has to be adapted to the diverse ecosystem of data sets and tools that will continue to flourish in genomic research. In particular, tools must be *findable, accessible, interoperable and reusable*’^6^, i.e., comply with the ‘FAIR’ principles that guide the modern vision of biological data science. In recognition of these principles, software components of the KnowEnG system are packaged using state-of-the-art technology^7^ that makes them highly portable and amenable to scalable execution in varying computing environments. A convenient way to access the system is through a web portal that links to a KnowEnG server (**Supplementary Note SN1**) running on Amazon Web Services (AWS). A user can upload their genomics data set as a spreadsheet and then execute available pipelines (**Supplementary Note SN2** and **Figure 1A, B**). Often, the results of one KnowEnG pipeline can be further analyzed using another pipeline, and the system facilitates such ‘handover’ between pipelines (**Figure 1D**). For added security and control, users may also create a personal instance of the KnowEnG server and web portal using their AWS accounts (**Supplementary Note SN3**). This design feature can help meet challenges of heavy computing loads faced by a public analytics server. Computationally savvy users may invoke the pipelines and avail of additional functionalities through Jupyter notebooks^8^ from a dedicated KnowEnG server. A third mode of access, created for cancer researchers, is via the NCI Cancer Genomics Cloud Resource built by Seven Bridges (SB-CGC)^9^, where users may directly access large cancer data sets, such as those generated by the NCI TCGA program^10^, and analyze them using KnowEnG pipelines without transferring the data from AWS. Through these varied access modes, KnowEnG facilitates accessibility, interoperability and reusability of its tools, marking a significant step towards realizing the ‘FAIR’ vision.

**Figure 1:**
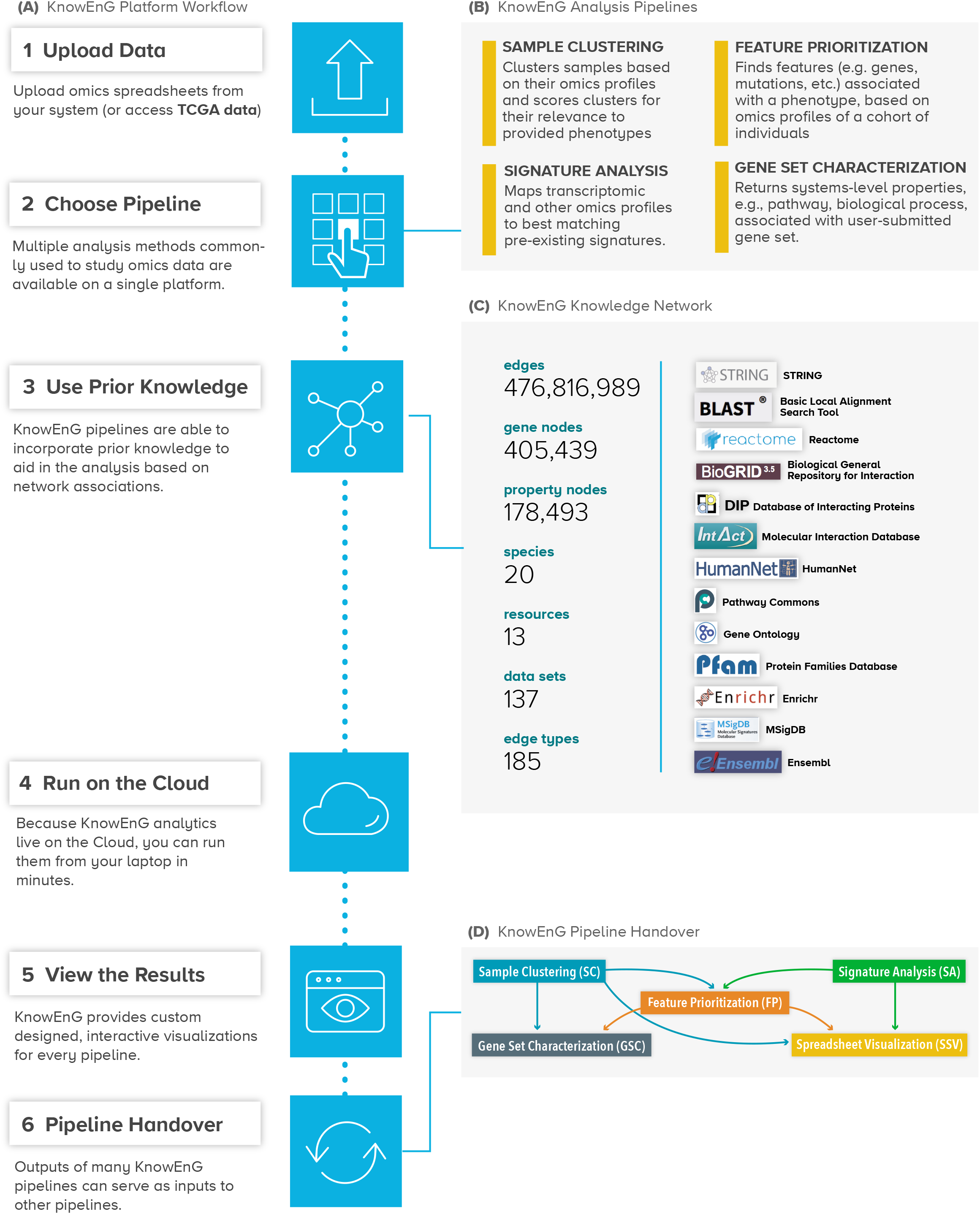
Overview of KnowEnG. **(A)** KnowEnG: portal and system for genomic analysis on the Cloud. **(B)** Analytical functionalities are organized as ‘pipelines’ for common tasks such as clustering, gene prioritization, gene set analysis and signature analysis. Each pipeline offers various options to customize the analysis, including use of prior knowledge. **(C)** Knowledge Network represents prior knowledge that may be used during analysis. Nodes represent genes and biological properties, while edges represent either annotations of gene properties or gene-gene relationships. Sources of information are shown on the right. **(D)** Output of one pipeline may be used as input for another pipeline through a convenient ‘handover’ mechanism in the KnowEnG portal, facilitating deeper and multi-faceted analysis of user’s data.

### Knowledge Network-guided analysis

An important feature of KnowEnG pipelines is that they can incorporate large-scale prior knowledge about genes into analyses of the user’s data set. A basic form of such ‘knowledge-guided’ analysis is already common, where the researcher performs statistical analysis of an experimental data set and then interprets the results in the light of prior knowledge from publicly available gene annotation repositories such as Gene Ontology (GO)^11^, Reactome^12^, etc. KnowEnG makes this analytic process more rigorous by adapting its statistical tools to be directly guided by the vast data in such public repositories. It also breaks the logistical barriers associated with utilizing large databases of prior knowledge, by co-locating its ‘knowledge-guided analysis’ tools with a diverse knowledgebase compiled from numerous popular repositories. The knowledgebase is encoded in a massive heterogeneous network called the ‘Knowledge Network’, whose nodes are genes/proteins and whose edges represent properties (e.g., pathway membership) and mutual relationships (e.g., protein-protein interaction) of the nodes (**Figure 1C**). The network represents annotations of 41 different types from 20 species and 13 different data sources, and includes 476M edges, 405K gene nodes, and 178K property nodes; the network is regularly updated via a ‘one-click’ internal system (**Supplementary Method SM1**). Users typically select the annotation type that is most relevant for guiding their analysis (**Supplementary Note SN4**), in the course of launching a pipeline. The Knowledge Network is also available as a stand-alone resource that allows sub-networks associated with a knowledge type to be retrieved (**Supplementary Note SN5**).

Here, we present the major functionalities, features and interfaces of the KnowEnG system in the context of two previously published and influential cancer data sets. The scope of KnowEnG analytics goes far beyond cancer analysis however, with the system supporting analysis of users’ genomics data from any of ~20 model organisms and its tools being applicable to any data set comprising gene-level measurements or scores for a collection of samples.

## Results

### Case study: Clustering of pan-cancer data

As a first demonstration of the analytic capabilities of KnowEnG, we describe how the ‘Sample Clustering’ pipeline can be used to group genomic profiles in a knowledge-guided manner. Clustering is one of the most widely used tools in bioinformatics^13^ and can help identify sub-groups of samples that represent distinct biological or pathological states^14^; patient stratification in cancer, where subtypes are defined based on molecular markers^15^, is a prime example. The same clustering tools are often applied to different types of genomic profiles, including gene expression, mutation counts, copy number mutations, etc^4^. However, clustering of somatic mutation profiles of cancer patients presents a significant obstacle, since each profile is sparse (a minuscule fraction of genomic loci are mutated) and has little direct similarity to other profiles. As an example of a data set that presents this challenge, we worked with somatic mutation profiles of 3276 tumor samples spanning 12 cancer types (**Supplementary Method SM2**) from the ‘pancan12’ data set generated by the TCGA consortium^4^. (This large data set provides a natural ‘ground truth’, viz., tumor type, for assessing clustering methods.) We first used the ‘standard’ mode of KnowEnG’s Sample Clustering pipeline, viz., Hierarchical Clustering, in six different algorithmic configurations, to identify 14 clusters (so as to match that in the original publication^4^) of tumor samples based on their somatic mutations. (The standard mode of this pipeline also offers K-means clustering.) This failed to produce meaningful clusters, and almost every clustering result exhibited strong ‘resolution bias’^16^, with one cluster comprising over 90% of the samples (**Supplementary Method SM3** and **Supplementary Table SM3.ST1(A)**). The sole exception was clustering with Jaccard similarity and complete linkage^17^, and even here the largest cluster had over 70% of the samples; we will refer to this below as the standard clustering. This initial analysis illustrates the challenge in clustering somatic mutation profiles: due to their high dimensionality and sparsity, biologically related profiles often do not harbor shared mutations and are not grouped together^18^, ultimately leading to many small and one or few large clusters.

### Knowledge-guided clustering of mutation profiles

Knowledge-guided clustering powered by the Knowledge Network offers a possible solution to the problem just noted. Here, prior knowledge of gene-gene relationships encoded in the network is used to recognize when somatic mutations in different genes may be functionally related, thus allowing more subtle forms of similarity between mutation profiles to be exploited in grouping patients. The knowledge-guided option of the Sample Clustering pipeline (**Figure 2A**) implements the ‘Network-based Stratification’ (NBS) algorithm of Hofree et al.^18^, where a random walk method makes patient mutation profiles less sparse by borrowing information from the Knowledge Network before the actual clustering step. We used knowledge-guided clustering with the HumanNet Integrated network (‘hnInt’)^19^ as prior knowledge to group patients into 14 clusters. (Note: All of the main analyses reported in this manuscript can be easily reproduced on the KnowEnG web server by following simple instructions described in **Supplementary Note SN6**.) This yielded more size-balanced clusters; the largest cluster included 30% of the 3,276 patients. To test if patient groups identified from mutation profiles are tied to their phenotypic characteristics, we performed Kaplan-Meier survival analysis (**Figure 2B**). A log-rank test revealed highly significant distinction across the clusters in terms of survival probabilities (p-value 3.7E-33), which was clearly better than that observed in the standard clustering (p-value 7.4E-10, **Supplementary Figure SM3.SF4**). Notably, the original clustering analysis of mutation profiles by Hoadley et al.^4^ was also knowledge-guided, relying on mutations in similar pathways to group related samples, and survival analysis of their original sample clusters produced similarly significant survival distinction (p-value 4.3E-29, **Supplementary Figure SM3.ST6**). The KnowEnG Sample Clustering pipeline, while producing comparable results in terms of survival distinction among clusters, stands out for its ease-of-use compared to executing the multi-step methods of the original analysis. For instance, the user avoids download and harmonization of prior knowledge, installation, and configuration of multiple software, data transformations between steps, and possibly arranging for computing resources capable of compute-intensive steps such as bootstrap sampling (explained below).

**Figure 2:**
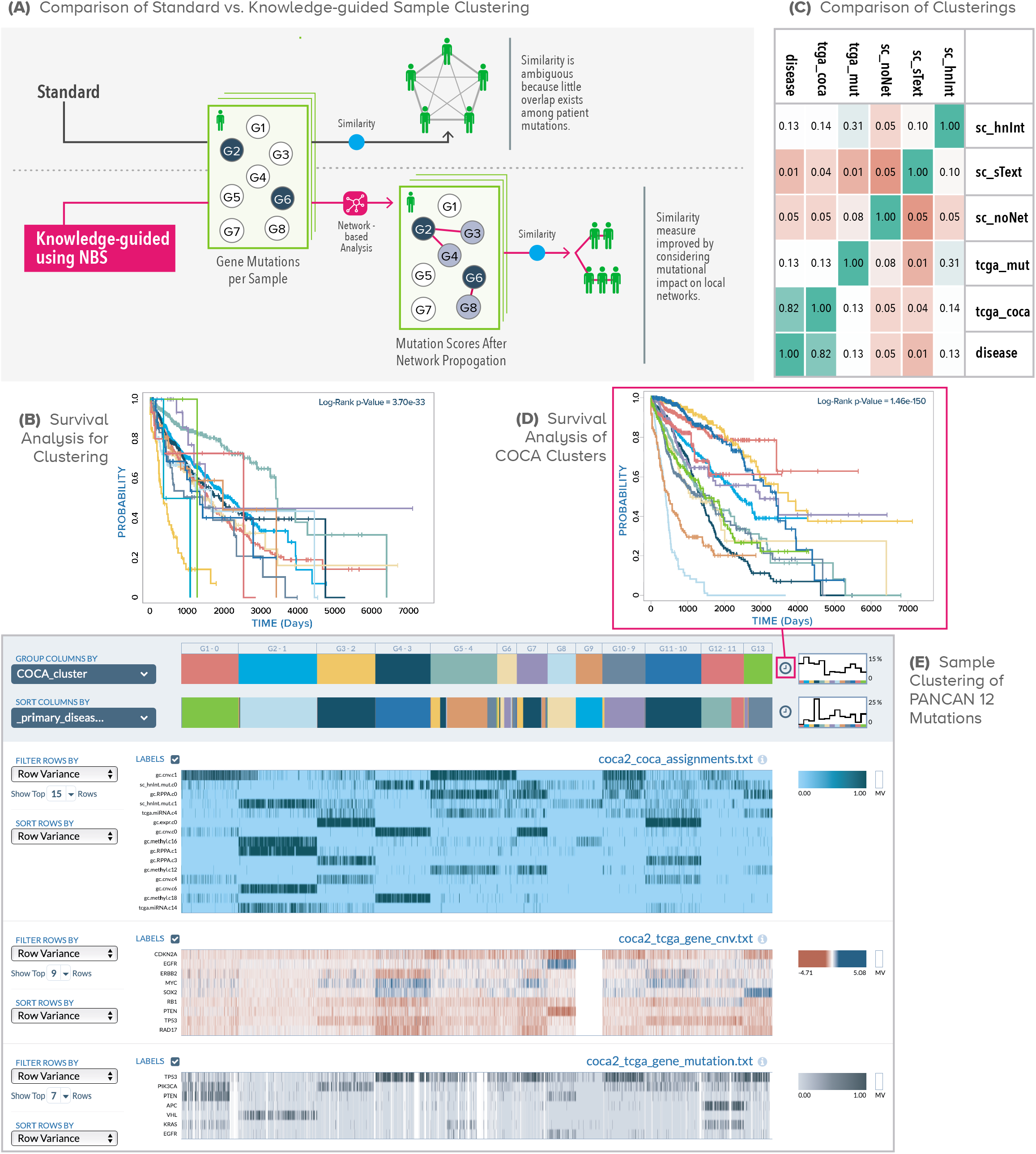
Sample Clustering. **(A)** Knowledge-guided Sample Clustering, illustrated in the context of somatic mutation profiles of cancer patients. Since mutations are rare, two patients may not have mutations to the same gene(s) and their mutual similarity will be modest. In the knowledge-guided mode, similarities between patient profiles are detected not only if the same genes are mutated but also if genes located proximally on a network are mutated; this ‘relaxed’ notion of mutation profile similarity leads to improved clustering. **(B)** Kaplan-Meier survival analysis of clusters from HumanNet-guided clustering of somatic mutation profiles. Each of 14 reported clusters is plotted as a separate survival curve, and the p-value of the multivariate log rank test is displayed. **(C)** Concordance between different clustering approaches, using Adjusted Rand Index (ARI). Three of these approaches use the Sample Clustering (‘sc’) pipeline, with HumanNet (‘hnNet’), STRING text-mining (‘sText’) or no network (‘noNet’) for guidance. Two clustering approaches are reproductions from the Hoadley et al. (‘tcga_mut’ obtained from mutation data, and ‘tcga_coca’ obtained from multi-omics data using COCA). The sixth clustering (‘disease’) is simply a grouping of patients by tumor type. **(D)** Kaplan-Meier survival analysis of 13 COCA clusters in pan cancer multi-omics data. Users may click the clock icon next to cluster assignments to access this display, which uses the current grouping criterion (configurable) for survival analysis. **(E)** Sample Clustering of pan cancer multi-omics profiles, displayed by the Spreadsheet Visualizer module. Patient profiles are grouped by overall cluster assignment using COCA. The top heatmap (blue) shows cluster assignments based on individual omics data types (‘expr’: expression, ‘RPPA’: proteomic, ‘CNV’: copy number variation, ‘methyl’: methylation, ‘miRNA’: microRNA). The heatmaps below show CNV data for select genes (middle) and mutation data for select genes (bottom), for the same patients. Users can configure the number of rows to display for each data source, the statistical criteria for selecting rows and their sorting order. The grouping criteria for samples (COCA cluster assignments here) can also be configured. User-selected clinical annotations of patients (primary disease in this view; color bar second from top) may also be displayed.

Delving deeper into the patient clusters obtained above, we asked whether the clusters recapitulate the tumor types of patients or whether they reveal new structures in the data. To this end, we calculated the adjusted rand index (ARI)^20^ between the clusters and tumor types and repeated the process for other approaches to sample clustering, including the multi-omics Cluster-Of-Cluster-Assignment (COCA) clustering reported in Hoadley et al.^4^ (**Figure 2C**). Interestingly, while there is a high concordance between tumor type and the COCA cluster labels of Hoadley et al.^4^ (ARI = 0.82), the same is not true for NBS-based clusters from the KnowEnG pipeline (ARI = 0.13) or for the pathway-based clustering of mutation profiles in the original study (ARI = 0.13). In other words, knowledge-guided clustering finds groups of patient mutation profiles that have strong correspondence with survival characteristics yet do not simply track tumor types, suggesting alternative levels of molecular similarity. We explored this possibility in detail (**Supplementary Note SN7**), and found the clusters to be characterized by mutations in genes from specific and distinct pathways, even when they are mixed in terms of tumor type representation.

### Clustering of multi-omics data

The standard clustering pipeline in KnowEnG may be applied to any type of spreadsheet data to cluster a collection of samples, while the knowledge-guided clustering pipeline may be used on any gene-level spreadsheet, where rows represent genes. We showcase this capability by performing ‘multi-omics clustering’ of the same cohort of patients as above. A major advantage of multi-omics profiling of patients is that their mutual relationships and hidden group structures revealed by each data type can be consolidated into a more integrative, higher-level clustering that is more informative than any one type of profile alone. This was demonstrated by Hoadley et al.^4^ through their ‘COCA’ (Clustering of Cluster Assignments) method. Mimicking their approach, we first clustered the above pan-cancer cohort of patients based on their gene expression, methylation, copy number variation, or protein abundance profiles (**Supplementary Method SM3, Supplementary Table SM3.ST1(D)**) separately, using standard clustering. (Knowledge-guided clustering may also be used for all of these profiles except methylation, which is not a gene-level data set.) In addition, we considered our knowledge-guided clustering of mutation data reported above and the miRNA clustering from the original publication^4^, thus arriving at six different ways to partition the cohort into clusters. Each such clustering assigns a cluster identifier to a patient, and we can thus describe the multi-omics profiles of the patient as a succinct ‘meta-profile’ of six cluster identifiers. We then used the standard clustering pipeline on these meta-profiles, arriving at 13 clusters (again mimicking the original published analysis^4^) that capture the six different omics data sets on the same patients. For this step, we employed the ‘bootstrap clustering’ option of the sample clustering pipeline, that typically yields more robust clustering^21^; the ease of employing this powerful feature is another example of value added by a cloud-based infrastructure. The steps where different clustering results were combined into common profiles require manipulations with multiple spreadsheets, each being the result of a separate cluster. These steps, as well as several other common matrix operations, are facilitated by KnowEnG through its so-called ‘mini pipelines’ that are available as notebooks in a Jupyter environment (**Supplementary Method SM4**).

### Interactive visualization

Results of the above multi-omics cluster analysis were visualized via the ‘Spreadsheet Visualizer’ module of KnowEnG (**Figure 2E**), which in addition to displaying multiple spreadsheets as a ‘heat map’, allows users to simultaneously visualize various other properties of samples (e.g., cluster assignments provided by COCA, selected clinical annotations such as age, survival months, and primary disease type), offers different ways of sorting, filtering and grouping the data, and provides useful descriptive statistics such as histograms, in an interactive manner. The interactive visualization also allows us to easily perform survival analysis of the displayed clusters, and we used this feature to find that the new multi-omics clusters are strongly concordant with tumor type (ARI = 0.72) and exhibit differences in survival probabilities (p-value 1.0E-150, **Figure 2D, Supplementary Method SM5**) far more prominently than the mutation-only analyses had revealed. The Spreadsheet Visualizer is a powerful data exploration and preliminary analysis tool in its own right (see **Supplementary Note SN8** for details) and can be utilized independently of the clustering pipeline.

### Clustering for patient stratification

As an illustration of how the Sample Clustering pipeline may be used for patient subtyping^15^, we next clustered breast cancer patients in the METABRIC dataset^22^ based on genes related to the epithelial to mesenchymal (EMT) transition, which is a process involved in metastasis. Following the approach in Emad et al.^23^, we clustered patients into two groups based on the expression of their EMT-related genes (**Supplementary Method SM6**). While standard mode of Sample Clustering did not result in clusters with distinct survival probabilities, the knowledge-guided mode achieved significant Kaplan Meier log-rank p-values using either the STRING^24^ text mining interaction network (‘sText’) (p = 3.1E-4) or the HumanNet hnInt’ network (p = 7.6E-4) (**Supplementary Figures SM6.SF3 and SM6.SF4**).

## Case study: Gene Prioritization for tumor types

A routinely conducted analysis of high-throughput omics profiles is in the determination of genes associated with particular phenotypic conditions or biological processes of interest. Discovery of differentially expressed genes^25^ by contrasting transcriptomic profiles before and after treatment or in case versus control experiments, or of genes whose expression correlates with a numeric phenotype such as drug response^26^ are prime examples. The Gene Prioritization pipeline in KnowEnG offers this functionality, given a spreadsheet of omics data (genes × samples) and a ‘phenotype spreadsheet’ (phenotypes × samples) that represents one or more phenotypic labels for each sample in the omics spreadsheet. As a simple demonstration of this pipeline, we analyzed expression data from tumor samples in the pancan12 data set introduced above, comparing each tumor type with all others using a t-test to identify significant differences in individual gene expression between the groups; this is the standard version of the pipeline (**Figure 3A, Supplementary Method SM7**).

**Figure 3:**
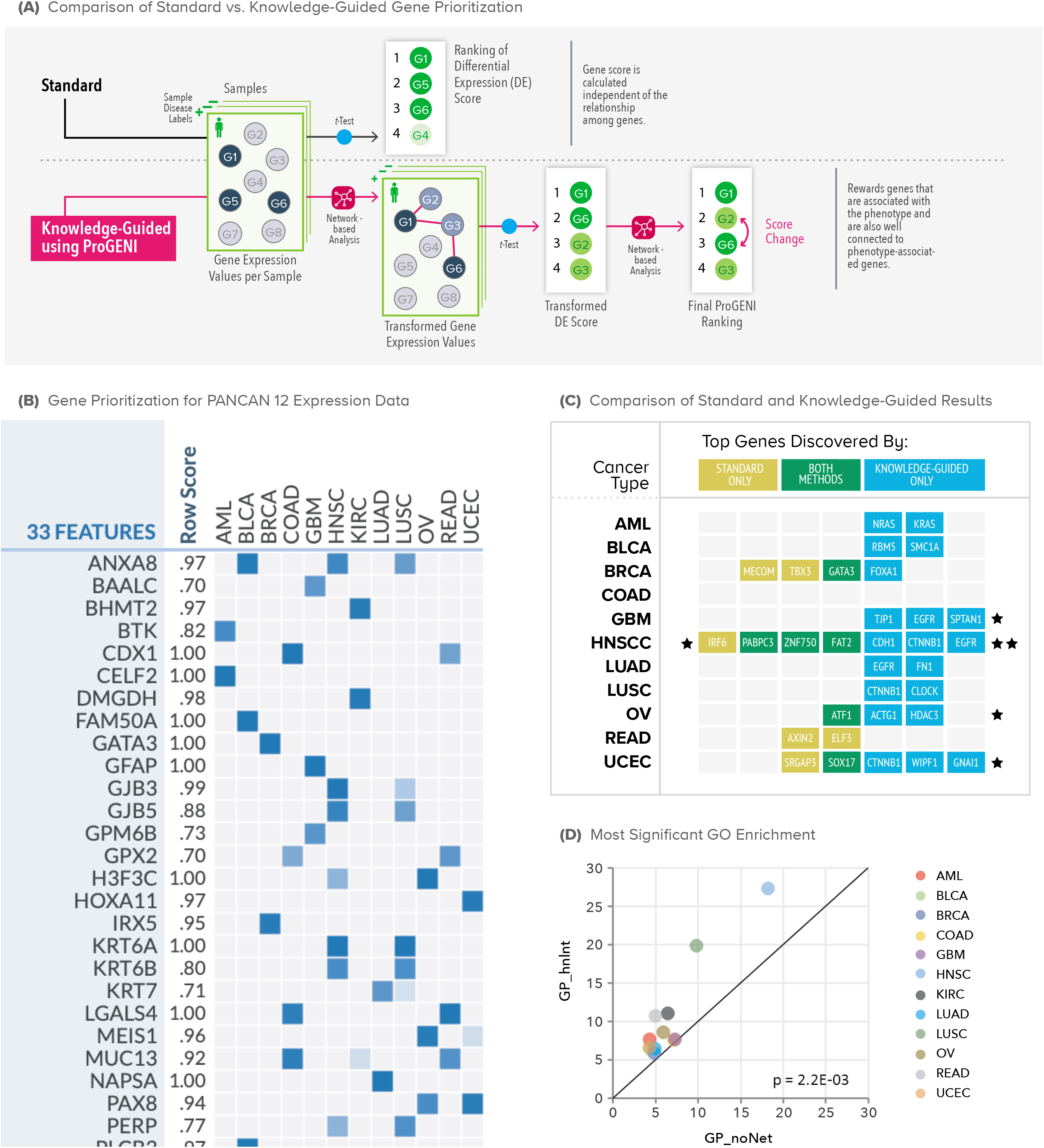
Gene Prioritization Pipeline. **(A)** In standard mode, each gene’s expression is tested for association with phenotypic labels, e.g., with a t-test. In the knowledge-guided mode (ProGENI algorithm [GP]), each gene’s expression is first transformed by taking into account expression levels of its network neighbors, and these ‘network-smoothed’ expression values are tested for association with phenotype. The resulting ranking of genes is subjected to second phase of network-based smoothing to obtain the final ranking. **(B)** Visualization of results from the Gene Prioritization pipeline, used here to identify top genes associated with each tumor type (based on expression data). Users may choose to analyze and visualize results for multiple phenotypes together, and configure how many top genes per phenotype the report should include. **(C)** Known driver genes for each tumor type that are highly prioritized by standard and/or knowledge-guided modes of Gene Prioritization. **(D)** Comparison between tumor type-related genes identified using the Gene Prioritization pipeline in standard mode (‘GP_noNet’) or knowledge-guided mode using HumanNet (‘GP_hnInt’), based on their enrichment for GO terms. The axes represent the negative logarithm (base 10) of p-value of enrichment between the set of highly prioritized genes (from either method) for a tumor type and the most enriched GO category for that set.

### Knowledge-guided gene prioritization

KnowEnG also offers a knowledge-guided mode of this pipeline, where the ProGENI algorithm of Emad et al.^27^ is used to incorporate a network encoding prior knowledge into the identification of phenotype-related genes (**Figure 3A**), using random walk-based techniques similar to those used in the NBS clustering approach^18^. We had previously tested ProGENI on the task of prioritizing drug response-related genes. Through systematic benchmarking, experimental validations and literature surveys we showed that it identifies phenotype-related genes more accurately compared to simple statistical methods as well as machine learning methods that do not utilize prior knowledge^28^. We now applied this algorithm, via the knowledge-guided gene prioritization pipeline, to identify top genes associated with each tumor type, based on expression data (**Figure 3B, Supplementary Method SM7**). (KnowEnG allows this analysis to be performed for all tumor types through one simple operation, rather than repeat it for each tumor type separately.)

### Gene prioritization finds driver genes

For an independent assessment of the above results, we compared the top 100 genes for each tumor type with drivers of that cancer as cataloged in the IntOGen database^29^ based on mutation and gene fusion data (**Figure 3C**). We observed overlaps between the two lists; for example, in head and neck squamous cell carcinoma (HNSCC) six of the highly prioritized genes are known drivers (Fisher’s exact test p-value 8.2E-4, **Supplementary Figure SM8.SF1**). A similar assessment of genes reported by the standard pipeline (without knowledge-guidance) revealed fewer overlaps with respective driver sets for all but two tumor types (**Figure 3C**). Often, common driver genes were identified by both versions of the pipeline, e.g., GATA3 for breast cancer (BRCA), but in many cases the knowledge-guided version reported known drivers that were missed by the standard pipeline, e.g., FOXA1 for BRCA, NRAS, and KRAS for acute myeloid leukemia (AML), and CDH1, CTNNB1 and EGFR for HNSCC. (ESR1, a well known marker of BRCA^30^, was ranked in the top 1.2% of all genes for BRCA, but ranked much worse for other tumor types.) Similar conclusions were reached when we repeated the assessment using a larger external set of tumor type drivers, based on both IntOGen and COSMIC databases^29,31^ (**Supplementary Method SM7**).

### Functional enrichment of prioritized genes

To gain further insights into the highly ranked genes reported for each tumor type in the above analysis, we subjected them to functional enrichment analysis through the Gene Set Characterization pipeline, whose standard version uses the Fisher’s exact test to assess the enrichment of a gene set for pre-specified annotations. This revealed various interesting pathways and Gene Ontology terms as being significantly associated with each tumor type (**Supplementary Method SM8**). For instance, glioblastoma (GBM)-related genes found by ProGENI were significantly associated with receptor proteins in the presynaptic active zone and excitatory synapse, whose altered expression can enhance gliomas ability to grow and survive^32^ (Bonferroni corrected p-value 6.0E-3). Similarly, Acute Myeloid Leukemia (AML)-related genes were enriched for platelet activation, shown to be related to blast proliferation^33^ (Bonferroni corrected p-value 2.0E-6). The extent to which significant functional properties can be associated with a gene set extracted by genomics analyses is one measure of the utility of that gene set^34^. Thus, we summarized the results of gene set characterization by noting the most statistically significant functional enrichment (of genes prioritized) for each tumor type. We noted that when the same process was repeated using genes reported by the standard gene prioritization pipeline the functional enrichments tended to be less prominent (**Figure 3D**), thus providing further evidence of the value of knowledge-guided gene prioritization. The same conclusion was reached when a different network (STRING text mining) was used in gene prioritization instead of the HumanNet integrated network (**Supplementary Method SM8**).

### Pan-cancer signature from prioritized genes

Sets of genes of particular relevance to a tumor type are often used as a ‘signature’ of that tumor, i.e., a representative gene set that captures much of the diagnostic or prognostic value of the entire expression profile. The PAM50 signature of breast cancer is a prime example^15^, being used for patient stratification based on expression of a small set of genes. We asked if the tumor-associated genes prioritized above for each tumor type together form a similar signature with prognostic value in a pan-cancer context. Indeed, we observed that pan-cancer subtypes obtained from clustering only the expression of the tumor-associated genes were just as predictive of survival (Kaplan Meier p-value 3.8E-175) as the above-mentioned clusters based on entire expression profiles (p-value 1.2E-169) (see **Supplementary Note SN9**). This finding was robust to the use of different networks (or no network) in the gene prioritization step.

## Case study: Signature Analysis and Gene Set Characterization on a third-party system

Our next case study makes use of a fourth pipeline – Signature Analysis (**Figure 4A**) – to study a transcriptomic data set of Esophageal Squamous Cell Carcinoma (ESCC) samples^5^, and also showcases how KnowEnG tools can be invoked on computing infrastructures external to the platform (**Figure 4B**). While the KnowEnG web-portal offers a flexible graphical user interface, advanced users performing bioinformatics analysis on a different computing framework may prefer to avail of KnowEnG pipelines on that external framework directly, without tedious transfer of data, intermediate results or code from one system to another.

**Figure 4:**
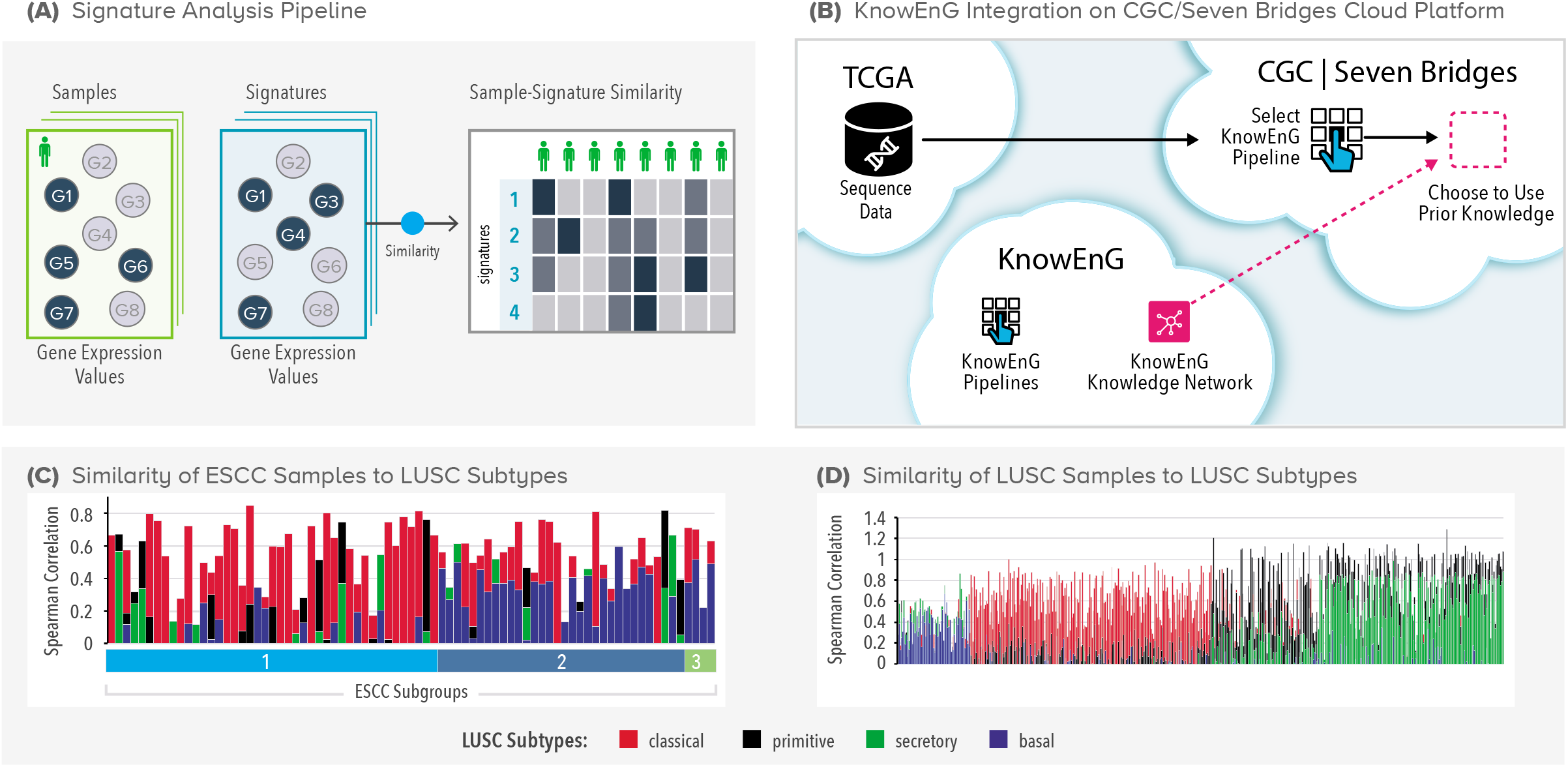
Signature Analysis Pipeline. **(A)** Each user-uploaded expression profile (sample) is matched against expression profiles in a pre-determined collection (signatures) and match scores for all sample-signature pairs are reported by the pipeline. **(B)** Signature Analysis and other KnowEnG pipelines can be executed seamlessly on the third party platform of Seven Bridges Cancer Genomics Cloud (CGC) that hosts a large repository of cancer data and associated tools. The pipelines are published on CGC as a native workflow and the Knowledge Network is transferred ‘under the hood’ from the KnowEnG Cloud when needed by a pipeline. **(C)** Signature analysis of 79 ESCC samples, distributed into three subgroups, matched against four LUSC signatures (subtypes) using Spearman’s Correlation Coefficient. **(D)** Signature analysis of 551 LUSC samples available on the CGC, matched against four LUSC signatures.

### Interoperability

KnowEnG currently offers such seamless interoperability with the Seven Bridges Cancer Genomics Cloud (SB-CGC), which provides researchers with secure access to public data sets such as TCGA and TARGET. We used SB-CGC to access RNA-seq data for the previously reported ESCC tumor samples^5^, and created a transcriptomic spreadsheet (genes × samples) for further analysis with KnowEnG pipelines in the SB-CGC environment (**Figure 4B, Supplementary Method SM9**). This is made possible by the publication of KnowEnG pipelines as native workflows on the SB-CGC, with simple graphical interfaces, and creates opportunities for synergistic use of functionalities offered by these two powerful genomics computing platforms. (External availability of KnowEnG pipelines includes seamless access to the massive Knowledge Network that supports knowledge-guided analysis.) Interoperability is an important tenet of the emerging vision of computing infrastructures of the future. It was achieved by using two emerging technologies – Docker containers^7^ to make the underlying software of each pipeline portable and Common Workflow Language (CWL)^35^ to provide a standardized description of the pipeline (**Supplementary Note SN10**). This alternative mode of KnowEnG usage also facilitates reproducibility and reusability; for instance, users may share their project on SB-CGC with collaborators. Thus, by ensuring interoperability and reusability, in addition to accessibility and findability already offered by the cloud-based web platform, the KnowEnG-CGC joint framework takes a major step towards the realization of the ‘FAIR’ principles of modern data science.

### Signature analysis for patient subtyping

Operating within the SB-CGC framework, we performed a signature analysis of 79 ESCC patients as reported in the original TCGA publication. Signature analysis^36^ is a widely used method in cancer informatics and has been used for various tasks such as identifying subtypes^15^, characterizing purity of tumor samples^37^, determining the abundance of immune cells in tumor microenvironment^38^, characterizing transitions involved in the invasion-metastasis cascade^23^, etc. Here, given a spreadsheet of transcriptomic profiles of a cohort of patients, and a second spreadsheet of pre-determined expression signatures, the pipeline finds the closest matching signature for each patient (**Figure 4A**). This often allows existing insights about the signature to shed light on clinical characteristics of the patient based on their molecular profile. Following in original publication, we matched ESCC samples to signatures representing four subtypes of lung squamous cell carcinoma (LUSC)^39^, since the two cancers are anatomically adjacent and previously established subtypes of LUSC may be relevant to ESCC as well (**Supplementary Method SM10**). We noted that one cluster of ESCC patients (‘ESCC1’, identified in the original publication) mostly (65%) resembled the classical subtype of LUSC, while the second main cluster (‘ESCC2’) mostly (63%) matched the basal subtype of LUSC (**Figure 4C**), and fewer samples matched the primitive and secretory subtypes. The correspondence discovered between *ab initio* detected ESCC subtypes and previously reported LUSC subtypes is generally consistent with the observations of the original TCGA esophageal carcinoma analysis, who note that tumors matching the classical expression subtype also had similar somatic alterations to the subtype and were associated with poor prognosis and chemotherapeutic resistance. To highlight the convenience of co-localizing the analysis workflows with the data on the SB-CGC, we reran the analysis by simply substituting an alternate TCGA dataset of LUSC tumor samples, again finding the classical subtype (40%) to be the most prevalent (**Figure 4D**).

### Pathway analysis of subtype-associated genes

Having categorized ESCC patients into one of four subtypes using signature analysis, we next used the standard gene prioritization pipeline to identify genes associated with each subtype, and subjected the resulting subtype-associated gene lists (**Supplementary Method SM11**) to further analysis using the gene set characterization pipeline introduced above. We now used the knowledge-guided version of this pipeline, which instead of performing the traditional ‘enrichment test’ between sets^40^, uses a random-walk algorithm with the user-provided gene set as ‘restart nodes’, to find property nodes of the Knowledge Network that are most related to the given gene set (**Figure 5A**). This class of algorithms has been successfully used to quantify the relationship between network nodes in a variety of domains such as web mining^41^ and social network analysis^42^. The KnowEnG pipeline uses an implementation called ‘DRaWR’ ^43^, the main advantage of which compared to enrichment tests is that it examines not only properties with which the given genes are annotated, but also the properties with which genes related to the given genes are annotated (**Supplementary Method SM11**). We have previously used DRaWR to characterize gene sets in Drosophila development^43^ and cancer^44^. Here, we used the DRaWR-based knowledge-guided gene set characterization pipeline with the HumanNet Integrated network^19^ as the underlying network to identify, for ESCC subtype-related genes, the most related pathways in the Enrichr Pathways Collection^45^. (The pipeline offers several options for the network as well as the properties to be ranked, see **Supplementary Method SM1**.) As a point of contrast, we also analyzed the gene sets with the standard version of the pipeline that uses the traditional Hypergeometric test approach^46^. **Figure 5C** tabulates 12 discovered pathway associations for ESCC subtypes that were reported by the DRaWR-based version of the pipeline, but not by the standard version. Even though these associations do not meet the traditional criterion of significant set overlap, there is support in the literature for seven of the 12 associations. Moreover, the top-ranked association was between basal subtype of ESCC and the gastric cancer network, which is credible given the close relationship between ESCC and gastric cancer (GCA), which are anatomically adjacent and share several risk factors^47^. Surprisingly, this association was not detected by the enrichment test performed in the standard pipeline. Another interesting example is the primitive subtype being linked to FOXM1 transcription factor network, but only by the DRaWR-based pipeline. FOXM1 has been found to be related to ESCC progression^48^ and to be a potential drug target; our finding of a specific association with the primitive subtype of ESCC suggests that the tumor subtype may be an important factor to consider in its therapeutic significance. We also found several subtype-pathway associations reported by both versions of the pipeline (**Figure 5B**). For instance, both the basal and classical subtypes were associated with NRF2 pathway^49^, the secretory subtype was linked to Syndecan-1 mediated signaling event^50^, and the primitive subtype to oxidation by Cytochromes P450^51^. Six of the 13 such associations found by enrichment-based as well as DRaWR-based gene set characterization had circumstantial evidence in the literature.

**Figure 5:**
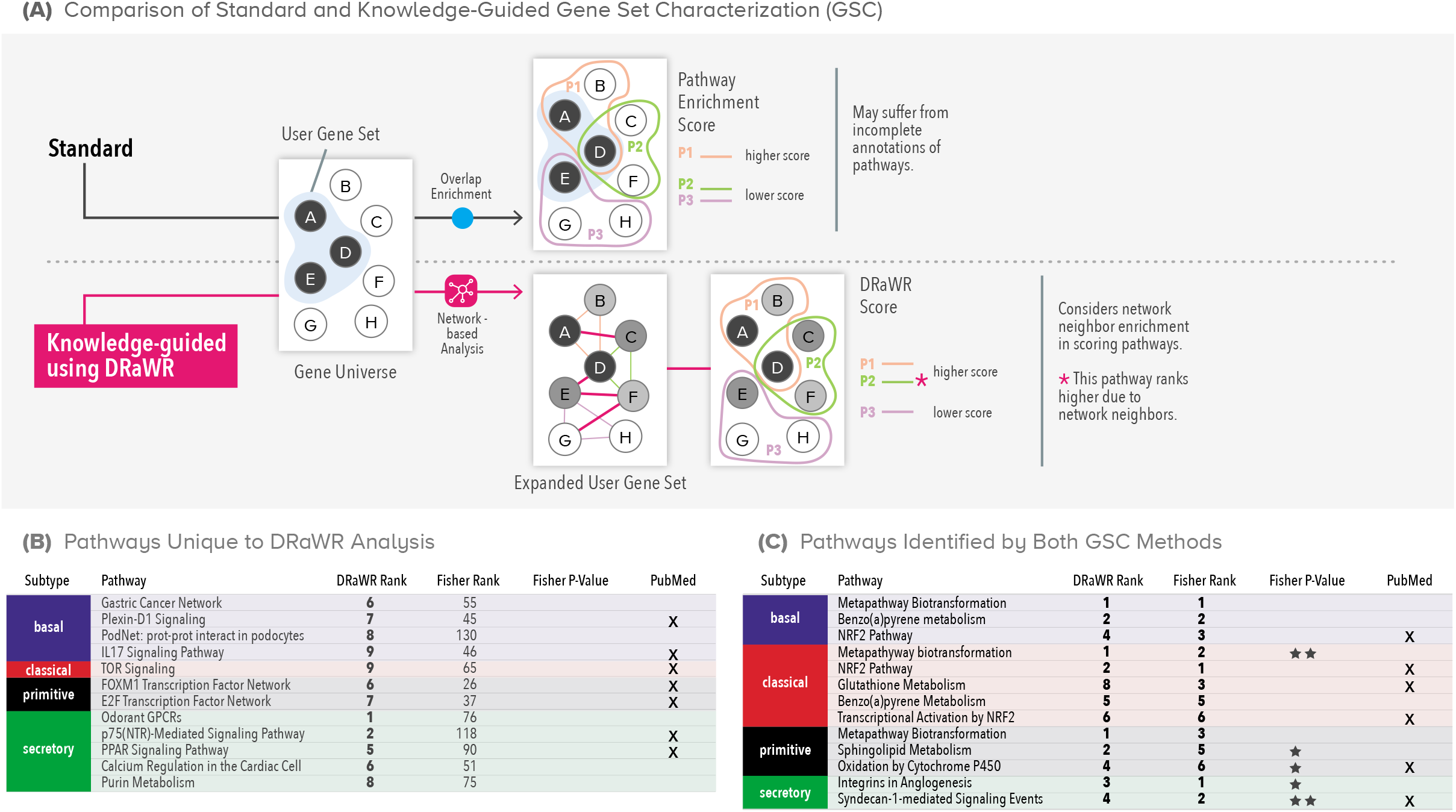
Gene Set Characterization Pipeline. **(A)** Common approaches to gene set characterization (GSC) examine the overlap between a user-provided gene set (e.g., genes A,D,E) and genes in a pathway (e.g., A,D,B in pathway P1). In the knowledge network-guided mode (algorithm DRaWR), the association between two gene sets is based not only on direct overlap between them but also on network-based proximity between them. **(B)** LUSC subtype-associated pathways found exclusively with network-guided GSC pipeline using DRaWR. **(C)** Pathways associated with LUSC subtypes found by standard as well as network-guided GSC pipelines.

In summary, this case study illustrates how different KnowEnG pipelines, in this case, beginning with signature analysis and followed by gene prioritization and gene set characterization, can be used in a workflow to not only relate patient profiles to previously reported cancer subtypes but also to glean novel insights about genes and pathways differentiating patients matched to different subtypes. We performed these analyses on a system external to KnowEnG (i.e., Seven Bridges CGC), but the same workflow may be executed on the KnowEnG platform as well, and the interface facilitates easy ‘stringing’ of multiple pipelines to enable such workflows.

## Discussion

KnowEnG is an analysis engine designed and implemented with the needs and trends of modern genomics research in mind. It embodies some of the most powerful ideas to have emerged in the field over the last decade, including knowledge-guided analysis, cloud-based storage and computing, machine learning and network mining algorithms, and the ‘FAIR’ principles for broader impact. KnowEnG draws inspiration from existing analytic tools and systems, such as geWorkBench^52^, GenePattern^53^, GeneMANIA^54^, etc., and attempts to combine some of their strengths and fill key gaps. For instance, a tool that offers powerful knowledge-guided analytics may be available mainly as a desktop system, with an online version of limited functionality and scalability. On the other hand, some tools provide scalable cloud-based and/or web-based execution but lack knowledge-guided analytical capabilities or only offer analysis of gene sets rather than matrices of omics data. Thus, a joint emphasis on knowledge-guided analysis of rich, spreadsheet-format data sets as well as full-strength online-accessibility and interoperability stands out as a hallmark of the KnowEnG system. Similarly, while tools such as Clustergrammer^55^ and shinyheatmap^56^ offer convenient means for visualization of spreadsheets, akin to KnowEnG’s Spreadsheet Visualizer module, the unique strength of KnowEnG comes from combining the power of interactive visualization with strong analytics. Popular web-based platforms such as cBioPortal^57^, Genomic Data Commons (GDC)^2^ and the UCSC Cancer Genome Browser^58^ also offer useful online analysis of spreadsheet data, but these are typically intended for data sets stored on those portals. In contrast, the main target data for KnowEnG tools are those provided by the researcher, either by direct upload or by selection from an external repository such as SB-CGC.

KnowEnG also offers a vision of genomic computing that is complementary to the dominant paradigm where software packages (e.g., in R or python) are installed on the user’s computer and executed locally. The current paradigm is convenient as long as data sets predominantly reside locally, but with the on-going movement towards massive data sets in the public domain^59^ and a clear need for moving tools to co-locate with these data, we expect the alternative paradigm embraced by KnowEnG to be increasingly relevant. Its main platform provides a convenient way to analyze the user’s uploaded spreadsheets while exploiting massive knowledge-bases encoded in the Knowledge Network, while its interoperability with major cloud-based platforms such as Seven Bridges CGC showcases the advantages of tools moving to data sources while maintaining the convenient ‘illusion’ of local computation. Finally, we note that while the case studies presented above are focused on cancer informatics, the tools of KnowEnG are applicable to a broad array of genomics data sets from a number of different species.

## Methods

The details of the datasets and KnowEnG analysis pipelines used in this article are fully described in the **Supplementary Methods**. The Supplementary Methods also includes additional interpretations for each analyses as well as all of the non-default run parameters needed to reproduce the results. Many subsections contain links to additional resources where the actual code, containers, or compute servers can be found. Additional information about the components of the KnowEnG platform and several related *ad hoc* analyses are also described in detail in the **Supplementary Notes**.

## Data Availability

The datasets analyzed during this study are public Cancer Genome Atlas (TCGA) datasets available from the UCSC Cancer Genome Browser^58^ or the Seven Bridges Cancer Genomics Cloud^9^. The data and parameters for the primary analyses are available in our GitHub repository [https://github.com/KnowEnG/quickstart-demos/tree/master/publication_data/blatti_et_al_2019] (more details in **Supplementary Note SN6**).

## Acknowledgements

We thank our NIH colleagues Ishwar Chandramouliswaran, Valentina di Francesco, Susan Gregurick, and Heidi Sofia for their guidance. We acknowledge the generous resource contributions from the National Center for Supercomputing Application, University of Illinois at Urbana-Champaign (UIUC); Mayo Clinic & Illinois Alliance for Technology-Based Healthcare; Computational Genomics Initiative (CompGen), UIUC; Roy Campbell Systems Research Group, UIUC; NIH-BD2K Common Credits pilot program; Office of Technology Management, UIUC; Cancer Center at Illinois, UIUC. We acknowledge the organizational support from the Carl R. Woese Institute of Genomic Biology, UIUC. We greatly appreciate the assistance from Seven Bridges Genomics Inc, and from the following UIUC personnel and students: Suyang Chen, Joerg Heintz, Henry Lin, Daniel Meling, Shreya Nagesh, Nathan T. Russell, Noor Shalabi, Jackson W.G. Vaughan, Paul Vijayakumar, Svetlana Vranic-Sowers, and Zhuojun Yao. This effort was part of the KnowEng BD2K Center supported by grant U54GM114838 awarded by NIGMS through funds provided by the trans-NIH Big Data to Knowledge (BD2K) initiative.

## Author contributions

Research Design/Consultation: C.B.III, A.E., K.R.K., L.W., R.M.W., J.S.S., N.S., and S.S.; Algorithm Development: C.B.III, A.E., D.L., J.X., J.H., N.S., and S.S.; Knowledge Network: C.B.III, C.S.P, A.T.E., and S.S.; User Interface: C.B.III, A.E., M.J.B., L.G., M.E., X.L., M.L., P.G., C.B.B., and S.S.; Infrastructure Development: M.J.B., M.E., P.R., J.G., X.L., O.S., M.L., P.G., E.L., X.C., U.R., N.S., C.B.B., and S.S.; Documentation: C.B.III, A.E., M.J.B., L.G., M.E., D.L., P.R., O.S., C.S.P, J.X., A.T.E., S.Sr., N.S., and C.B.B.; Manuscript Writing: C.B.III, A.E., M.J.B., L.G., M.E., N.S., C.B.B., and S.S.; Leadership: C.B.III, A.E., M.J.B., O.S., R.M.W., J.S.S., C.V.J., J.H., U.R., N.S., C.B.B., and S.S.

