## Supplementary Methods Document for "Knowledge-guided analysis of ‘omics’ data using the KnowEnG cloud platform"

#### Supplementary Methods 1: Constructing the Knowledge Network

##### Overview

The KnowEnG Center has developed and published a pipeline for collecting and harmonizing prior knowledge about genes and proteins and formatting that data for knowledge-guided analysis pipelines. This pipeline organizes these datasets as a massive heterogeneous network, called the Knowledge Network. The goal of this pipeline is to incorporate as many external sources and make the process of transforming their newest datasets into the Knowledge Network as simple, fast, and transparent as possible (Supp. Fig. SM1.SF1). Roughly, there are four stages to the Knowledge Network Build Pipeline:

1. SETUP, which downloads and imports the latest Ensembl (Zerbino, et al., 2018) gene name mapping databases into a local Redis [https://redis.io/ ] database
2. PARSE, which downloads, parses, and performs entity mapping on the latest version of external public database collections of gene interactions and annotations,
3. IMPORT, which stores the harmonized data and provenance information into a MySQL [https://www.mysql.com/ ] database, and
4. EXPORT, which outputs the interactions and entity maps in the Knowledge Network as flat files formatted for the KnowEnG analysis tools.

The Knowledge Network Build Pipeline is decomposed into many small tasks by species or external public database that we run in parallel across a cloud environment (Supp. Fig. SM1.SF2). We use Apache Mesos [http://mesos.apache.org/ ] to manage the resources and scheduling of these tasks. Apache's Marathon [https://mesosphere.github.io/marathon/ ] framework is given responsibility for deploying the Redis and MySQL databases during the build, and its Chronos [https://mesos.github.io/chronos/ ] framework manages the dependencies between the stages of the build and the many tasks. All tasks are run within an environment provided by our Docker Image, KN_Builder. The build pipeline is designed to run from start to finish with a single command line so it can be automated and regular updates be published to stable location such as AWS S3 [https://aws.amazon.com/s3/ ].

The design philosophy of the Knowledge Network Build Pipeline is to preserve the original annotations and interactions from the public data sources as carefully as possible. Interactions between entities where at least one cannot be mapped with the Ensembl resources are marked and prevented from publication in the final exported Knowledge Network. If multiple evidences exist for the same type of relation, the strongest one is exported and the others are marked as redundant. Metadata relating to the original interaction and the mapping decisions are stored in our MySQL database, as well as the licensing and citation information from the original source. For the final export, subnetworks of a single species and type are reported as 1) a list of weighted, undirected edges using the Ensembl stable identifiers along with references back to their original source line, 2) a map from the stable identifies to common aliases and descriptions, and 3) a file that describes the provenance of the contained data.

#### Methods

For the results in this paper, we performed a complete build of the Knowledge Network in June of 2017, tagged as KN-20rep-1706. This build was focused on twenty species available from Ensembl of particular research interest (Supp. Table SM1.ST1). The build extracted data from 13 external public data sources listed in Supp. Table SM1.ST2. This build extracted 137 different datasets that contained 185 different edge types. The combined Knowledge Network contained 405,439 gene nodes, 178,493 annotation/property nodes, and 476,816,989 edges. In the analysis in this paper, we only use the human subnetworks of the Knowledge Network (taxon id 9606). Specifically, we use the edge types of the Knowledge Network that represent integrated interaction scores from humanNet (hn_IntNet) (Lee, et al., 2011), text mining scores from STRING (STRING_textmining) (Szklarczyk, et al., 2015), annotations from Gene Ontology (gene_ontology) (Ashburner, et al., 2000), and pathway gene sets from Enrichr (enrichr_pathway) (Chen, et al., 2013).

#### Resources

##### Knowledge Network Build Pipeline

Docker Image [https://hub.docker.com/r/knoweng/kn_builder/ ]
GitHub Repository for Launcher [https://github.com/KnowEnG/KnowNet_Pipeline_Tools]
GitHub Repository for Container [https://github.com/KnowEnG/KN_Builder/ ]
Build Pipeline Documentation [http://knowredis.knoweng.org/# ]
Knowledge Network Resources [https://knoweng.org/kn-tools/ ]

##### KN-20rep-1706 Knowledge Network

High Level Content Summary [https://knoweng.org/kn-overview/ ]
Detailed Content Summary [https://knoweng.org/kn-data-references/ ]
Downloadable Content Links
[https://github.com/KnowEnG/KN_Fetcher/blob/master/Contents.md ]

### Supplementary Methods 2: Description of the TCGA PANCAN Dataset

#### Methods

In our first two case studies, we performed our analysis on multi-omics tumor sample data from the TCGA PANCAN analysis of 12 cancer types (Hoadley, et al., 2014). For five types of omics data, we downloaded the omics spreadsheet and the clinical annotations from the UCSC Cancer Genome Browser (Goldman, et al., 2015) in September 2017. The miRNA data from the original study was not available here. We extracted the overall survival days and indicator data from the clinical annotations, "_OS" and "_OS_IND". We also relied on the three clusterings of the samples from the original TCGA publication: their disease annotation ("_primary_disease"), their pathway guided clustering ("_PANCAN_mutation_PANCAN"), and their Cluster-Of-Cluster-Assignments clustering ("_PANCAN_Cluster_Cluster_PANCAN"). Information about these omics files, the number of samples, and the number of omics features can be found in Supp. Table SM2.ST1.

For the first case study, we examined the sample mutation data in particular, which contained mutation data for 39,674 genes on 3276 samples (Supp. Table SM2.ST2). For every (gene, sample) pair in this genomic spreadsheet, there is a "1", indicating that in that sample there was at least one nonsynonymous somatic mutation in the coding sequence of that gene or there is a "0", indicating otherwise. For the Cluster-Of-Cluster-Assignment analysis, we also used the omics data of four of the remaining data types ("CGA_PANCAN12_RPPA_RBN-2015-01-28","TCGA_PANCAN12_exp_HiSeqV2-2015-01-28","TCGA_PANCAN12_hMethyl-2015-01-28","TCGA_PANCAN12_genecopynumber-2015-01-28") and the clustering information from the clinical annotations for miRNA, "_PANCAN_miRNA_PANCAN". The number of clusters in each of the original six clusterings performed by TCGA can be seen in Supp. Table SM2.ST1. In the second case study for gene prioritization, we focused on the population-normalized log2(RPKM + 1) gene expression data from TCGA_PANCAN12_exp_HiSeqV2-2015-01-28 for 16,114 genes in 3598 samples.

### Supplementary Methods 3: KnowEnG Sample Clustering and its Application to TCGA Mutation Dataset

#### Overview

The KnowEnG Sample Clustering pipeline is a data-mining tool that performs the clustering of samples based on the similarity of their feature profiles. This type of analysis is useful in bioinformatics when given a large collection of samples described by their "omics" (genomic, transcriptomic, etc.) profiles and you want to find a limited number of subtypes (or sample clusters). The Sample Clustering pipeline is available in two modes: 1) a "knowledge-guided" mode that integrates gene-based feature profiles with prior knowledge from the KnowEnG Knowledge Network and 2) a "standard" mode that performs the more common without the prior knowledge. In both modes, these omic subtypes may then be correlated with the phenotypes of the samples in order to understand and use them for various prediction tasks (e.g. prognosis, treatment outcome). The Sample Clustering pipeline supports the common paradigm of consensus clustering (Monti, et al., 2003), where subsets of the data are clustered independently many times and the global co-occurrence of samples in the same cluster are tracked. This mode is compute intensive, but it allows for a more robust clustering of the data that is less reliant on the starting or convergence conditions of an individual clustering.

##### Knowledge-Guided Sample Clustering

The knowledge-guided mode of the Sample Clustering pipeline is inspired by Network Based Stratification (Hofree, et al., 2013). The fundamental idea of this analysis is that it is difficult to accurately assess the distance between samples in sparse genomic datasets in which the value assigned to most genomic features for a sample is zero (e.g. a dataset that records the genes with somatic mutations in cancer patients). The hypothesis is that a clustering similarity measure becomes more reliable when you allow the sparse sample features to propagate across local neighborhoods in a known gene-gene interaction network. The KnowEnG Sample Clustering pipeline enables users to run this network-guided analysis on their gene-level omics spreadsheets with many different types of interaction networks.

##### Standard Sample Clustering

The KnowEnG Sample Clustering pipeline supports k-means and hierarchical clustering of the sample columns of the omics spreadsheet in its standard mode. In this mode, the features are not required to be related to genes and the Knowledge Network is ignored.

#### User Inputs

The Sample Clustering pipeline has two primary inputs:

1. a required “omics spreadsheet” which will have its samples divided into subtypes/clusters based on the distance between their feature profiles, and
2. an optional “phenotype spreadsheet” that will contain numerical or categorical information on the samples that will be used to describe how well the identified clusters associate with known phenotypes of interest.

The “omics” spreadsheet is expected to be a matrix of numeric values with named rows representing features and named columns representing samples (Supp. Fig. SM3.SF1). *For the knowledge-guided mode of Sample Clustering, the features must correspond to genes and be labeled with common names or identifiers, and the matrix must contain only non-negative values*.

The phenotypic spreadsheet is expected to be a matrix where the named rows represent the samples of the genomic spreadsheet, and there is one named column for each observed phenotype (Supp. Fig. SM3.SF2). The Sample Clustering pipeline will only consider categorical and numerical phenotypes. More information about the formatting of these input files can be found at our data preparation resource [https://github.com/KnowEnG/quickstart-demos/blob/master/pipeline_readmes/README-DataPrep.md ].

#### User Parameters

There are a number of parameters that the user must select to run the Sample Clustering Pipeline.

##### Global Parameters

All modes of Sample Clustering require the user to select the

- [number_clusters] to group the samples into and specify

If consensus clustering is desired, the user must also set the parameters for

- [number_bootstraps] - an integer value
- [bootstrap_percentage] - percent of samples to randomly select in each bootstrap

##### Knowledge-Guided Only Parameters

In the prior knowledge guided mode, the user must specify the

- [species] - out of the twenty species in the Knowledge Network of the gene features
- [interaction_network] - the gene-gene network available in the Knowledge Network for that species to use in the transformation of each sample's features
- [network_percentage] - controls the extent of influence the interaction network has in the transformation

##### Standard Clustering Only Parameters

In the standard clustering mode, the user must specify the

- [clustering_algorithm] - currently supports k-means or hierarchical clustering

If hierarchical clustering is selected, the user must also supply the

- [affinity_metric] - currently supports Euclidean, Manhattan, and Jaccard distances
- [linkage_criteria] - currently supports clustering assignments using average, complete, and Ward linkages

#### Data Preprocessing

Once the user selects the Sample Clustering inputs and parameters, a simple preprocessing step occurs before the main algorithm. First, columns with missing values and row features with duplicate names in the omics spreadsheet are dropped from the analysis. If the user supplied a phenotype spreadsheet, then there is a quick check that the sample names in the "omics" and phenotypic spreadsheets overlap, otherwise an error is returned.

##### Knowledge-Guided Only

In the knowledge-guided mode, the numeric values of the spreadsheet must be non-negative. The input gene names and identifiers are first mapped to stable Ensembl identifiers of the appropriate [species] using the KN_Mapper tool (see Supp. Note SN5) and the Redis database of gene aliases that accompanies the current Knowledge Network build. Unmapped rows (either missing or ambiguous mappings) are dropped along with the rows that contain a duplicated mapped gene identifier. Finally in this mode, the gene identifiers of the spreadsheet are compared to the gene identifiers of the selected [interaction_network]. If there is no intersection between these identifier lists, then the algorithm is terminated.

#### Description of Algorithm

##### Standard Sample Clustering

Depending on the [clustering_algorithm], the standard clustering mode of the KnowEnG Sample Clustering pipeline uses the sklearn implementations of k-means [http://scikit-learn.org/stable/modules/generated/sklearn.cluster.KMeans.html ] and hierarchical clustering [http://scikit-learn.org/stable/modules/generated/sklearn.cluster.AgglomerativeClustering.html ] at its core. These methods are run with the default settings and the user supplied [number_clusters], [affinity_metric], and [linkage_criteria]. If bootstrap consensus clustering is not used, then this core clustering result is returned as the final clustering assignments. If [number_bootstraps] is greater than one, then our consensus clustering implementation creates that many copies of the omics spreadsheet, each with only [bootstrap_percentage] of the samples, and the core clustering algorithm is repeated on each. The consensus matrix is built by counting the number of times each pair of samples occurred in each bootstrap clustering. Finally, the final cluster assignments are found by one last application of sklearn's k-means on the consensus matrix.

##### Random Walks with Restart

Many of the KnowEnG knowledge-guided analysis methods rely on the basic guilt-by-association method known as Random Walk with Restart (RWR). This method belongs to a family of algorithms that model how information spreads within a graph or network that includes the personalized PageRank algorithm (Page, et al., 1999). The idea is that information on the nodes of a graph start in a seed state. Node values are updated using some percentage of information that is allowed to propagate across the network neighbors and the remaining percentage from the original seed state. For well-connected, aperiodic networks, this update procedure can be repeated and will converge to a stationary node state where the values on the nodes no longer change significantly. These final stationary node states represent a blending of the original seed and the network information.

In KnowEnG, any [interaction_network] from the Knowledge Network can be used in the RWR algorithm. The weighted interaction network is treated as undirected and converted to a transition probability matrix, T, that captures the probability of following a weighted edge from every starting node to every target node in one step in the undirected network. A sample's original gene feature vector is treated as the seed state over all nodes, $V_0$. The final network-smoothed vector, $V_F$, is calculated using steps of a random walk with restart where the vector at each iteration, $V_i$ is defined by:

$$V_i = a * T * V_{i-1} + (1-a) * V_0$$

where $a$ is the [network_percentage] user parameter that dictates the amount of influence the network propagation has on the final result. When the smoothed node state vector converges to a set tolerance, the final vector can be substituted as the description of the sample in further machine learning algorithms.

##### Knowledge-Guided Clustering

The knowledge-guided clustering mode of the Samples Clustering pipeline supports the same single clustering or consensus clustering structure of the standard mode, but replaces the core clustering method (Supp. Fig. SM3.SF3). This implementation is based off of the algorithm proposed in (Hofree, et al., 2013) and a more detailed description can be found there. The first step of this new core clustering method is to transform each sample's original gene feature vector into a network-smoothed feature vector using random walks with restart (RWR). These transformed, non-negative feature vectors are represented in a genes by samples matrix. The clustering of the samples in the knowledge-guided mode applies Non-negative Matrix Factorization (NMF) (Lee and Seung, 1999; Li and Ngom, 2012) to approximate the network-smoothed feature matrix, S, as the product of two reduced-dimension matrices, W and H, where S = WH (Cai, et al., 2008). The two factorization matrices will have reduced dimension size to the [number_clusters], and the largest value across the k components for each sample in the H matrix will determine the cluster that sample is assigned. The implementation of NMF in our code follows the method used by (Hofree, et al., 2013), but converges more quickly because we halt iteration once the clustering becomes stable rather than when the change in ||WH - S|| becomes arbitrarily small.

##### Phenotype Correlation

If a phenotype matrix is supplied, then the final clustering clusters are evaluated for associations with the supplied phenotypes. Each phenotype is classified as categorical or numerical, and the appropriate test statistic and significance is calculated using scipy functions. For numerical phenotypes, we return the significance p-value of a one-way ANOVA [https://docs.scipy.org/doc/scipy-0.18.1/reference/generated/scipy.stats.f_oneway.html ] that tests the null hypothesis that all clusters have the same population mean. For categorical phenotypes, we build the observed contingency table and return the significance p-value of a chi-squared test [https://docs.scipy.org/doc/scipy-0.15.1/reference/generated/scipy.stats.chi2_contingency.html ] for the null hypothesis that the categories and the clusters are independent.

#### Pipeline Outputs

##### KnowEnG Platform Interface

Running the KnowEnG Sample Clustering pipeline in the KnowEnG Platform will produce results that can be interactively viewed using our Cluster visualization tool. This tool includes the same functionality as the Spreadsheet Visualizer (see Supp. Note SN8), with the addition of default settings specific to clustering. The original ”omics” spreadsheet is shown as a heatmap, and the columns are grouped by their cluster assignment and sorted by their similarity to the cluster mean (silhouette scores). The omics row features are sorted by the degree of relevance to the cluster assignments. The user can resort these by feature variance. If multiple bootstraps were performed, the consensus matrix is shown below the omics data. Here the matrix shows samples by samples with each cell representing the consensus score. Single-row heatmaps showing phenotypes and/or specific data features can be displayed in alignment with the other visuals and by default include a score indicating correlation to the cluster groupings. These scores can be recalculated for other groupings on the fly. Many additional features can be used to study the data distributions and the relationships between the clustering, the sample phenotypes, and the “omics” features. Survival curves can be generated for each cluster as well as any categorical phenotype, if this serial time/event information is included in the data.

##### Downloadable Files

The primary downloadable files of the Sample Clustering pipeline are the sample cluster assignments and the consensus matrix that is constructed when multiple bootstraps were performed. There are also silhouette scores provided to indicate cluster quality as well as any phenotype associations if a phenotypic spreadsheet was provided. The cluster centers are reported as a separate output, as well as the top 100 highest scoring gene features for each, which can be passed directly to Gene Set Characterization. *In knowledge-guided mode, additional files containing metadata about the [interaction_network], data preprocessing, and pipeline run are also provided.* More information about the outputs of the pipeline and their structure can be found at [https://github.com/KnowEnG/quickstart-demos/blob/master/pipeline_readmes/README-SC.md].

#### Methods

We applied the Standard Clustering mode of the KnowEnG Sample Clustering pipeline to the spreadsheet of somatic mutation data from the TCGA PANCAN12 study (see Supp. Methods SM2). We ran [clustering_algorithm] hierarchical clustering several times using different combinations of [affinity_metrics] and [linkage_criteria] without consensus clustering. All runs were performed with [number_clusters] set to 14, the number of clusters returned in the somatic mutation analysis of the original TCGA publication (Hoadley, et al., 2014). The Docker image tags at the time of the analysis were knowengdev/data_cleanup_pipeline:10_12_2017 and knowengdev/general_clustering_pipeline:04_22_2018. The results of these runs are summarized in Supp. Table SM3.ST1 (A) and Supp. Fig. SM3.SF4. The clinical data that accompany the TCGA data were also uploaded in the analysis, and the Kaplan-Meier plots and p-values were calculated on the "_OS" and "_OS_IND" fields with Spreadsheet Visualizer. The best standard clustering result was with Jaccard affinity and average linkage, which we call "sc_noNet".

We then applied the knowledge-guided mode of the KnowEnG Sample Clustering pipeline to the same genomic and phenotypic spreadsheets. This analysis was performed twice with two different [interaction_networks]. The first network, "HumanNet Integrated Network" [https://s3.amazonaws.com/KnowNets/KN-20rep-1706/userKN-20rep-1706/Gene/9606/hn_IntNet/9606.hn_IntNet.edge ], was the 469,784 edge, 15,999 node integrated network from the HumanNet project (Lee, et al., 2011) built from their method for combining edge types. The second network, "STRING Text Mining from Abstracts" [https://s3.amazonaws.com/KnowNets/KN-20rep-1706/userKN-20rep-1706/Gene/9606/STRING_textmining/9606.STRING_textmining.edge ], was the 8,511,457 edge, 18,146 node network from the STRINGdb project (Szklarczyk, et al., 2015) that aggregates gene co-occurrence across species in literature abstracts. Both knowledge-guided runs were performed without consensus clustering, with 14 [number_clusters] as in the original paper, and with the [network_percentage] fixed at 50%. The Docker image available at the time of the run was knowengdev/samples_clustering_pipeline:04_09_2018. Of the original 39,674 features, 8,483 were removed due to ambiguous/incomplete name mapping and/or duplication. The results of these runs and the Kaplan-Meier p-values from the Spreadsheet Visualizer are summarized in Supp. Table SM3.ST1 (B) and listed in Supp. Table SM3.ST2. Plots for these runs are found in Figure 2C and Supp. Fig. SM3.SF5.

We next examined the consistency of our best standard clustering result, "sc_noNet", our two network based clusterings, "sc_hnInt" and "sc_sText", and the three prior clusterings/groupings of the samples from the original TCGA paper (Hoadley, et al., 2014). These baseline groupings were extracted from the clinical data: 1) "disease" from their disease annotation ("_primary_disease"), "tcga_mut" from their pathway guided clustering ("_PANCAN_mutation_PANCAN"), and "tcga_coca" from their Cluster-Of-Cluster-Assignments clustering ("_PANCAN_Cluster_Cluster_PANCAN") (Supp. Table SM3.ST1 (C) and Supp. Table SM3.ST2, and Supp. Fig. SM3.SF6). We calculated the similarity between every pair of clusterings with the adjusted Rand Index score [http://scikit-learn.org/stable/modules/generated/sklearn.metrics.adjusted_rand_score.html ] and report the results in Figure 2D. We also compared the clusters of our best network-informed clustering, "sc_hnInt" to the previously discovered clusters from the TCGA paper using the same one-sided Fisher's exact test available through the KnowEnG implementation of the standard Gene Set Characterization pipeline. The negative log10 p-values of enrichment between the pairs of clusters are shown in Supp. Table SM3.ST3.

Finally, we performed the Standard Clustering mode of the KnowEnG Sample Clustering pipeline to each of the four remaining data types (expr, methyl, cnv, and RPPA) for which we had omics spreadsheets (see Supp. Methods SM2 and Supp. Table SM2.ST1). We ran [clustering_algorithm] hierarchical clustering using 'euclidean' [affinity_metrics] and 'Ward' [linkage_criteria] without consensus clustering. All runs were performed with the number of clusters found in the original paper (Hoadley, et al., 2014) for the corresponding data type. The results of these runs are summarized in Supp. Table SM3.ST1 (D). These four clusterings are referred to as "gc_cnv", "gc_expr", "gc_methyl", and "gc_RPPA".

#### Resources

##### Sample Clustering Pipeline

KnowEnG Platform Tool [https://platform.knoweng.org/static/#/pipelines/sample_clustering ]
Quickstart Guide [https://knoweng.org/wp-content/uploads/2017/08/SC_Quickstart.pdf ]
YouTube Tutorial [https://www.youtube.com/watch?v=nAZzN100B3w ]
Data Preparation Guidelines [https://github.com/KnowEnG/quickstart-demos/blob/master/pipeline_readmes/README-DataPrep.md]
Downloadable Results Description [https://github.com/KnowEnG/quickstart-demos/blob/master/pipeline_readmes/README-SC.md ]

##### Seven Bridges Cancer Genomics Cloud

Public Tool [https://cgc.sbgenomics.com/public/apps/#mepstein/knoweng-samples-clustering-public/ ]
Quickstart Guide [https://knoweng.org/wp-content/uploads/2018/07/SC_CGC_Quickstart.pdf ]

##### Docker and GitHub Repositories

Knowledge-Guided Docker [https://hub.docker.com/r/knowengdev/samples_clustering_pipeline/]
Knowledge-Guided GitHub [https://github.com/KnowEnG/Samples_Clustering_Pipeline ]
Standard Clustering Docker [https://hub.docker.com/r/knowengdev/general_clustering_pipeline/]
Standard Clustering GitHub [https://github.com/KnowEnG/General_Clustering_Pipeline ]
Cluster Evaluation Docker [https://hub.docker.com/r/knowengdev/clustering_evaluation/ ]
Cluster Evaluation GitHub [https://github.com/KnowEnG/Clustering_Evaluation ]
Data Cleanup Docker [https://hub.docker.com/r/knowengdev/data_cleanup_pipeline/ ]
Data Cleanup GitHub [https://github.com/KnowEnG/Data_Cleanup_Pipeline ]
Pipeline Utilities Docker [https://hub.docker.com/r/knowengdev/base_image/ ]
Pipeline Utilities GitHub [https://github.com/KnowEnG/KnowEnG_Pipelines_Library ]

### Supplementary Methods 4: KnowEnG Jupyter Tools and Combining Multi-omics Clusterings of TCGA Data

#### Overview

JupyterHub notebooks provide users with a web-browser interface for Python kernel code running on the cloud server hardware, allowing users to conveniently from a local machine complete big data computation on the server where the data is stored and analysis is executed. Notebooks avoid the need to move data through slow network connections and allow the use of cloud resources to complete data-wrangling tasks. The current collection of KnowEnG notebooks provide the means to perform basic spreadsheet matrix transformations such as transposing, selecting, thresholding and more specific, complex data manipulations without requiring a user to understand programming or transfer the data to a local spreadsheet software.

##### User Experience

KnowEnG provides a JupyterHub [https://jupyterhub.readthedocs.io/en/stable/] server instance that gives user access to secure notebooks for data manipulation tasks. Login to the JupyterHub server is handled by the open source access management platform CILogon [https://www.cilogon.org/home] that allows users to researchers to use login credentials from their home institutions or other existing research platform.

The JupyterHub server is separate from the KnowEnG platform, so users wishing to switch between the two must currently manually download and upload the files between servers. Inputs and outputs are (.tsv) tab separated value spreadsheet files. Once on the landing page of the Jupyter notebook, users may click on the 'user_data/' directory, and then use the file upload button on the upper right of the page to upload spreadsheet files. By clicking on the folder icon, the user can return to the landing page and start the transformation notebook by clicking the Spreadsheets_Transformation.ipynb file. Once all the cells of the notebook are running, selection of input files and data transformations is accomplished by using the labeled controls. An hourglass icon will appear in the figure tab when calculations are running, and when finished the results will appear in the 'results/' directory. More detailed instructions and examples can be found are the following resource: [https://github.com/KnowEnG/Spreadsheets_Transformation/blob/master/docs/notebook_Readme.md ].

The containerized versions of the spreadsheet transformations may be run wherever other pipeline containers are run if security requirements allow. The Docker images for the single-user local-server are located at: [https://hub.docker.com/r/knowengdev/spreadsheets_transformation]. Users wishing to set up their own multi-user JupyterHub instance may use the following Docker image: [https://hub.docker.com/r/knowengdev/jupyter_notebooks]

#### Description of Algorithm

The data transformations available in the current version of the KnowEnG Spreadsheets_Transformation.ipynb notebook include:

| | |
|---|---|
| Category to Binary | convert single spreadsheet column to multiple (one per label) binary columns |
| Transpose | rotate the matrix such that rows become columns and columns become rows |
| Intersect | extract the rows in common from two spreadsheets into a new spreadsheet |
| Merge | combine all the rows and columns in two spreadsheets in a new spreadsheet |
| Select Rows | extract smaller spreadsheet from a spreadsheet and a list of rows |
| Average on Labels | process spreadsheet and sample labels to find averages per label |
| Subset on Label | extract smaller spreadsheet that contains only samples with phenotype label |
| Misc Transforms | take absolute value, z-transform, log-transform, or threshold of spreadsheet |
| Descriptive Stats | report min, max, sum, mean, median, stand dev, and variance by row or column |

### Methods

From the UCSC Cancer Genome Browser PANCAN12 TCGA clinical data column, "_PANCAN_miRNA_PANCAN", we extracted the miRNA clusterings in the original paper (Hoadley, et al., 2014). This clusterings was saved as a two column file, the first column with the TCGA sample identifiers and the second with the clinical data column cluster names.

We also downloaded the sample cluster assignment file, “sample_labels_by_cluster.txt”, from five runs of the Sample Clustering pipeline that we performed on the remaining five omics data types. Standard clustering with ’euclidean’ [affinity_metric] and ‘Ward’ [linkage_criteria] was used to produce the clusterings for “gc_methyl”, “gc_expr”, “gc_cnv”, and “gc_RPPA”. Knowledge-guided clustering using the HumanNet Integrated network, “sc_hnInt” (see Supp. Table SM3.ST1), was used for gene-level somatic mutation data. More details about these five clusterings can be found in Supp. Methods SM3.

Once we had gathered the files for all six data types, we uploaded them to the “user_data/CLUSTERS/” directory available in the KnowEnG Jupyter notebook hub server and removed the example files. We then started the “notebooks/Spreadsheet_Transformation.ipynb” notebook, ran all cells, and selected the “Category to binary” transformation mini-pipeline selecting “CLUSTERS” and “process directory” as the two parameters. This converted our six clustering files into a single clusters by samples matrix with binary 0/1 values indicating the sample’s membership in the cluster. This output file, “CLUSTERS_categorical_binary.tsv”, was found in the “/results/” directory of the notebook interface. We moved this output to the “user_data/” along with a list of the 3524 samples used in the COCA analysis and reran the “Select Rows” cell and launched the minipipeline with the two files to remove the unwanted TCGA samples. Then, we moved the resulting output, ”CLUSTERS_categorical_binary_select_rows.tsv”, to “user_data/”, reran the “Transpose” cell, and launched that minipipeline with this file to reorient the final output matrix, "CLUSTERS_categorical_binary_select_rows.tsv". Finally, we downloaded this spreadsheet further analysis in the KnowEnG Platform. We renamed the cluster feature names for clarity, and the resulting table is available in Supp. Table SM4.ST1.

#### Resources

Public Tool [https://knowtebook.knoweng.org/hub/login ]
Quickstart for JupyterHub
[https://github.com/KnowEnG/Spreadsheets_Transformation/blob/master/docs/notebook_Readme.md ]
Spreadsheet Transformation Docker Repository
[https://hub.docker.com/r/knowengdev/spreadsheets_transformation ]
Spreadsheet Transformation GitHub
[https://github.com/KnowEnG/Spreadsheets_Transformation ]

### Supplementary Methods 5: COCA Analysis of Multi-omics TCGA Data

#### Methods

We next attempted to recreate the Cluster-Of-Cluster-Assignment (COCA) analysis from the TCGA PANCAN analysis of 12 cancer types (Hoadley, et al., 2014) using the KnowEnG Platform. To do this, we simply need to submit the matrix we constructed in Supp. Methods SM5 to the standard mode of the Sample Clustering pipeline and use the consensus clustering feature. The multi-omics matrix we created in Supp. Methods SM5 was a clusters by samples matrices with 0/1 membership indicator values that contained clusters from six different types of omics data. This matrix was a "recreation" of the original's papers COCA analysis except using the results of the KnowEnG Sample Clustering on each original omics data spreadsheet (except miRNA, which was exactly identical). This "recreation" matrix is saved as Supp. Table SM4.ST1.

We submitted the "recreation" multi-omics matrix to the standard mode of Sample Clustering. We asked for hierarchical clustering to produce 13 [number_clusters] using the 'euclidean' [affinity_metrics] and the 'ward' [linkage_criteria]. We performed these runs with consensus clustering, with 200 [number_bootstraps] bootstraps, sampling 80% [bootstrap_percentage] of the tumor samples for each one. This is similar to the analysis in the original paper, which found 13 clusters using consensus clustering with hierarchical clustering and 80% sampling percentage. In the original paper, they used Pearson correlation as the affinity metric and ran the method for 1000 bootstrap iterations. The Docker image tags at the time of the analysis were knowengdev/data_cleanup_pipeline:10_12_2017 and knowengdev/general_clustering_pipeline:04_22_2018.

The resulting clustering, "recreation consensus COCA" is of equal quality to the original COCA clusters reported in the TCGA PANCAN analysis (Hoadley, et al., 2014). This evaluation was done using the Kaplan-Meier plots (Figure 2E, Supp. Fig. SM5.SF1) and p-values (Supp. Table SM3.ST1 (E)) calculated from survival analysis on the "_OS" and "_OS_IND" clinical annotation fields.

### Supplementary Methods 6: KnowEnG Sample Clustering and its Application to METABRIC Transcriptomic Dataset

#### Methods

##### Data Collection

Following (Emad, et al., 2017), we obtained a list of 253 genes that have been previously shown to be involved in the epithelial to mesenchymal transition (EMT) from (Taube, et al., 2010). In addition, we downloaded the transcriptomic and clinical data corresponding to the METABRIC study (Curtis, et al., 2012) from OASIS [http://oasis-genomics.org/] and the supplemental material of the original paper. We focused our analysis on the 1058 samples with gene expression and overall survival information. In all analysis, the log2 transformed and Z normalized probe intensities were used. Fig SM6.SF1 shows the Kaplan-Meier survival analysis of different PAM50 subtypes in this dataset.

##### Clustering of samples based on the EMT signature

We used standard mode of the Sample Clustering pipeline in KnowEnG to cluster samples above into two groups using only the expression of the EMT genes (henceforth the EMT signature). We ran [clustering_algorithm] hierarchical clustering using [affinity_metrics] Euclidean and [linkage_criteria] of 'average', 'complete', and 'Ward' without consensus clustering. The Docker image tags at the time of the analysis were knowengdev/data_cleanup_pipeline:01_04_2018 and knowengdev/general_clustering_pipeline:04_22_2018. Of the three clusterings obtained, two (obtained using average and complete linkage) were significantly biased: one cluster contained only one sample and the other cluster contained all other samples. Hence, we focused on results obtained using the 'Ward' linkage that provided less biased results. The Kaplan-Meier survival analysis did not show a significant distinction between the survival of the two clusters ($p = 0.12$, log-rank test, Supp. Fig. SM6.SF2), confirming the results of (Emad, et al., 2017).

Next, we sought to determine whether the knowledge-guided Sample Clustering of KnowEnG could improve these results. Since this pipeline only accepts non-negative "omic" input datasets, we obtained the absolute value of the (normalized) EMT gene expression profiles. Intuitively, a value close to zero means that the gene is expressed close to the mean, while a large positive value shows that the gene is highly or lowly expressed compared to other genes (i.e. the gene is differentially expressed). We ran the knowledge-guided Sample Clustering using integrated HumanNet (hnInt) (Fig SM6.SF3) and STRING TextMining (sText) (Fig SM6.SF4). In both cases, the p-value of Kaplan-Meier log rank test was significant: $p = 7.56E-4$ for hnInt and $p = 3.08E-4$ for sText, showing significant improvement compared to the standard mode of clustering.

### Supplementary Methods 7: KnowEnG Feature Prioritization and its Application to TCGA PANCAN12 Transcriptomic Datasets

#### Overview

A widely popular application of high throughput omics profiling is identification of features (e.g. genes, proteins, mutations) likely to underlie particular phenotypic conditions or biological processes of interest. The Feature Prioritization pipeline in KnowEnG offers this general functionality, given a spreadsheet of features (features x samples) and a “phenotype spreadsheet” (phenotypes x samples) that represents one or more phenotypic labels for each sample in the feature spreadsheet. The features may correspond to omics data (e.g. expression of genes, somatic mutations, etc.), but they may also be more general (e.g. clinical features such as age).

##### Knowledge-Guided Feature Prioritization (Gene Prioritization)

One of the major capabilities provided in KnowEnG is knowledge-guided gene prioritization, which identifies genes that are most related to a phenotype of interest. For this mode of operation, the feature spreadsheet should contain gene-level omics measurements. To enable this capability, KnowEnG utilizes a generalized version of a computational tool called ProGENI (Emad, et al., 2017), which we have previously developed. This method is built based on the premise that if known biological relationships of the genes (in the form of a gene interaction network) is properly included in this task, both the direct and indirect influence of the genes on the phenotype can be captured, resulting in a more accurate prioritization. In (Emad, et al., 2017), this method was used to identify genes whose basal mRNA expression determines the sensitivity of cell lines to various drugs. Through extensive evaluations and follow-up siRNA knockdown experiments, we showed that this method significantly improves the accuracy of state-of-the-art gene prioritization methods (Emad, et al., 2017).

##### Standard Gene Prioritization

In cases where the feature spreadsheet does not contain gene-level measurements, this pipeline supports standard feature prioritization. In this mode, the Knowledge Network is not used.

#### User Inputs

The Gene Prioritization pipeline has two primary inputs:

1. a required “feature spreadsheet” that contains measurements corresponding to different features of a collection of biological samples, and
2. a required “phenotype spreadsheet” that contains numerical (e.g., drug response, patient survival, etc.) or categorical (e.g., cancer subtype, metastatic status, experiment vs. control) information on the samples.

The feature spreadsheet should be formatted as a (features x samples) spreadsheet. The first column should contain the feature names and the first row should contain the sample names. If this spreadsheet contains missing values, several options are provided to handle these missing values (see User Parameters). The phenotype data is submitted as a separate spreadsheet formatted as samples x phenotype. The first column should contain the sample names and the first row should contain the name of different phenotypes. The phenotype file can contain continuous-valued, binary, or categorical data. The missing values are allowed in the phenotype spreadsheet. For each phenotype, only samples with values will be used for the prioritization task. For more details about the input spreadsheets, see Supp. Fig. SM3.SF1 and SM3.SF2 in Supp. Methods SM3 and data preparation in: [https://github.com/KnowEnG/quickstart-demos/blob/master/pipeline_readmes/README-DataPrep.md ].

#### User Parameters

After submission of the two spreadsheets above, the user is given various options depending on their data and goal.

##### Global parameters

The user can choose how to deal with missing values in the omics spreadsheet:

- [omics_missing_value] - the options for handling missing values (NA) are
    - Average: Replace the missing value of a feature in a sample with the average expression of the same feature across other samples.
    - Remove: Remove any samples that contain missing values.
    - Reject: Reject the user's spreadsheet if any missing value exists.

Several parameters are needed for the underlying algorithm:

- The user is asked whether they would like to use the Knowledge Network. For the standard Feature Prioritization pipeline the user should choose 'No' and for the knowledge-guided mode 'Yes' .
- [primary_prioritization_method] - options are t-test (for binary) or absolute Pearson correlation (for continuous-valued phenotype). If the standard mode of this pipeline is used, the [primary_prioritization_method] is used for prioritization. In the knowledge-guided mode, [primary_prioritization_method] is only one part of the whole prioritization pipeline. With a categorical phenotype (with more than two categories), a t-test should be chosen in standard Feature Prioritization mode and the pipeline will automatically identify genes related to one category vs. all other categories for every single category.
- [num_exported_features_per_phenotype] - the number of high priority features to return in the output result files that are formatted for further pipeline analysis.

If bootstrap sampling is selected, the following options need to be specified:

- [num_bootstraps] - an integer value
- [bootstrap_percentage] - percent of the samples to be randomly selected in each bootstrap sampling

##### Knowledge-Guided Only Parameters

To choose the network to be used in the analysis, the user should select:

- [species] - out of the twenty species in the Knowledge Network of the gene features.

- [interaction_network] - the gene-gene network available in the Knowledge Network for the selected species to use in the Gene Prioritization

Several parameters are needed for the underlying algorithm:
- [network_percentage] - controls the extent of influence the interaction network has in the transformation
- [num_response_correlated_features] - number of genes to be used as the restart set in the second RWR of the pipeline

#### Preprocessing of the data

Once the user selects the pipeline inputs and parameters, a simple preprocessing step occurs before the main algorithm.

Preprocessing of the feature spreadsheet: First, based on the user's selected option ([features_missing_value]), the missing values in the feature spreadsheet are handled. Then, rows or samples with duplicate names are dropped from the analysis. Also, if a feature name is missing, the corresponding row is removed from the analysis.

Preprocessing of the phenotype spreadsheet: First, there is a quick check to ensure that the sample names in the feature and phenotypic spreadsheets overlap, otherwise an error is returned. Missing values (NAs or NaNs) are accepted in the phenotype spreadsheet. If a phenotype of interest contains those values, samples for which the phenotype value is missing are not included in the analysis. If several phenotypes are provided in the phenotype spreadsheet, each phenotype is handled separately, ensuring that the maximum numbers of samples are used in the analysis. If t-test is chosen, the phenotype spreadsheet must contain phenotypes with at least two distinct categories, otherwise it is rejected. If number of categories is larger than two, then the phenotype spreadsheet is expanded to enable the comparison of each category with all other categories pooled together (one vs. all strategy).

##### Knowledge-Guided Only

In the knowledge-guided mode, the input gene names and identifiers of the feature spreadsheet are mapped to stable Ensembl identifiers of the appropriate [species] using the KN_Mapper tool and the Redis database of gene aliases that accompanies the current Knowledge Network build. Unmapped rows (either missing or ambiguous mappings) are dropped along with the rows that contain duplicated mapped gene identifier. Finally, the gene identifiers of the spreadsheet are compared to the gene identifiers of the selected [interaction_network]. If there is no intersection between these identifier lists, then the algorithm is terminated.

#### Description of Algorithm

##### Knowledge-Guided Mode:

Given the gene-level omics (feature) spreadsheet and the selected [interaction_network], this method first performs a network-based smoothing (using a random walk with restart algorithm, described in Supp. Methods SM3) of the omics data such that the new value assigned to each gene also includes the omics characteristics of the gene's network neighbors, properly weighted to reflect the strength of their relationship. For details of this step, see (Emad, et al., 2017). Then, these smoothed values are compared with the phenotypic labels of the samples and using a statistical test ([primary_prioritization_method]) the top [num_response_correlated_features] genes are selected that are most "associated" (both positively and negatively) with the phenotype. This set of genes is then used as the restart set in an RWR algorithm on the same network, to obtain a score (i.e. equilibrium probability) for all the genes in the network. These scores are then normalized with respect to another score that represents the global topology of the network in order to remove the network bias. This normalized score is used to rank all the genes and identify top genes most related to the phenotype in light of the Knowledge Network. If bootstrap option is selected by the user, the procedure above is repeated [num_bootstraps] times, on randomly selected subsets of the samples (size of the subsets are specified using [bootstrap_percentage]) and the generated ranked lists of the genes are aggregated into a final robust ranking. A complete description of this method is provided in (Emad, et al., 2017).

###### Standard Mode:

In this mode, the method used for feature prioritization is determined by [primary_prioritization_method]. The feature values of the samples are compared to the phenotypic labels of the samples and using a statistical test ([primary_prioritization_method]) features are ranked. If the bootstrap option is selected by the user, the procedure above is repeated [num_bootstraps] times on randomly selected subsets of the samples (size of the subsets are specified using [bootstrap_percentage]), and the generated ranked lists of the features are aggregated into a final robust ranking.

#### Pipeline Outputs

##### KnowEnG Platform Interface

The Feature Prioritization pipeline in the KnowEnG platform produces a visualization component that allows users to interactively view the results of the pipeline. The amount of data being visualized in the heatmap can be controlled by choosing the number of top features displayed per phenotype and turning on and off individual phenotypes. This allows users to both reduce the scale of the heatmap visualization and to focus on those aspects of the analysis that are of most interest. The users are also able to see "top features" score curves for each phenotype, which may help guide the investigation of results. Colored cells in the visualization heatmap represent the score of a given feature/phenotype correlation. Darker colors generally indicate a more significant correlation. Specifically, the darkest color in the color scale is always assigned to the greatest score found among all features for all of the displayed phenotypes. The lightest color in the color scale is assigned to the lowest score among the selected "top features" for all of the displayed phenotypes, and this assignment is updated whenever the user adjusts the number of "top features." Scores less than the lowest "top feature" score are mapped to gray. Interactive sorting options and heatmap cell rollovers further help to illuminate the results.

##### Downloadable Files

The downloadable zip archive contains several files. One file contains only the ranked list of features for each phenotype. Separately for each phenotype, there is a file that contains the ranked list of features and the scores that were used to rank them and several other types of information. Also, one file contains the top ranked [num_exported_features_per_phenotype] features for each phenotype that is formatted such that it can be directly sent to the Gene Set Characterization pipeline. More information about the outputs of the pipeline and their structure can be found at [https://github.com/KnowEnG/quickstart-demos/blob/master/pipeline_readmes/README-FP.md].

#### Methods

In our second case study, we sought to identify genes associated with each tumor type in the PANCAN12 dataset (Hoadley, et al., 2014). We downloaded the expression spreadsheet and clinical annotations in the file "TCGA_PANCAN12_exp_HiSeqV2-2015-01-28.tgz" from the UCSC Cancer Genome Browser (Goldman, et al., 2015) in September 2017. This dataset contained expression of 16,114 genes in 3598 samples. To run the standard Feature Prioritization, we selected the 'average' option for [features_missing_value] to impute missing gene expression values. For each cancer type, the phenotype label of each sample represented whether the sample corresponds to the cancer type of interest ('1') or not ('0'). We used t-test as [primary_prioritization_method] for comparing the expression values and the phenotype and did not use the bootstrap sampling option.

For the knowledge-guided Gene Prioritization, we ran the analysis with both STRING text mining (sText) and HumanNet integrated (hnInt) network for [interaction_network] (see Supp. Methods SM3 for more details on these networks) in separate tests. We selected the probability of restart as 0.5 (i.e. the [network_percentage] equal to 50), used t-test as [primary_prioritization_method] for comparing the smoothed expression values and the phenotype, selected the [num_response_correlated_features] as 50, and did not use the bootstrap sampling option.

The list of top 100 genes for each cancer type is provided in Supplementary Tables SM7.ST1, SM7.ST2 and SM7.ST3 when using the standard Feature Prioritization (noNet), the knowledge-guided prioritization with hnInt and with sText, respectively. Next, we compared the number of top genes (for each cancer type) that are identified using pairs of these methods (Supp. Table SM7.ST4). In average, approximately 52.72% of the identified genes differed between any two methods, with the biggest difference observed between hnInt and sText results with an average difference of 58.25%. Next, we compared the similarity between top 100 genes of any two cancer types using the same method. The results are provided in Supp. Tables SM7.ST5, SM7.ST6 and SM7.ST7 for noNet, hnInt, and sText, respectively. The highest similarity was observed between colon_adenocarcinoma and rectum_adenocarcinoma with 78% using noNet, 71% using hnInt, and 67% using sText. head_&_neck_squamous_cell_carcinoma and lung_squamous_cell_carcinoma also showed a high degree of similarity with 27% using noNet, 31% using hnInt, and 50% using sText.

#### Resources

##### Gene Prioritization Pipeline

KnowEnG Platform Tool
[https://platform.knoweng.org/static/#/pipelines/feature_prioritization]
Quickstart Guide [https://knoweng.org/wp-content/uploads/2017/08/GP_Quickstart.pdf]
YouTube Tutorial [https://www.youtube.com/watch?v=Vp76-Oz-Yuc ]
Data Preparation Guidelines [https://github.com/KnowEnG/quickstart-demos/blob/master/pipeline_readmes/README-DataPrep.md]
Downloadable Results Description [https://github.com/KnowEnG/quickstart-demos/blob/master/pipeline_readmes/README-FP.md]

##### Seven Bridges Cancer Genomics Cloud

Public Tool [https://cgc.sbgenomics.com/public/apps#mepstein/knoweng-geneprioritization-public/gene-prioritization-workflow/ ]
Quickstart Guide [https://knoweng.org/wp-content/uploads/2018/02/GP_CGC_Quickstart.pdf]

##### Docker and GitHub Repositories

Knowledge-Guided Docker [https://hub.docker.com/r/knowengdev/gene_prioritization_pipeline/ ]
Knowledge-Guided GitHub [https://github.com/KnowEnG/Gene_Prioritization_Pipeline ]
Standard Docker [https://hub.docker.com/r/knowengdev/feature_prioritization_pipeline/ ]
Standard GitHub [https://github.com/KnowEnG/Feature_Prioritization_Pipeline ]
Data Cleanup Docker [https://hub.docker.com/r/knowengdev/data_cleanup_pipeline/ ]
Data Cleanup GitHub [https://github.com/KnowEnG/Data_Cleanup_Pipeline ]
Pipeline Utilities Docker [https://hub.docker.com/r/knowengdev/base_image/ ]
Pipeline Utilities GitHub [https://github.com/KnowEnG/KnowEnG_Pipelines_Library ]

### Supplementary Methods 8: Characterization of Top Genes Identified for Each Tumor Type in TCGA PANCAN12

#### Methods

To assess the relevance of the genes identified using knowledge-guided as well as standard modes of Feature Prioritization pipeline for each cancer type (Supp. Methods SM8), we obtained the list of cancer driver genes from IntOGen] [www.intogen.org] (Rubio-Perez, et al., 2015) and COSMIC [https://cancer.sanger.ac.uk/cosmic ] (Forbes, et al., 2017). The list of driver genes from IntOGen, which contained 475 genes, is shown in Supp. Table SM8.ST1. The list of driver genes from COSMIC, which contained 699 genes, is shown in Supp. Table SM8.ST2.

##### Characterization of top genes using cancer-specific driver genes

First, we sought to determine the overlap between top genes identified for each cancer type (referred to as hnInt, sText, and noNet following the notation in Supp. Methods SM8) with the driver genes of the same cancer in IntOGen. IntOGen did not contain driver genes for KIRC. Also, the driver genes for COAD and READ were reported in a single group. Supp. Table SM8.ST3 shows the list of driver genes annotated in IntOGen, separately for each cancer type present in the PANCAN12 dataset. Next, we used the standard mode of KnowEnG Gene Set Characterization pipeline (which implements Fisher's Exact test) to compare the list of top 100 genes identified for each cancer type with the drivers of that cancer (see Supp. Methods SM13 for a description of this pipeline). Supp. Figs. SM8.SF1 and SM8.SF2 show the number of driver genes (and their corresponding p-values) for hnInt vs. noNet and sText vs. noNet, respectively. As can be seen in these figures, for five (out of eleven) cancer types, the enrichment p-values of top genes identified using knowledge-guided analysis (hnInt or sText) are smaller than 0.05, while this is true only for one cancer type when using the standard mode of Gene Prioritization (noNet).

##### Characterization of top genes using all driver genes

Next, we sought to determine the overlap between top genes identified for each cancer type with all the driver genes in IntOGen and COSMIC. Similar to the previous analyses, we used the standard mode of KnowEnG Gene Set Characterization pipeline to compare the list of top 100 genes identified for each cancer type with cancer driver genes. Supp. Figs. SM8.SF3 and SM8.SF4 show the number of driver genes from IntOGen (and their corresponding p-values) for hnInt vs. noNet and sText vs. noNet, respectively. As can be seen in these figures, the enrichment p-value of top genes identified using hnInt for six cancer types and using sText for five cancer types are smaller than 0.05, while this is true only for two cancer types when using the standard mode of Gene Prioritization (noNet).

Supp. Figs. SM8.SF5 and SM8.SF6 show the number of driver genes from COSMIC (and their corresponding p-values) for hnInt vs. noNet and sText vs. noNet, respectively. As can be seen in these figures, the enrichment p-value of top genes identified using hnInt for six cancer types and using sText for five cancer types are smaller than 0.05, while this is true only for two cancer types when using the standard mode of Gene Prioritization (noNet).

##### Characterization of top genes using all driver genes

To gain further insights into the highly ranked genes reported for each tumor type, we performed functional enrichment analysis using KnowEnG's Gene Set Characterization pipeline (standard mode) to identify Gene Ontology (GO) terms most associated with these gene sets. Supp. Tables SM8.ST4, SM8.ST5, and SM8.ST6 show the top 10 enriched GO terms for each cancer type, where the top 100 genes are identified using hnInt, sText, and noNet, respectively. The extent to which significant functional properties can be associated with a gene set extracted by genomics analyses is one measure of the utility of that gene set (Choobdar, et al., 2018). Thus, we summarized the results of gene set characterization by noting the most statistically significant functional enrichment (of genes prioritized) for each tumor type (Supp. Table SM8.ST7). We noted that the enrichments were significantly more prominent when genes were identified using hnInt compared to noNet (p=2.2E-3, Wilcoxon signed rank test), as depicted in Supp. Figs. SM8.SF7 and SM8.SF8. Similarly, the enrichments were significantly more prominent when genes were identified using sText compared to noNet (p=4.4E-3, Wilcoxon signed rank test), as depicted in Supp. Fig. SM8.SF7 and SM8.SF9. These results provide further evidence of the value of knowledge-guided gene prioritization.

### Supplementary Methods 9: Selection of TCGA Data from SB-CGC

#### Methods

In this case study, we sought to recreate part of the genomic subtype signature analysis performed by the Cancer Genome Atlas Research Network in their recent study on the genomic characterization of oesophageal carcinoma (Cancer Genome Atlas Research, et al., 2017). We decided to recreate this analysis, not only with KnowEnG developed tools, but also as demonstration of the portability of those tools. For this reason, all of the analyses in this case study are run with our tools on the Seven Bridges [https://www.sevenbridges.com/ ] NCI Cancer Genomic Cloud (SB-CGC) [http://www.cancergenomicscloud.org/ ].

The first step of this case study was to extract the relevant gene expression datasets from the SB-CGC and reformat them into a transcriptomics spreadsheet to be analyzed by KnowEnG pipelines. We created two datasets, one for Esophageal Carcinoma (ESCA), the original samples studied in the TCGA paper (Cancer Genome Atlas Research, et al., 2017), and the other for Lung Squamous Cell Carcinoma (LUSC), a related cancer type. In order to create each of these datasets, we used the SB-CGC “Data Browser” feature to explore and select the appropriate TCGA RNA-seq data. To do this, a user 1) selects the “TCGA GRCh38” dataset, 2) adds files with “Data category” = “Transcriptome Profiling” and “Data format” = “TXT”, and 3) and selects investigations where the “Disease Type” is one of the two disease types of interest, either ESCA or LUSC. This produces a list of files for each cancer type, which then need to be copied to an SB-CGC project. When we performed these steps, each sample produced three files, ‘*.htseq.counts.tz’, *.FPKM-UQ.txt’, ‘*.FPKM.txt’. For our recreation, we kept only the ‘*.FPKM.txt’ files and deleted the other two types. More detailed information about this selection process can be found here [https://github.com/KnowEnG/KnowEnG_CWL/tree/master/CGC#gathering-the-tcga-input-files ].

This selection process obtained 173 files for ESCA samples and 551 files LUSC samples (Supp. Table SM9.ST1). We then used the KnowEnG Spreadsheet Builder, a custom tool built specifically for converting TCGA files and their metadata on the SB-CGC into omics and phenotypic spreadsheets for KnowEnG Analysis. For each of the two datasets, we ran the Spreadsheet Builder workflow [https://cgc.sbgenomics.com/public/apps#mepstein/knoweng-spreadsheetbuilder-public/spreadsheet-builder/ ] on SB-CGC using the FPKM files as inputs. We ran these workflows using the ‘sample_id’ as the “Metadata Sample ID Key” that matches samples between the omics and phenotypic spreadsheets. We also ran with a Filter Threshold of “1” and a Filter Minimum Percentage of “0.1”, which filtered out genes that did not have a FPKM of at least 1 in at least 10% of samples. After filtering, we were left with 17842 gene features. Finally, we set the Spreadsheet Builder “Normalization Flag” which scales the log2(FPKM +1) to mean zero and standard deviation 1 and outputs these z-scores of the log transformed expression values. Each of these two workflow runs took 5-10 minutes on the SB-CGC, costing ten cents in direct AWS costs, using spot instances. The runs produced a transcriptomic spreadsheet for each cancer type and a phenotypic spreadsheet that contained the corresponding clinical TCGA data. The combined phenotypic spreadsheets are available in (Supp. Table SM9.ST1).

#### Resources

##### Spreadsheet Builder Pipeline

###### Seven Bridges Cancer Genomics Cloud

Spreadsheet Builder Workflow Tool
[https://cgc.sbgenomics.com/public/apps#mepstein/knoweng-spreadsheetbuilder-public/spreadsheet-builder/ ]
Tutorial for Workflow [https://github.com/KnowEnG/KnowEnG_CWL/tree/master/CGC#running-the-spreadsheet-builder-workflow ]

###### Docker and GitHub Repositories

Spreadsheet Builder CWL [https://cgc.sbgenomics.com/raw/mepstein/knoweng-spreadsheetbuilder-public/spreadsheet-builder/1 ]
Spreadsheet Builder Docker [https://hub.docker.com/r/mepsteindr/spreadsheet_preprocess/ ]
Spreadsheet Builder GitHub [https://github.com/KnowEnG/SpreadsheetPreprocess ]
Pipeline Utilities Docker [https://hub.docker.com/r/knowengdev/base_image/ ]
Pipeline Utilities GitHub [https://github.com/KnowEnG/KnowEnG_Pipelines_Library ]

### Supplementary Methods 10: KnowEnG Signature Analysis on LUSC Subtypes

#### Overview

The KnowEnG Signature Analysis pipeline is a basic analysis tool that calculates the similarity between the shared features of a sample and a known omics signature. This type of analysis is useful in bioinformatics when you are given a large collection of samples described by their "omics" (genomic, transcriptomic, etc.) profiles and want to map them to a limited library of subtypes characterized from previous studies.

##### User Inputs

The Signature Analysis pipeline has two primary inputs:

1. A required "omics spreadsheet" that will have its samples mapped to the subtype signatures based on the similarity between their feature profiles, and
2. A required "signature spreadsheet" that contains the omics features and average values of known omics subtypes of interest.

Both the “omics” and “signature” spreadsheet are expected to be a matrices of numeric values with named rows representing features and named columns representing samples or signatures respectively (see Supp. Fig. SM3.SF1). It is important that the measurements represented in the matrices are similarly derived and therefore appropriate for comparison, e.g. both spreadsheets contain gene expression values and were normalized in analogous ways. Also, each feature’s name must match exactly between the omics and signature spreadsheet for that feature to be included in the comparison. More information about the formatting of the input files can be found at our data preparation resource [https://github.com/KnowEnG/quickstart-demos/blob/master/pipeline_readmes/README-DataPrep.md ].

##### User Parameters

There is only a single parameter that the user must select to run the Signature Analysis Pipeline:

- [similarity_measure] to define the similarity between each sample and signature. Currently ‘cosine’ similarity is supported as well as ‘Pearson’ and ‘spearman’ correlation.

##### Data Preprocessing

A simple preprocessing step occurs before the main Signature Analysis algorithm that checks that there are no missing or non-numeric values and then extracts and reorders the shared features between the omics and signature spreadsheets.

##### Description of Algorithm

Once the omics and signature spreadsheet are harmonized with the shared features in the same order, the KnowEnG Signature Analysis pipeline calculates the similarity between every sample in the omics spreadsheet and every signature using the [similarity_measure] selected by the user.

##### Pipeline Outputs

###### KnowEnG Platform Interface

Running the KnowEnG Signature Analysis pipeline in the KnowEnG Platform will produce results that can be viewed interactively. The Signature Analysis visualization, similar to the Feature Prioritization pipeline visualization, provides a panel that enables control of the amount of data being visualized in the primary heatmap result. By filtering on a signature score threshold across samples and turning on and off individual signatures, the user can both reduce the scale of the heatmap visualization and focus on those aspects of the analysis that are of most interest. The user is also able to see score curves for samples by signatures, which may help guide the investigation of results.

###### Downloadable Files

There are two primary downloadable files of the Signature Analysis pipeline. The first is a signatures by samples matrix where each cell contains the calculated similarity value. The second is a samples by signatures matrix of indicator 0/1 values, where a 1 indicates that the signature is the best match for the input sample. More information about the outputs of the pipeline and their structure can be found at [https://github.com/KnowEnG/quickstart-demos/blob/master/pipeline_readmes/README-SA.md ].

#### Methods

For this case study, we wished to reproduce Extended Data Figure 6 [https://www.nature.com/articles/nature20805/figures/12 ] of the TCGA analysis of oesophageal carcinomas (Cancer Genome Atlas Research, et al., 2017). In order to do this, we needed first to retrieve the LUSC genomics subtypes collected for the original analysis. These lung squamous cell carcinoma signatures were created from mRNA expression measured by microarrays in a study by Wilkerson et al. (Wilkerson, et al., 2010). We downloaded the "predictor.centroids" data from this study from the supplementary website [http://cancer.unc.edu/nhayes/publications/scc/]. This file contains four subtypes of LUSC, 'basal', 'primitive', 'classical', and 'secretory' described by their normalized expression values of 208 genes. We used the "File" management features of the SB-CGC to upload this signature spreadsheet into our project.

We next ran the Signature Analysis workflow [https://cgc.sbgenomics.com/public/apps#mepstein/knoweng-signature-analysis-public/ ] in the SB-CGC for each of two transcriptomic samples spreadsheets we had constructed previously, ESCC and LUSC (see Supp. Methods SM10). In the SB-CGC implementation, the workflow maps the gene aliases from both the samples and the signatures spreadsheets to common Ensembl identifiers using the KN_Mapper (see Supp. Note SN5) before running the Signature Analysis pipeline, so we provided the [species_taxon_id] of '9606' for human. For each samples spreadsheet, we provided our LUSC signatures and used the 'spearman' correlation as our [similarity_measure] to run the Signature Analysis pipeline. Each workflow run took under 5 minutes to run and cost five cents using spot instances. Eleven of the original 208 signature genes were unmapped by KN_Mapper.

For the "similarity_matrix" output file from the run with ESCA samples, we extracted 79 samples that were flagged as esophageal squamous cell carcinoma (ESCC) in the original analysis (Cancer Genome Atlas Research, et al., 2017). Note, eleven of the 90 samples in the original analysis are not present in the SB-CGC TCGA dataset. These ESCC samples were placed into one of three subgroups by the original analysis (ESCC1, ESCC2, and ESCC3), which we make available in the first column of Supp. Table SM9.ST1. We removed negative Spearman correlations and plot the correlations of the samples to the 4 LUSC subtypes in the style of Extended Data Figure 6 from the original paper (Cancer Genome Atlas Research, et al., 2017) (see Figure 4C). We also repeated this process from the "similarity_matrix" output file from the run with LUSC samples to produce Figure 4D. For the 5 groups of samples "ESCC1", "ESCC2", "ESCC3", "other ESCA", and "LUSC", we calculated the average non-negative correlation of these samples to each subtype and counted the number of samples with a best match to each subtype (Supp. Table SM10.ST1). The results are not identical to the results presented in the original Extended Data Figure 6 for a few possible reasons: 1) the normalization procedure in the original paper does not match the SB-CGC FPKM data and our normalization, 2) the original figure used Pearson rather than Spearman correlation, 3) a few genes were removed from our correlation calculation due to mapping ambiguities.

We continued the analysis of the 79 ESCC samples by finding out for each tumor subtype, which genes were differentially expressed between the samples that mapped to that subtype and the samples that mapped to other subtypes. This was done using the standard mode of the KnowEnG Feature Prioritization pipeline (see Supp. Methods SM8) on the SB-CGC [https://cgc.sbgenomics.com/public/apps#workflow/mepstein/knoweng-geneprioritization-public/gene-prioritization-workflow ]. We took the assignment matrix of each ESCC sample to its best match signature (similarity_matrix.binary.tsv) from the ESCA Signature Analysis run and filtered for the 79 ESCC samples (Supp. Table SM10.ST2). We then launched the Feature (Gene) Prioritization workflow on the SB-CGC. For the omics spreadsheet input, we used the ESCA transcriptomic samples spreadsheet that was also the input to Signature Analysis. For the phenotypic spreadsheet input, we used the assignment matrix mentioned above. We also provided the species taxonomy identifier again (9606) and requested the 't_test' [primary_prioritization_method]. We chose to not run the knowledge-guided mode, so no network related parameters were specified. The method was also run without bootstrap sampling. We ran this workflow on the SB-CGC, which completed in under 5 minutes and for less than five cents using spot instances.

The result was a list of 100 genes for each of four tumor subtypes reported in the top_genes_per_phenotype_matrix.txt (Supp. Table SM10.ST3). We compared the genes of these four top100 gene sets to each other and to the set of 208 genes from the original LUSC signatures (Supp. Table SM10.ST4). While 31 genes were shared in the top100 prioritized genes for the basal and classical subtypes, the secretory subtype was very distinct with no genes overlapping the other top100 lists. These four top100 lists and the top_genes_per_phenotype_matrix of Feature Prioritization were used as inputs to the Gene Set Characterization workflow described in the next section.

#### Resources

##### Signature Analysis Pipeline

KnowEnG Platform Tool [https://platform.knoweng.org/static/#/pipelines/signature_analysis ]
Data Preparation Guidelines [https://github.com/KnowEnG/quickstart-demos/blob/master/pipeline_readmes/README-DataPrep.md]
Downloadable Results Description [https://github.com/KnowEnG/quickstart-demos/blob/master/pipeline_readmes/README-SA.md ]

##### Seven Bridges Cancer Genomics Cloud

Public Tool [https://cgc.sbgenomics.com/public/apps#mepstein/knoweng-signature-analysis-public/ ]
Quickstart Guide [https://knoweng.org/wp-content/uploads/2018/07/SA_CGC_Quickstart.pdf ]
Combined Workflow Tutorial [https://github.com/KnowEnG/KnowEnG_CWL/tree/master/CGC#running-the-gene-set-characterization-workflow ]

##### Docker and GitHub Repositories

Signature Analysis Docker [https://hub.docker.com/r/knowengdev/signature_analysis_pipeline/ ]
Signature Analysis GitHub [https://github.com/KnowEnG/Signature_Analysis_Pipeline ]
Data Cleanup Docker [https://hub.docker.com/r/knowengdev/data_cleanup_pipeline/ ]
Data Cleanup GitHub [https://github.com/KnowEnG/Data_Cleanup_Pipeline ]
Pipeline Utilities Docker [https://hub.docker.com/r/knowengdev/base_image/ ]
Pipeline Utilities GitHub [https://github.com/KnowEnG/KnowEnG_Pipelines_Library ]

### Supplementary Methods 11: KnowEnG Gene Set Characterization

#### Overview

The KnowEnG Gene Set Characterization pipeline performs the important role in the analysis of gene sets derived from omics data to identify the most related, previously curated pathways, functions, or experimentally annotated genes. This analysis provides researchers with context and information about their gene set of interest that can spur further hypotheses and investigations. The Gene Set Characterization pipeline is available in two modes: 1) a “knowledge-guided” mode that integrates analysis of prior knowledge gene annotations with known gene-gene relationships from the KnowEnG Knowledge Network and 2) a “standard” mode that performs the most common statistical enrichment test.

##### Knowledge-Guided Gene Set Characterization

The knowledge-guided mode of the Gene Set Characterization pipeline is based on the method of Discriminative Random Walk With Restart (DRaWR) (Blatti and Sinha, 2016). The fundamental idea of this analysis is that curated gene set annotations are incomplete, biased, and noisy. By integrating these annotations with additional knowledge about characterized relationships between genes and by calculating network-based distances between gene sets, we can find relevant and informative relationships that would not be captured by standard gene set overlap/enrichment. The KnowEnG Gene Set Characterization allows users to run this network-guided analysis on their gene set with many different types of gene interaction networks.

##### Standard Sample Gene Set Characterization

The KnowEnG Gene Set Characterization pipeline supports the standard statistical overlap test frequently used in gene enrichment analysis. In this mode, the user has access to the extensive collection of gene set annotations in the KnowEnG platform but does not make use of the gene-gene interaction networks from the Knowledge Network.

#### User Inputs

The primary input of the Gene Set Characterization pipeline is one or more gene sets. This can be provided to the KnowEnG platform in either of two ways:

1. For a single gene set, a list of gene symbols can be pasted into the platform with each gene symbol on its own line.

2. For multiple user gene sets, a “spreadsheet” of gene set membership can be uploaded with genes for rows, separate user gene sets as columns, and 0/1 indicator values.

*For the knowledge-guided mode of Gene Set Characterization, the indicator values of the spreadsheet matrix can be substituted with non-negative “importance” values of the gene’s membership to the gene set. For example, the absolute value of the differential expression (DE) fold change or the negative log of DE significance p-value can be used as importance scores*.

More information about the formatting of input files can be found at our data preparation resource [https://github.com/KnowEnG/quickstart-demos/blob/master/pipeline_readmes/README-DataPrep.md ].

#### User Parameters

There are a number of parameters that the user must select to run the Gene Set Characterization Pipeline.

##### Global Parameters

All modes of Gene Set Characterization require the user to select the

- [species] - for the user gene sets out of the twenty species in the Knowledge Network
- [public_collection] - gene sets to compare their user submitted gene sets to. The public collections available in KnowEnG, for example, include Gene Ontology, Protein Domain Family annotations, GEO expression gene sets. The platform allows the users to select any number of these collections for simultaneous analysis.

##### Knowledge-Guided Only Parameters

In the prior knowledge guided mode, the user must specify the

- [interaction_network] - the gene-gene network available in the Knowledge Network for the selected species to use to create connections between the annotated genes
- [network_percentage] - the extent of influence the interaction network has in contributing to the ranking of annotation gene sets.

#### Data Preprocessing

Once the user selects the Gene Set Characterization inputs and parameters, a simple preprocessing step occurs before the main algorithm. If there are any missing or non-positive values, then the pipeline halts and produces a failure message. The input gene names and identifiers are first mapped to stable Ensembl identifiers of the appropriate [species] using the KN_Mapper tool (see Supp. Note SN5) and the Redis database of gene aliases that accompanies the current Knowledge Network build. Unmapped rows (either missing or ambiguous mappings) are dropped along with the rows that contain duplicated mapped gene identifier. If any negative values are present, the pipeline will return an error message. If the user provided their input gene set as a pasted gene list, then the platform converts this list into a single column spreadsheet format. This spreadsheet will have one row for each Ensembl protein coding gene with a 1 for each gene that is in the user pasted gene list and a zero elsewhere.

#### Description of Algorithm

##### Standard Gene Set Characterization

The KnowEnG Platform provides users with the capability to run standard statistical enrichment of their submitted gene set(s) with the many annotation gene sets of the [public_collection]. This statistical test is performed with the one-sided Fisher's exact test [https://docs.scipy.org/doc/scipy-0.14.0/reference/generated/scipy.stats.fisher_exact.html at its core. For each user gene set and public collection gene set, the test is performed and the significance of the overlap p-value is returned. The gene universe for the test is defined as the intersection of the user's gene universe and the specific public collection's gene universe.

##### Knowledge-Guided Clustering

The knowledge-guided mode of the Gene Set Characterization pipeline implements the DRaWR algorithm in (Blatti and Sinha, 2016) and a more detailed description can be found there. The first step of this approach is to combine the [public_collection] gene annotations with the gene-gene [interaction_network] to construct a single heterogeneous network with both gene and annotation nodes and annotation-gene edges from the [public_collection] data and gene-gene edges from the [interaction_network]. The weights on these edges are then normalized for each of the two different edge types separately. A "baseline" random walk with restart (described in more detail in Supp. Methods SM3) is performed with all genes providing the restart node set and the [network percentage] providing the contribution of the heterogeneous network edges (Supp. Fig. SM11.SF1). For each user gene set "query", a second random walk is performed using only the gene nodes of that gene set as the RWR restart set. If "importance" scores were provided in the submitted spreadsheet, the probability of returning to a gene of the restart set is proportional to its normalized importance score. Finally, the difference between the converged node state vector value of the "query" RWR and the "baseline" RWR is calculated, and the annotation nodes with the greatest difference are returned. These high-ranking annotation nodes are, according to the RWR guilt-by-association process, unusually well connected to the user gene set relative to their overall connection to genes in the network.

#### Pipeline Outputs

##### KnowEnG Platform Interface

Running the KnowEnG Gene Set Characterization (GSC) Pipeline in the KnowEnG Platform will produce results that can be viewed interactively. The GSC visualization displays the association score between user gene sets and the public gene sets calculated selected during pipeline run. This score is displayed in a heatmap view that can be manipulated using various sorting functions to help users visually group the strongest associations. The tool also allows users to drill into each cell for information about the degree of overlap and size and function of the public set. A hierarchical list of collection membership is also provided. Here the user can drill down and easily explore membership, as well as filter the heatmap view by toggling the display of the public gene sets individually or by category.

Downloadable Files

The primary downloadable file of the Gene Set Characterization pipeline is the ranking of [public_collection] annotation terms for their relatedness to the user-submitted gene set(s) along with the corresponding scores from the appropriate analysis mode. *In knowledge-guided mode, additional files containing metadata about the [interaction_network], data preprocessing, and pipeline run are also provided.* More information about the outputs of the pipeline and their structure can be found at [https://github.com/KnowEnG/quickstart-demos/blob/master/pipeline_readmes/README-GSC.md].

#### Methods

For each of the four top-100 gene lists that were derived from the subtype annotation of the ESCC samples (see Supp. Methods SM12, Supp. Table SM10.ST4), we performed the standard and the knowledge-guided mode of the Gene Set Characterization Workflow in the SB-CGC [https://cgc.sbgenomics.com/public/apps#workflow/mepstein/knoweng-genesetcharacterization-public/gene-set-characterization ]. The input gene set spreadsheet for this step is the top_gene_per_phenotype_matrix.txt from the previous workflow task runs. Both runs were done using the taxonomy identifier for humans (9606) with the 'enrichr_pathway' public gene set collection from the Knowledge Network which is from the Enrichr (Chen, et al., 2013) gene set resource and their extracted pathways from WikiPathways (Slenter, et al., 2018) and NCI Pathway Interaction Database (Schaefer, et al., 2009) [https://s3.amazonaws.com/KnowNets/KN-20rep-1706/userKN-20rep-1706/Property/9606/enrichr_pathway/9606.enrichr_pathway.edge ]. This collection had 30,214 gene annotations for 646 pathways, 437 from WikiPathways and 209 from NCI. For the knowledge-guided mode run, we selected the 469,784 edge, 15,999 node, HumanNet (Lee, et al., 2011) Integrated gene-gene [interaction_network] [https://s3.amazonaws.com/KnowNets/KN-20rep-1706/userKN-20rep-1706/Gene/9606/hn_IntNet/9606.hn_IntNet.edge ] and a [network_percentage] of 50%. Each workflow run took under 5 minutes to run and cost less than five cents using spot instances.

The gsc_results.txt results of these two runs were downloaded from the SB-CGC and compiled to compare the top ranking annotation genes sets for each tumor subtype between the standard, 'Fisher', and the knowledge-guided, 'DRaWR' mode of the Gene Set Characterization workflow. Supp. Table SM11.ST1 shows the top 10 pathway annotations for each subtype for both modes. We found that the two different modes frequently agree in their top 10 returned annotation gene sets. Thirteen pathway-subtype associations ranked in the top 10 results for both modes (Supp. Table SM11.ST2). Six of these 15 were supported by existing literature relating the pathways to squamous cell cancer studies and outcomes (Peng, et al., 2017; Schmelzle, et al., 2011; Szumilo, et al., 2009; Zhang, et al., 2018). We were also interested in examining the pathway annotations that only were discovered by the knowledge-guided method. Twelve pathway-subtype associations were in the top 10 for 'DRaWR' mode and not in the top 25 for 'Fisher' mode (Supp. Table SM11.ST3). Among these, seven associations were supported by literature that related the pathway to esophageal or squamous cell cancers (Casazza, et al., 2010; Chen, et al., 2012; Ebihara, et al., 2004; Kiyosue, et al., 2013; Liao, et al., 2011; Song, et al., 2018; Wang, et al., 2011). Five pathway-subtype associations were only in the top 10 for the 'Fisher' mode and not in the top 25 for the knowledge-guided run.

#### Resources

**Gene Set Characterization Pipeline**
KnowEnG Platform Tool
[https://platform.knoweng.org/static/#/pipelines/gene_set_characterization ]
Quickstart Guide [https://knoweng.org/wp-content/uploads/2017/08/GSC_Quickstart.pdf ]
YouTube Tutorial [https://www.youtube.com/watch?v=nP4wtVZOY3E ]
Data Preparation Guidelines [https://github.com/KnowEnG/quickstart-demos/blob/master/pipeline_readmes/README-DataPrep.md]
Downloadable Results Description [https://github.com/KnowEnG/quickstart-demos/blob/master/pipeline_readmes/README-GSC.md ]

**Seven Bridges Cancer Genomics Cloud**
Public Tool [https://cgc.sbgenomics.com/public/apps/#mepstein/knoweng-genesetcharacterization-public/]
Quickstart Guide [https://knoweng.org/wp-content/uploads/2017/12/GSC_CGC_Quickstart.pdf ]
Combined Workflow Tutorial
[https://github.com/KnowEnG/KnowEnG_CWL/tree/master/CGC#running-the-signature-analysis-workflow ]

**Docker and GitHub Repositories**
Gene Set Characterization Docker
[https://hub.docker.com/r/knowengdev/geneset_characterization_pipeline/ ]
Gene Set Characterization GitHub
[https://github.com/KnowEnG/GeneSet_Characterization_Pipeline ]
Data Cleanup Docker [https://hub.docker.com/r/knowengdev/data_cleanup_pipeline/ ]
Data Cleanup GitHub [https://github.com/KnowEnG/Data_Cleanup_Pipeline ]
Pipeline Utilities Docker [https://hub.docker.com/r/knowengdev/base_image/ ]
Pipeline Utilities GitHub [https://github.com/KnowEnG/KnowEnG_Pipelines_Library ]

### Supplementary Methods Tables

**Supplementary Table SM1.ST1. Species in Knowledge Network.**
For each species, lists the taxon identifier for the species as well as the number of Knowledge Network gene nodes, total number edges, and different data source files related to that species.

**Supplementary Table SM1.ST2. External Public Data Repositories.**
For each data source, lists the number of Knowledge Network property/annotation nodes and total number edges derived from that sources as well as the PubMed identifier.

**Supplementary Table SM2.ST1. TCGA Datasets.**
For each omics data type (rows), shows the acronym used for that data type, the number of omics features measured, the number of samples profiled, and the number of clusters produced from the clustering of the data in the original paper.

**Supplementary Table SM2.ST2. TCGA Mutation Data Samples.**
For each primary disease type (rows), shows the acronym used for that disease and the number of samples from a specific “sample location type” (columns).

**Supplementary Table SM3.ST1. PANCAN12 Mutation Sample Clusterings.**
Shows statistics for each clustering of the mutation samples. The first three columns indicate the tool used to perform the clustering as well as additional details about the parameter settings. The next three columns indicate the number of clusters created as well as the size and percentage of the largest cluster. Finally, the last column shows the Kaplan-Meier p-value of the significance of the relationship between the clustering and survival outcome. Separated into sections by A) standard Sample Clustering on mutation data, B) knowledge-guided Sample Clustering on mutation data, C) original TCGA clusterings of samples, D) standard Clustering on alternative data types, and E) COCA analysis with Sample Clustering Pipeline.

**Supplementary Table SM3.ST2. PANCAN12 Mutation Samples Cluster Assignments.**
For each PANCAN12 Mutation Sample in the omics spreadsheet (row), shows the cluster assignment for the six selected clusterings: “sc_noNet” - best hierarchical clustering, “sc_hnInt” and “sc_sText” - network-guided clusterings with HumanNet Integrated and STRING TextMining networks respectively, “disease” - grouping by TCGA primary disease, and “tcga_mut” and “tcga_coca”, mutation-only and COCA clustering analysis in original TCGA paper.

**Supplementary Table SM3.ST3. Enrichment between Clusters.**
For a cluster from our best network-guided method clusterings, “sc_hnInt”, and from one of three original TCGA clusterings, shows the negative log10 p-value of the one-sided Fisher’s exact test for enrichment. The original clusterings are “disease” - grouping by TCGA primary disease, and “tcga_mut” and “tcga_coca”, mutation-only and COCA clustering analysis in TCGA PANCAN12 paper.

**Supplementary Table SM4.ST1. Combined Multi-omics Clustering Matrix.**
A matrix with 0/1 membership indicator values for each PANCAN12 sample (columns) and each different cluster from the original tcat.miRNA clustering, four standard KnowEnG Sample Clusterings of other data types, and one “sc_hnInt” knowledge-guided Sample Clustering on the mutation data.

**Supplementary Table SM7.ST1. Top 100 genes without using the Knowledge Network.**
This table provides a ranked list of top 100 genes identified for each cancer type using the standard mode of operation of Gene/Feature Prioritization pipeline (without the use of Knowledge Network).

**Supplementary Table SM7.ST2. Top 100 genes using HumanNet Integrated Network.**
This table provides a ranked list of top 100 genes identified for each cancer type using the knowledge-guided mode of operation of Gene Prioritization pipeline using the HumanNet Integrated Network.

**Supplementary Table SM7.ST3. Top 100 genes using STRING Text Mining.**
This table provides a ranked list of top 100 genes identified for each cancer type using the knowledge-guided mode of operation of Gene Prioritization pipeline using the STRING Text Mining Network.

**Supplementary Table SM7.ST4. Cross-dataset Intersection.**
This table provides the number of top 100 genes that are shared between the results of different methods for each cancer type.

**Supplementary Table SM7.ST5. Intersection for Prioritization Results without using the Knowledge Network.**
This table provides the number of top 100 genes that are shared between different cancer types, where the top genes are identified using the standard (noNet) mode of Feature Prioritization.

**Supplementary Table SM7.ST6. Intersection for Prioritization Results using HumanNet Integrated.**
This table provides the number of top 100 genes that are shared between different cancer types, where the top genes are identified using the knowledge-guided mode of operation of the Gene Prioritization pipeline using the HumanNet Integrated network.

**Supplementary Table SM7.ST7. Intersection for Prioritization Results using STRING Text Mining.**
This table provides the number of top 100 genes that are shared between different cancer types, where the top genes are identified using the knowledge-guided mode of operation of the Gene Prioritization pipeline using the STRING Text Mining network.

**Supplementary Table SM8.ST1. List of IntOGen driver genes.**
This table provides the list of all the driver genes in IntOGen.

**Supplementary Table SM8.ST2. List of COSMIC driver genes.**
This table provides the list of all the driver genes in COSMIC.

**Supplementary Table SM8.ST3. List of cancer-specific IntOGen driver genes.**
This table shows the list of driver genes for each cancer type present in the PANCAN12 dataset.

**Supplementary Table SM8.ST4. Most significantly enriched GO terms using hnInt.**
This table shows the top 10 GO terms and their enrichment p-values for each cancer type present in PANCAN12 dataset. The enrichment analysis was performed using the standard mode of KnowEnG's Gene Set Characterization pipeline (using Fisher's exact test). The gene sets for each cancer type contained the top 100 genes identified using knowledge-guided gene prioritization with hnInt.

**Supplementary Table SM8.ST5. Most significantly enriched GO terms using sText.**
This table shows the top 10 GO terms and their enrichment p-values for each cancer type present in PANCAN12 dataset. The enrichment analysis was performed using the standard mode of KnowEnG's Gene Set Characterization pipeline (using Fisher's exact test). The gene sets for each cancer type contained the top 100 genes identified using knowledge-guided gene prioritization with sText.

**Supplementary Table SM8.ST6. Most significantly enriched GO terms using noNet.**
This table shows the top 10 GO terms and their enrichment p-values for each cancer type present in PANCAN12 dataset. The enrichment analysis was performed using the standard mode of KnowEnG's Gene Set Characterization pipeline (using Fisher's exact test). The gene sets for each cancer type contained the top 100 genes identified using standard Gene Prioritization (noNet).

**Supplementary Table SM8.ST7. P-value of the most significantly enriched GO terms.**
This table shows the top p-value associated with the most enriched GO term for each cancer type and each prioritization method.

**Supplementary Table SM9.ST1. SB-CGC Datasets.**
Shows the phenotypic information about the RNA-seq samples extracted from the SB-CGC TCGA dataset for ESCA and LUSC cancer types. Three columns were added to the left side of the table as an indicator of which samples relate to which cancer type. The first column shows which samples were mapped to ESCC subgroups in the original analysis.

**Supplementary Table SM10.ST1. Signature Analysis of ESCA and LUSC Samples.**
The TCGA samples from ESCA were divided into four "Sample Groups", three groups of ESCC samples defined in (Cancer Genome Atlas Research, et al., 2017), and one other for all

remaining samples. We calculated the average non-negative correlation of these samples to each LUSC subtype signature and counted the number of samples with a best match to each subtype.

**Supplementary Table SM10.ST2. Best Match Signatures of ESCC Samples.**
Filtered output of Signature Analysis, mapping each of 79 ESCC samples to their best match LUSC subtype signature.

**Supplementary Table SM10.ST3. Top100 Gene Sets for Each Tumor Subtype from ESCC Samples.**
Output of Feature (Gene) Prioritization, capturing top-100 gene lists for each tumor subtype based on the ESCC samples of that subtype compared to all other ESCC samples.

**Supplementary Table SM10.ST4. Overlap Between Top100 Gene Sets.**
For the row and column gene set, shows the number of genes that are present in both. All gene sets are 100 genes except the LUSC-sig_genes set which is the 197 mapped genes from the original LUSC signatures.

**Supplementary Table SM11.ST1. Comparison of Results on ESCC Subtype Gene Sets.**
Shows the ranking of pathway annotation terms for the four top-100 differentially expressed gene lists for each of the four subtypes from the ESCC tumor samples. Each row represents a “Pathway”, “Tumor Subtype” association identified in the top 10 by either the standard, “Fisher”, mode or knowledge-guided, “DRaWR”, mode of the Gene Set Characterization pipeline. The “Source” and “Size” of the pathway annotation gene set provided, as well as the annotation gene set rank for both methods. The “Fisher_pval” significance of the enrichment of the pathway and tumor subtype gene list is also provided. The “Category” shows whether the association is in the top 10 for only one or both modes.

**Supplementary Table SM11.ST2. Shared Results on ESCC Subtype Gene Sets.**
Shows the ranking of pathway annotation terms for the four top-100 differentially expressed gene lists for each of the four subtypes from the ESCC tumor samples. Each row represents a “Pathway”-“Tumor Subtype” association identified in the top 10 by both the standard, “Fisher”, mode or knowledge-guided, “DRaWR”, mode of the Gene Set Characterization pipeline. The “Source” and “Size” of the pathway annotation gene set provided, as well as the annotation gene set rank for both methods. The “Fisher_pval” significance of the enrichment of the pathway and tumor subtype gene list is also provided.

**Supplementary Table SM11.ST3. DRaWR Specific Results on ESCC Subtype Gene Sets.**
Shows the ranking of pathway annotation terms for the four top-100 differentially expressed gene lists for each of the four subtypes from the ESCC tumor samples. Each row represents a “Pathway”-“Tumor Subtype” association identified in the top 10 by the knowledge-guided, “DRaWR”, mode and not the standard, “Fisher”, mode of the Gene Set Characterization pipeline. The “Source” and “Size” of the pathway annotation gene set provided, as well as the

annotation gene set rank for both methods. The “Fisher_pval” significance of the enrichment of the pathway and tumor subtype gene list is also provided.

### Supplementary Methods References

Ashburner, M., *et al.* Gene ontology: tool for the unification of biology. The Gene Ontology Consortium. *Nat Genet* 2000;25(1):25-29.

Blatti, C. and Sinha, S. Characterizing gene sets using discriminative random walks with restart on heterogeneous biological networks. *Bioinformatics* 2016;32(14):2167-2175.

Cai, D., *et al.* Non-negative matrix factorization on manifold. In, *Data Mining, 2008. ICDM'08. Eighth IEEE International Conference on*. IEEE; 2008. p. 63-72.

Cancer Genome Atlas Research, N., *et al.* Integrated genomic characterization of oesophageal carcinoma. *Nature* 2017;541(7636):169-175.

Casazza, A., *et al.* Sema3E-Plexin D1 signaling drives human cancer cell invasiveness and metastatic spreading in mice. *J Clin Invest* 2010;120(8):2684-2698.

Chen, D., *et al.* Increased IL-17-producing CD4(+) T cells in patients with esophageal cancer. *Cell Immunol* 2012;272(2):166-174.

Chen, E.Y., *et al.* Enrichr: interactive and collaborative HTML5 gene list enrichment analysis tool. *BMC Bioinformatics* 2013;14:128.

Choobdar, S., *et al.* Open Community Challenge Reveals Molecular Network Modules with Key Roles in Diseases. *bioRxiv* 2018:265553.

Curtis, C., *et al.* The genomic and transcriptomic architecture of 2,000 breast tumours reveals novel subgroups. *Nature* 2012;486(7403):346-352.

Ebihara, Y., *et al.* Over-expression of E2F-1 in esophageal squamous cell carcinoma correlates with tumor progression. *Dis Esophagus* 2004;17(2):150-154.

Emad, A., *et al.* Knowledge-guided gene prioritization reveals new insights into the mechanisms of chemoresistance. *Genome Biol* 2017;18(1):153.

Emad, A., *et al.* An epithelial-mesenchymal-amoeboid transition gene signature reveals molecular subtypes of breast cancer progression and metastasis. *bioRxiv* 2017:219410.

Forbes, S.A., *et al.* COSMIC: somatic cancer genetics at high-resolution. *Nucleic Acids Res* 2017;45(D1):D777-D783.

Goldman, M., *et al.* The UCSC Cancer Genomics Browser: update 2015. *Nucleic Acids Res* 2015;43(Database issue):D812-817.

Hoadley, K.A., *et al.* Multiplatform analysis of 12 cancer types reveals molecular classification within and across tissues of origin. *Cell* 2014;158(4):929-944.

Hofree, M., *et al.* Network-based stratification of tumor mutations. *Nat Methods* 2013;10(11):1108-1115.

Kiyosue, T., *et al.* Immunohistochemical location of the p75 neurotrophin receptor (p75NTR) in oral leukoplakia and oral squamous cell carcinoma. *Int J Clin Oncol* 2013;18(1):154-163.

Lee, D.D. and Seung, H.S. Learning the parts of objects by non-negative matrix factorization. *Nature* 1999;401(6755):788.

Lee, I., *et al.* Prioritizing candidate disease genes by network-based boosting of genome-wide association data. *Genome Res* 2011;21(7):1109-1121.

Li, Y. and Ngom, A. A new Kernel non-negative matrix factorization and its application in microarray data analysis. *CIBCB* 2012;371:378.

Liao, Y.M., Kim, C. and Yen, Y. Mammalian target of rapamycin and head and neck squamous cell carcinoma. *Head Neck Oncol* 2011;3:22.

Monti, S., *et al.* Consensus clustering: a resampling-based method for class discovery and visualization of gene expression microarray data. *Machine learning* 2003;52(1-2):91-118.

Page, L., *et al.* The PageRank citation ranking: Bringing order to the web. In.: Stanford InfoLab; 1999.

Peng, L., *et al.* Inhibition of glutathione metabolism attenuates esophageal cancer progression. *Exp Mol Med* 2017;49(4):e318.

Rubio-Perez, C., *et al.* In silico prescription of anticancer drugs to cohorts of 28 tumor types reveals targeting opportunities. *Cancer Cell* 2015;27(3):382-396.

Schaefer, C.F., *et al.* PID: the Pathway Interaction Database. *Nucleic Acids Res* 2009;37(Database issue):D674-679.

Schmelzle, M., *et al.* Esophageal cancer proliferation is mediated by cytochrome P450 2C9 (CYP2C9). *Prostaglandins Other Lipid Mediat* 2011;94(1-2):25-33.

Slenter, D.N., *et al.* WikiPathways: a multifaceted pathway database bridging metabolomics to other omics research. *Nucleic Acids Res* 2018;46(D1):D661-D667.

Song, L., Wang, X. and Feng, Z. Overexpression of FOXM1 as a target for malignant progression of esophageal squamous cell carcinoma. *Oncol Lett* 2018;15(4):5910-5914.

Szklarczyk, D., *et al.* STRING v10: protein-protein interaction networks, integrated over the tree of life. *Nucleic Acids Res* 2015;43(Database issue):D447-452.

Szumilo, J., *et al.* Expression of syndecan-1 and cathepsins D and K in advanced esophageal squamous cell carcinoma. *Folia Histochem Cytobiol* 2009;47(4):571-578.

Taube, J.H., *et al.* Core epithelial-to-mesenchymal transition interactome gene-expression signature is associated with claudin-low and metaplastic breast cancer subtypes. *Proc Natl Acad Sci U S A* 2010;107(35):15449-15454.

Wang, W., *et al.* Enhanced PPAR-gamma expression may correlate with the development of Barrett's esophagus and esophageal adenocarcinoma. *Oncol Res* 2011;19(3-4):141-147.

Wilkerson, M.D., *et al.* Lung squamous cell carcinoma mRNA expression subtypes are reproducible, clinically important, and correspond to normal cell types. *Clin Cancer Res* 2010;16(19):4864-4875.

Zerbino, D.R., *et al.* Ensembl 2018. *Nucleic Acids Res* 2018;46(D1):D754-D761.

Zhang, J., *et al.* Nrf2 and Keap1 abnormalities in esophageal squamous cell carcinoma and association with the effect of chemoradiotherapy. *Thorac Cancer* 2018;9(6):726-735.
