## Supplementary Methods Figures for "Knowledge-guided analysis of ‘omics’ data using the KnowEnG cloud platform"

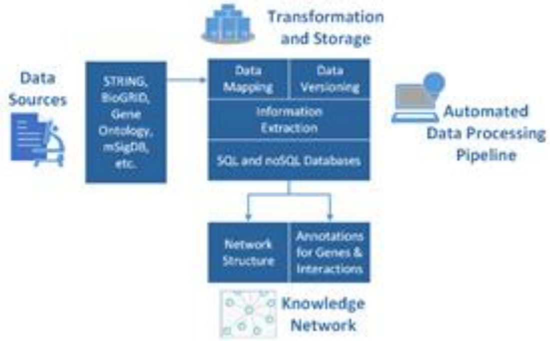

**Supplementary Figure SM1.SF1. Overview of the Knowledge Network Build Pipeline.**
Overall flow of data from original external public repositories to final exported KnowEnG Knowledge Network.

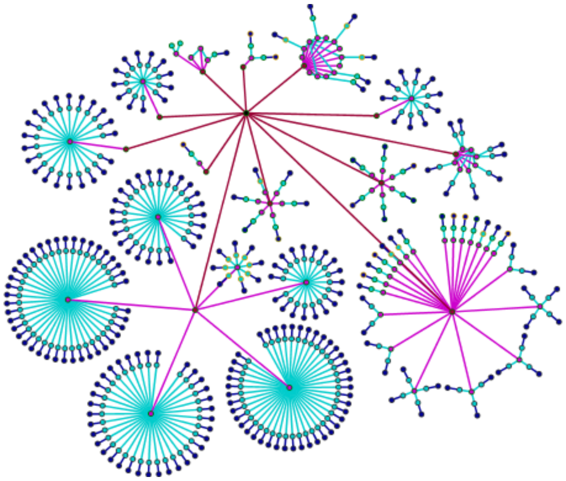

**Supplementary Figure SM1.SF2. Parallelized Task Execution Example.**
Each node in this diagram is a containerized task in the Build Pipeline and the edges represent dependencies between tasks. The node color shows the type of task: check external source (red), download external file (magenta), parse external data format (cyan), and map interaction entities (dark blue). The node borders show an example status of each task: successful (green), running (yellow), or queued (black). For a particular size of the cloud deployment, the diagram shows how tasks are partitioned and can be executed simultaneously.

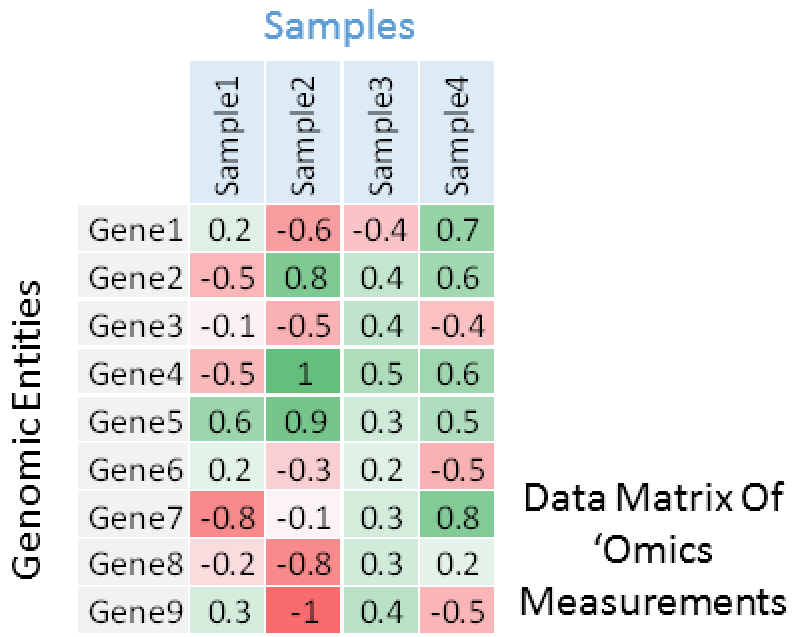

| | Sample1 | Sample2 | Sample3 | Sample4 |
|---|---|---|---|---|
| Gene1 | 0.2 | -0.6 | -0.4 | 0.7 |
| Gene2 | -0.5 | 0.8 | 0.4 | 0.6 |
| Gene3 | -0.1 | -0.5 | 0.4 | -0.4 |
| Gene4 | -0.5 | 1 | 0.5 | 0.6 |
| Gene5 | 0.6 | 0.9 | 0.3 | 0.5 |
| Gene6 | 0.2 | -0.3 | 0.2 | -0.5 |
| Gene7 | -0.8 | -0.1 | 0.3 | 0.8 |
| Gene8 | -0.2 | -0.8 | 0.3 | 0.2 |
| Gene9 | 0.3 | -1 | 0.4 | -0.5 |

**Supplementary Figure SM3.SF1. Sample of "omics" Spreadsheet.**
Users must input genomic spreadsheets were the genes features are the rows and the samples are the columns.

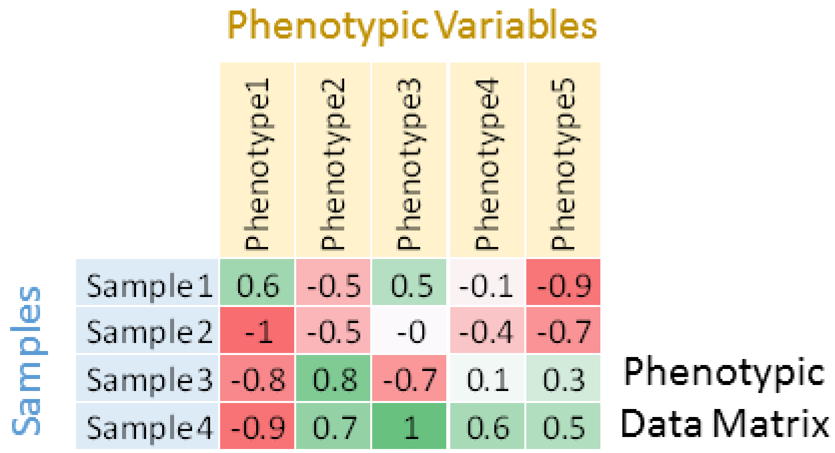

| | Phenotype1 | Phenotype2 | Phenotype3 | Phenotype4 | Phenotype5 |
|---|---|---|---|---|---|
| Sample1 | 0.6 | -0.5 | 0.5 | -0.1 | -0.9 |
| Sample2 | -1 | -0.5 | -0 | -0.4 | -0.7 |
| Sample3 | -0.8 | 0.8 | -0.7 | 0.1 | 0.3 |
| Sample4 | -0.9 | 0.7 | 1 | 0.6 | 0.5 |

**Supplementary Figure SM3.SF2. Sample of Phenotypic Spreadsheet.**
Users must input phenotypic spreadsheets were the samples are the rows and each column represent a distinct phenotype.

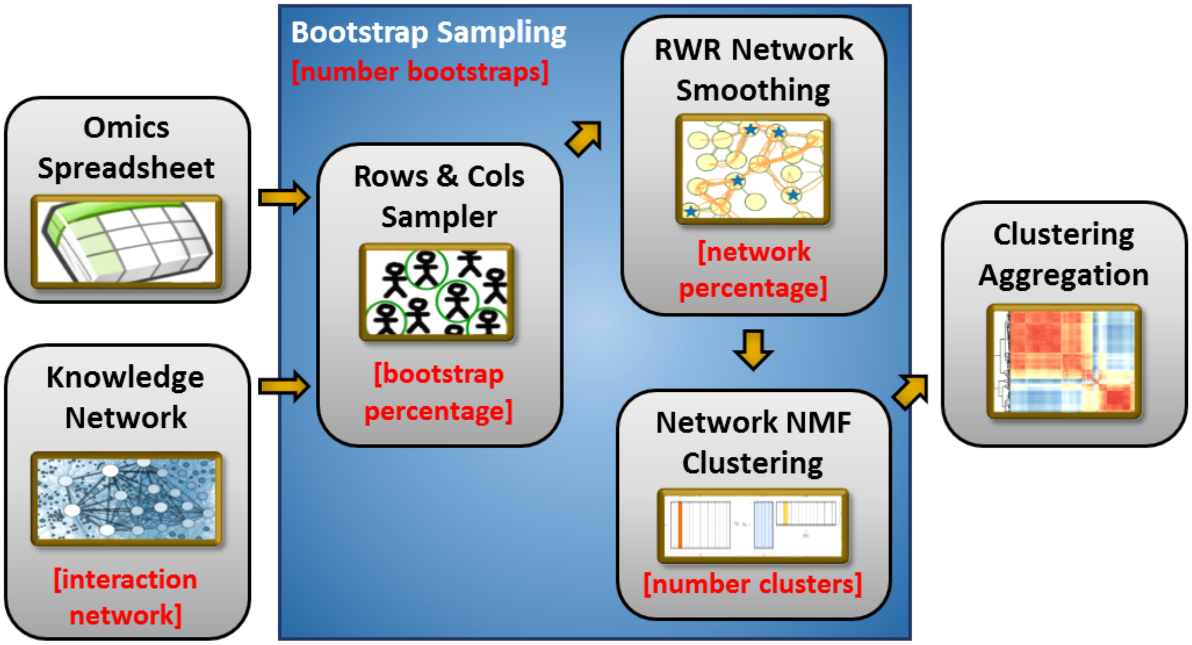

**Supplementary Figure SM3.SF3. Overview of Knowledge-Guided Sample Clustering.**
For each bootstrap, the core clustering algorithm smooths the original features using the knowledge network and then performs NMF clustering. The consensus matrix is built from the bootstraps and finally clustered with K-means. Steps are shown with their corresponding user parameters (in red).

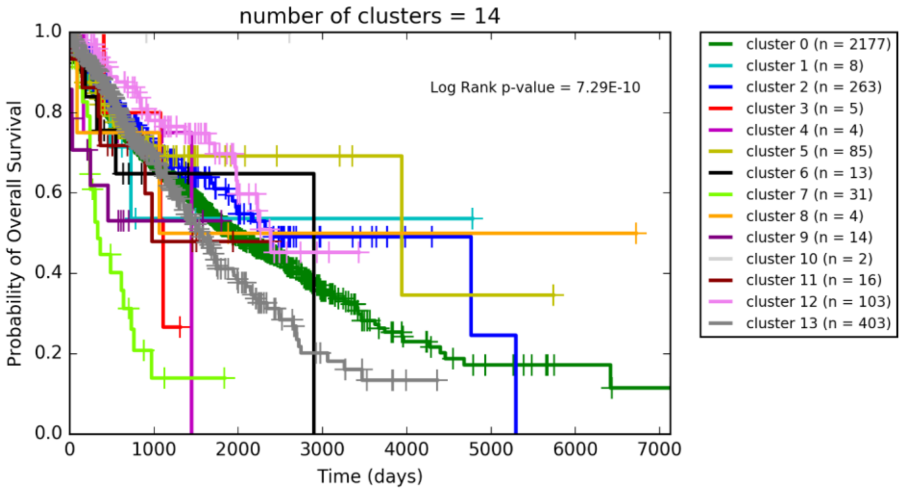

**Supplementary Figure SM3.SF4. Survival Analysis of “sc_noNet” Clustering.**
Each cluster is plotted as a separate survival curve in the Kaplan-Meier plot and the p-value of the multivariate log rank test is reported.

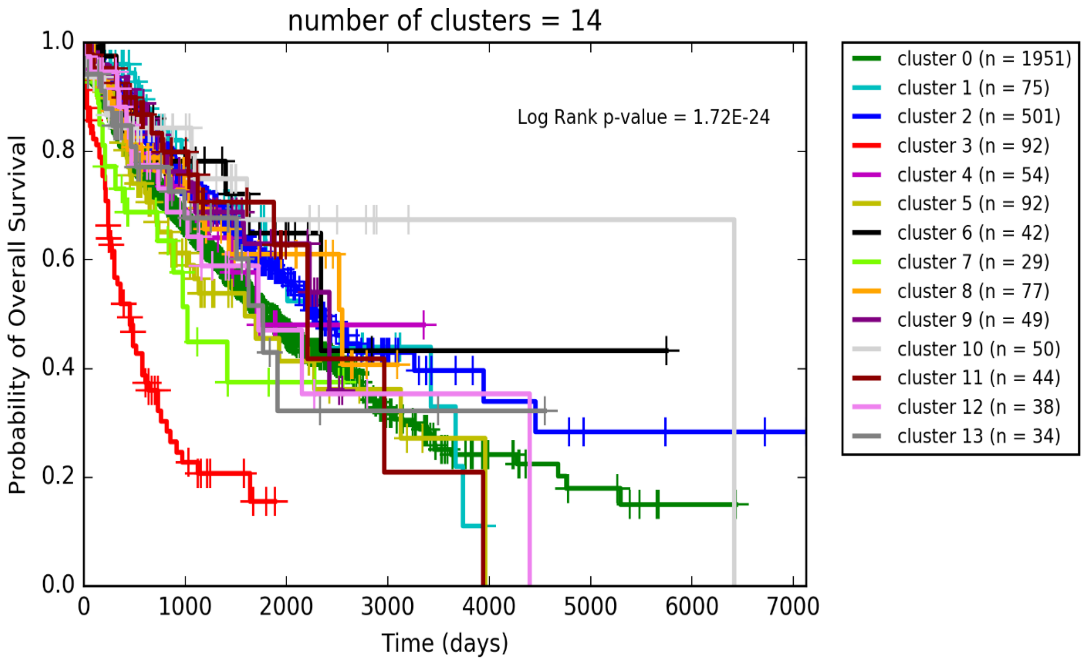

**Supplementary Figure SM3.SF5. Survival Analysis of "sc_sText" Clustering.**
Each cluster is plotted as a separate survival curve in the Kaplan-Meier plot and the p-value of the multivariate log rank test is reported.

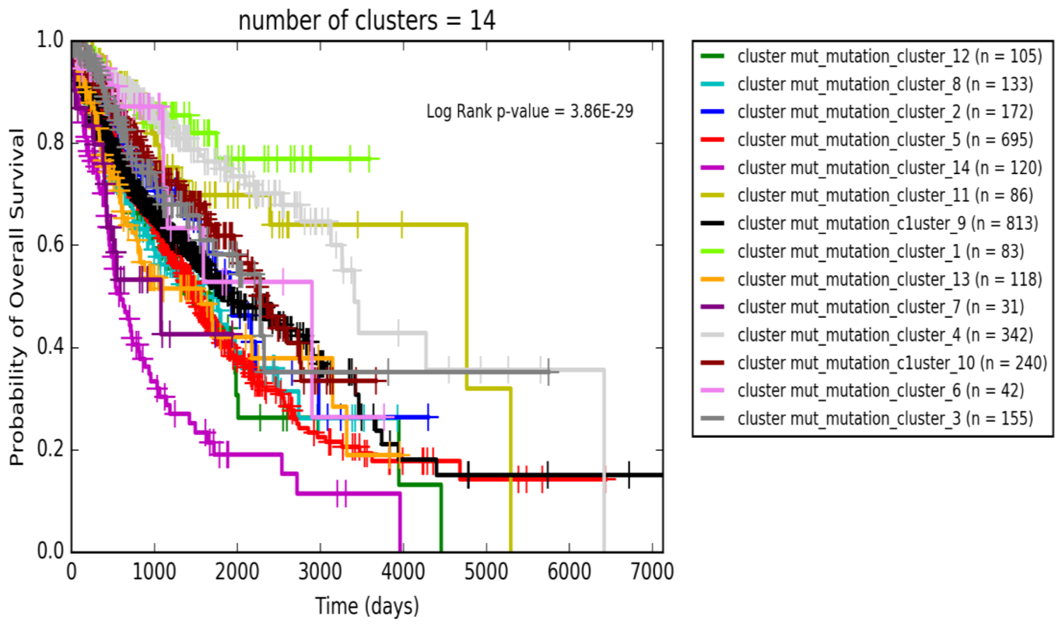

**Supplementary Figure SM3.SF6. Survival Analysis of "tcga_mut" Clustering.**
Each cluster is plotted as a separate survival curve in the Kaplan-Meier plot and the p-value of the multivariate log rank test is reported.

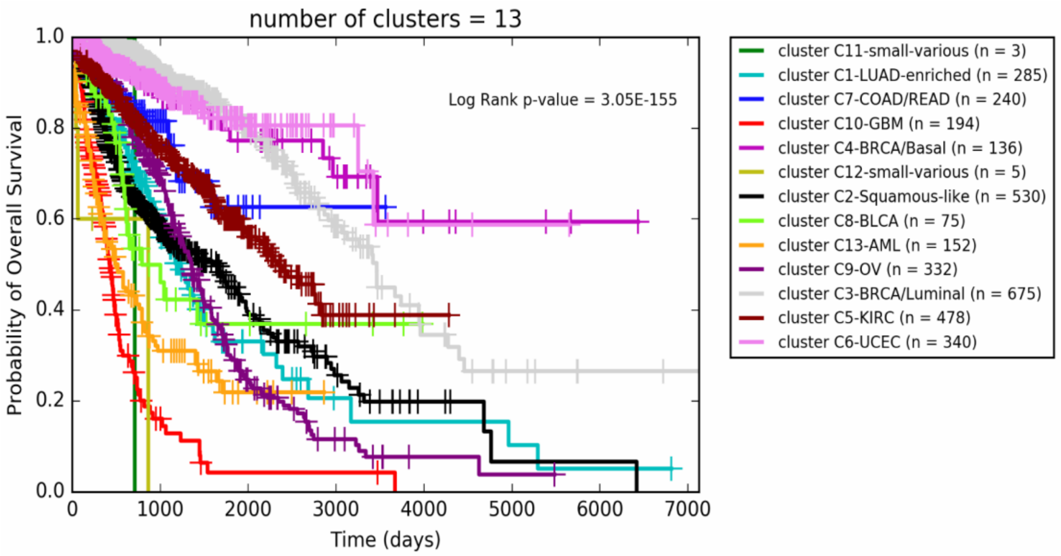

**Supplementary Figure SM5.SF1. Survival Analysis of "tcga_coca" Original COCA .**
Each cluster is plotted as a separate survival curve in the Kaplan-Meier plot and the p-value of the multivariate log rank test is reported.

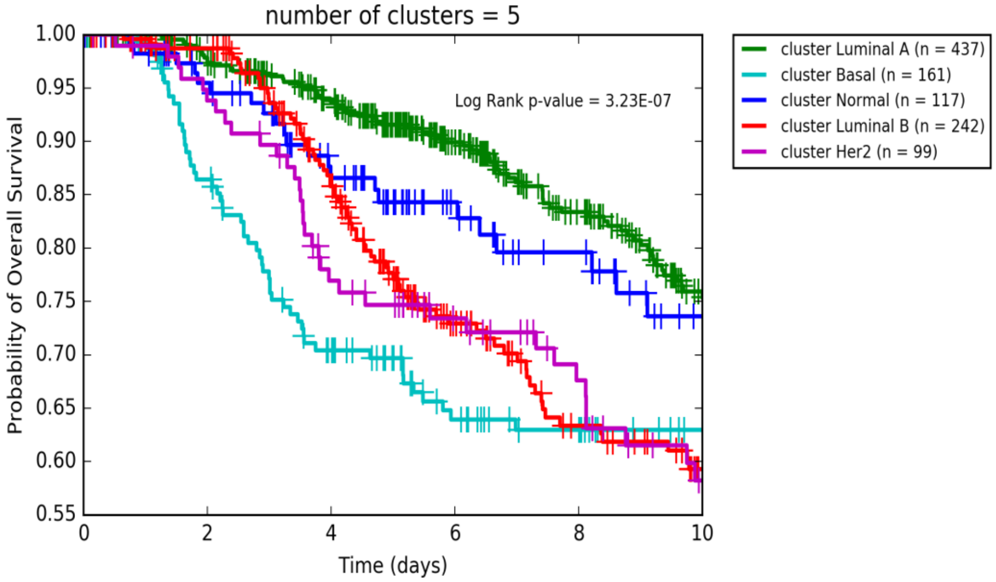

**Supplementary Figure SM6.SF1. Survival probabilities of PAM50 subtypes of METABRIC samples.**
Each cluster is plotted as a separate survival curve in the Kaplan-Meier plot and the p-value of the multivariate log rank test is reported.

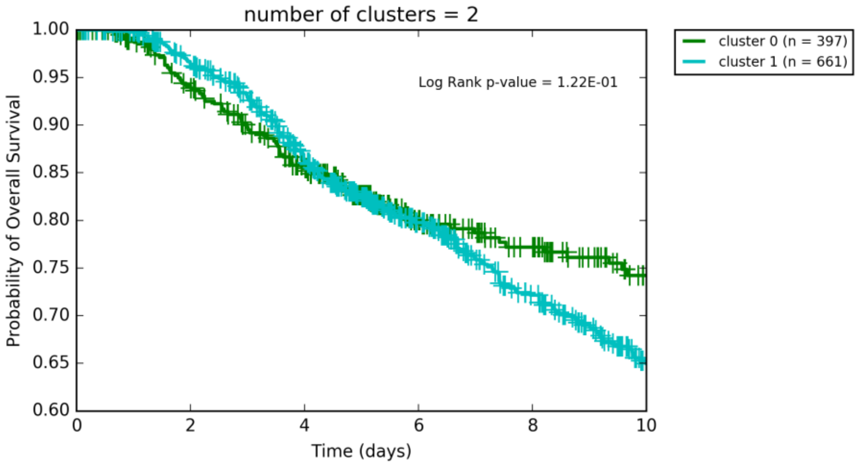

**Supplementary Figure SM6.SF2. Survival Analysis of METABRIC samples using EMT signature and Standard Sample Clustering.**
Each cluster is plotted as a separate survival curve in the Kaplan-Meier plot and the p-value of the log rank test is reported.

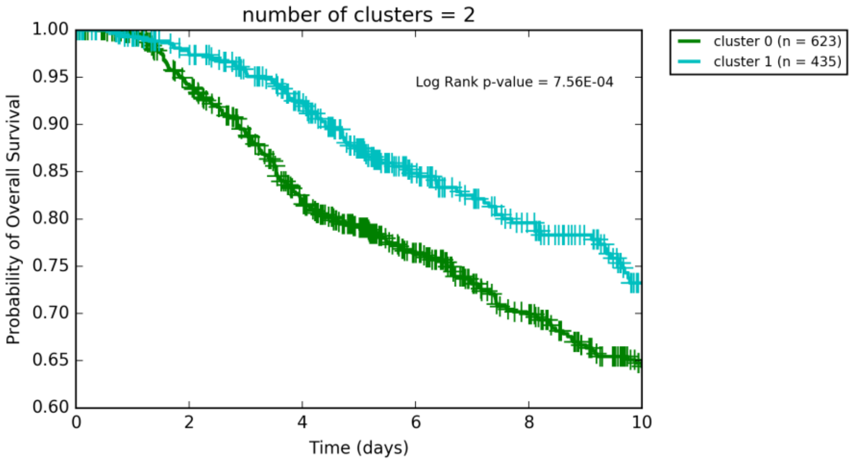

**Supplementary Figure SM6.SF3. Survival Analysis of METABRIC samples using EMT signature and Knowledge-guided Sample Clustering with hnInt.**
Each cluster is plotted as a separate survival curve in the Kaplan-Meier plot and the p-value of the log rank test is reported.

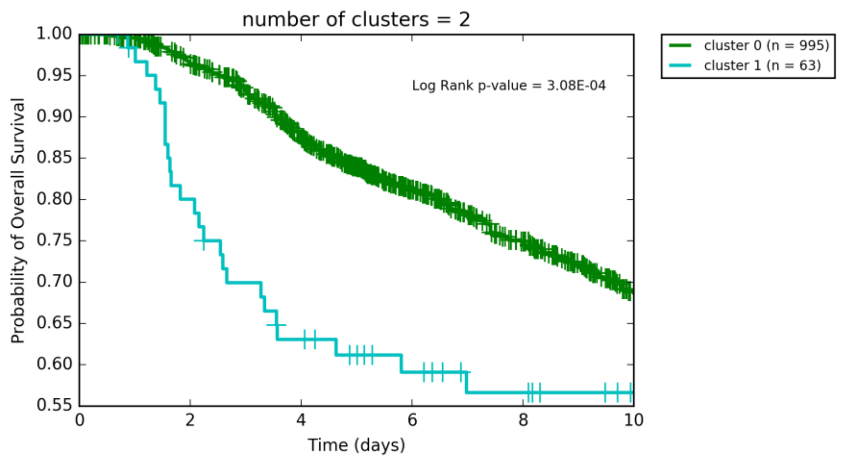

**Supplementary Figure SM6.SF4. Survival Analysis of METABRIC samples using EMT signature and Knowledge-guided Sample Clustering with sText.**

Each cluster is plotted as a separate survival curve in the Kaplan-Meier plot and the p-value of the log rank test is reported.

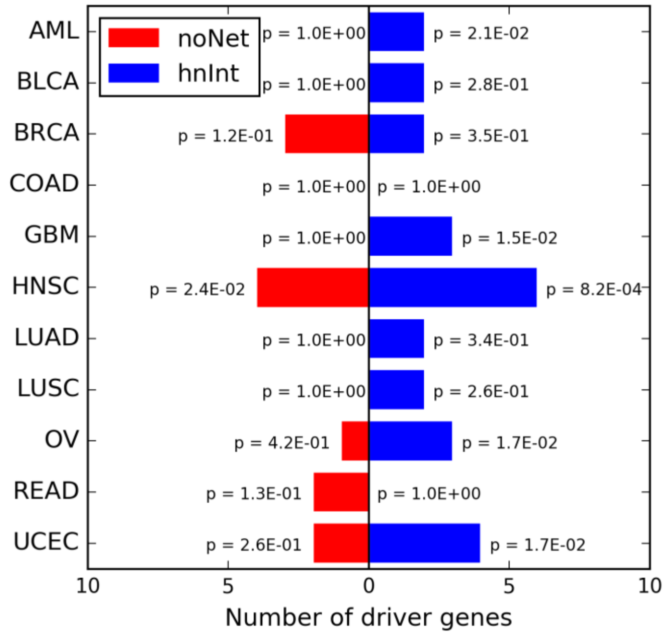

**Supplementary Figure SM8.SF1. IntOGen Cancer-Specific Drivers among Top Genes Identified using noNet and hnInt.**

The bars represent the number of cancer-specific driver genes (from IntOGen) among the top 100 genes identified using noNet (red) and hnInt (blue) for each cancer type. The p-values represent the significance of enrichment, calculated using Fisher's exact test.

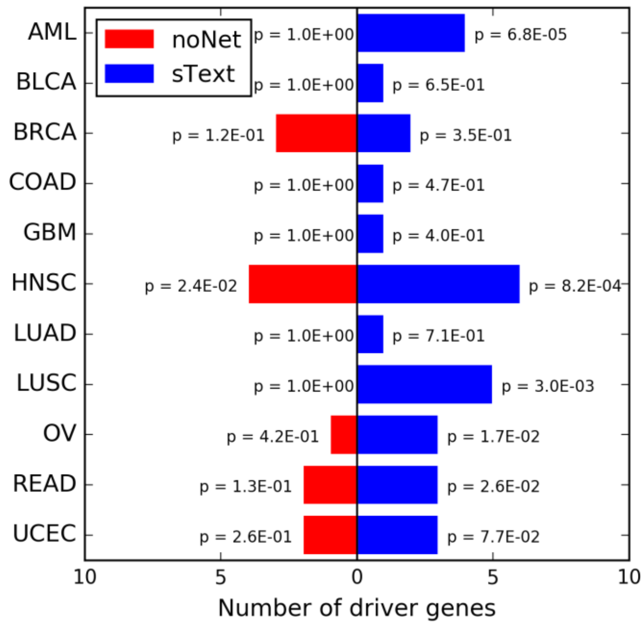

**Supplementary Figure SM8.SF2. IntOGen Cancer-Specific Drivers among Top Genes Identified using noNet and sText.**

The bars represent the number of cancer-specific driver genes (from IntOGen) among the top 100 genes identified using noNet (red) and sText (blue) for each cancer type. The p-values represent the significance of enrichment, calculated using Fisher's exact test.

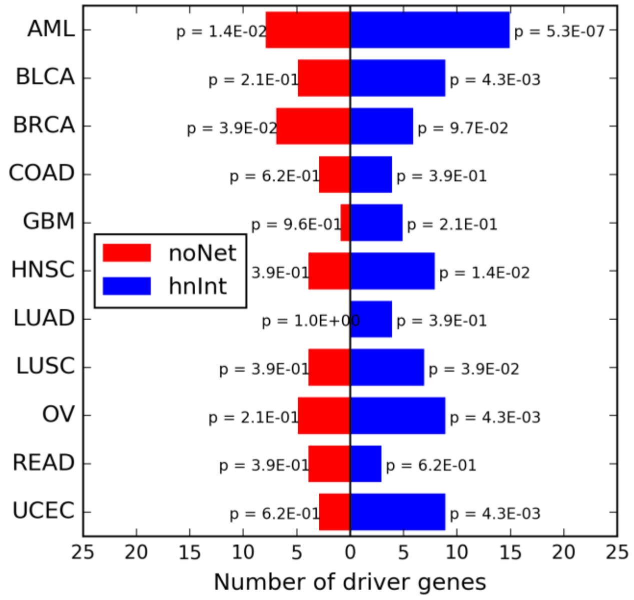

**Supplementary Figure SM8.SF3. IntOGen Driver Genes among Top Genes Identified using noNet and hnInt.**

The bars represent the number of driver genes (from IntOGen) among the top 100 genes identified using noNet (red) and hnInt (blue) for each cancer type. The p-values represent the significance of enrichment, calculated using Fisher's exact test.

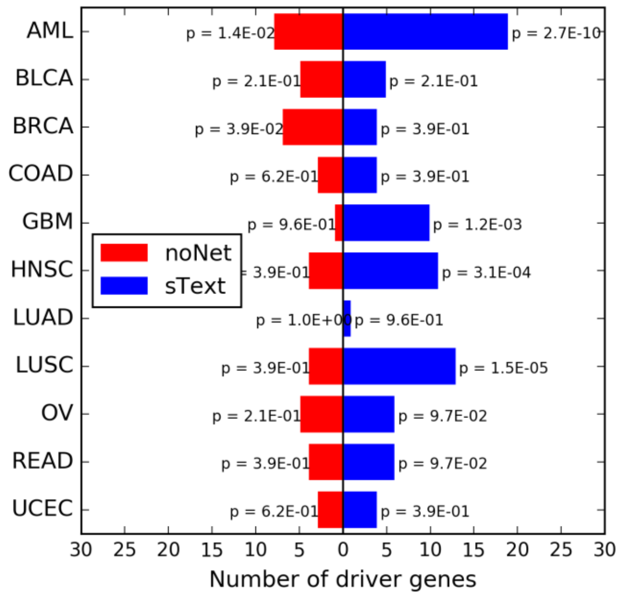

**Supplementary Figure SM8.SF4. IntOGen Driver Genes among Top Genes Identified using noNet and sText.**

The bars represent the number of driver genes (from IntOGen) among the top 100 genes identified using noNet (red) and sText (blue) for each cancer type. The p-values represent the significance of enrichment, calculated using Fisher's exact test.

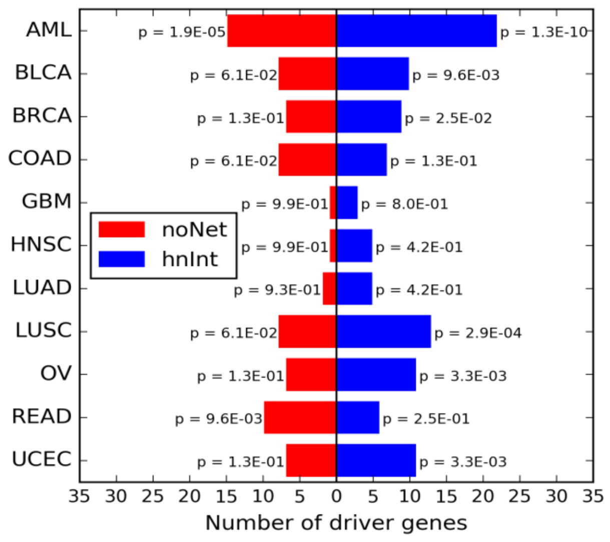

**Supplementary Figure SM8.SF5. COSMIC Driver Genes among Top Genes identified Using noNet and hnInt.**

The bars represent the number of driver genes (from COSMIC) among the top 100 genes identified using noNet (red) and hnInt (blue) for each cancer type. The p-values represent the significance of enrichment, calculated using Fisher's exact test.

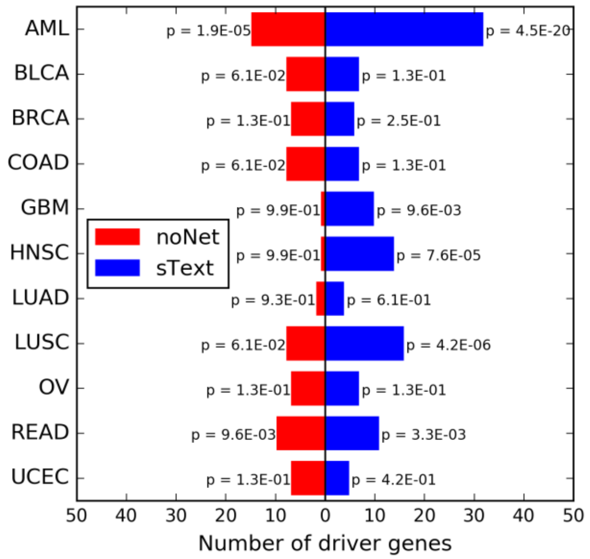

**Supplementary Figure SM8.SF6. COSMIC Driver Genes among Top Genes identified Using noNet and sText.**

The bars represent the number of driver genes (from COSMIC) among the top 100 genes identified using noNet (red) and sText (blue) for each cancer type. The p-values represent the significance of enrichment, calculated using Fisher's exact test.

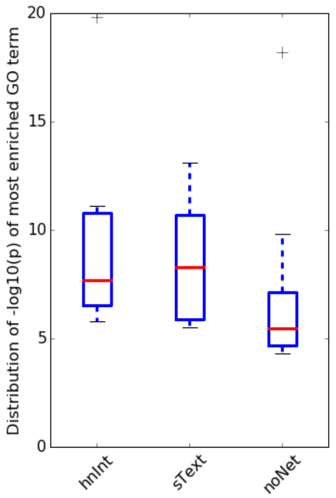

**Supplementary Figure SM8.SF7. Distribution of Most Enriched GO terms for Different Prioritization Methods.**

For each cancer type and each prioritization method, the list of top 100 genes is identified and GO enrichment analysis (using Fisher's exact test) is performed to identify the most significantly enriched GO term. Each box shows the distribution of -log10(p) of these significantly enriched GO terms for different cancer types. The red line represents the median.

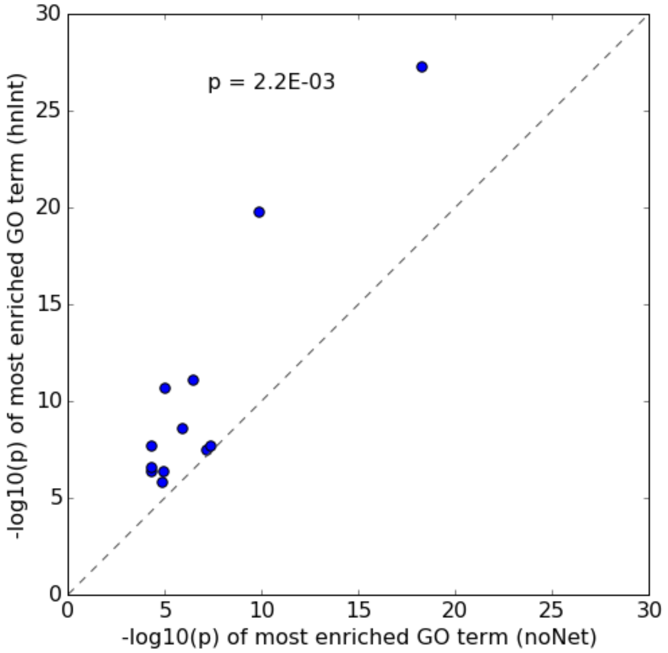

**Supplementary Figure SM8.SF8. Most Enriched GO terms using hnInt versus noNet.** Each circle represents one cancer type. The p-value of improvement is calculated using Wilcoxon signed rank test.

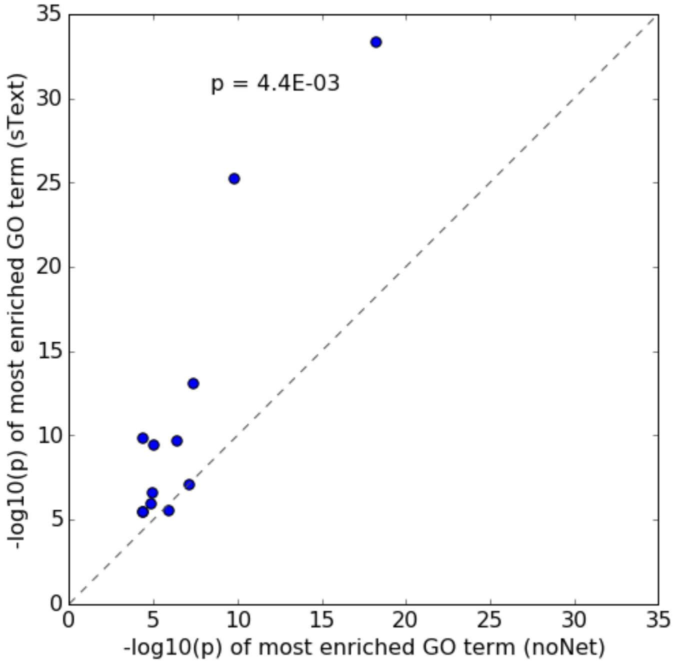

**Supplementary Figure SM8.SF9. Most Enriched GO terms using sText versus noNet.** Each circle represents one cancer type. The p-value of improvement is calculated using Wilcoxon signed rank test.

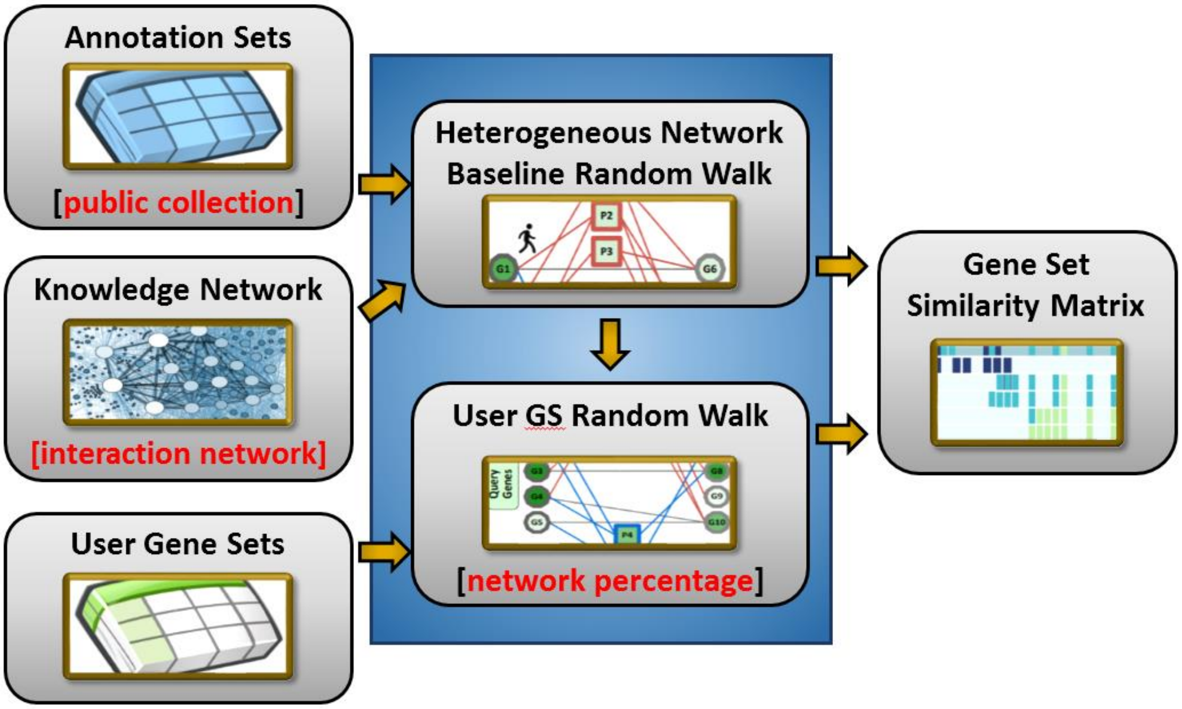

**Supplementary Figure SM11.SF1. Overview of Knowledge-Guided Gene Set Characterization.**

A heterogeneous network is built from the selected gene-gene [interaction_network] and the annotation gene sets of the [public_collection]. A "baseline" random walk is allowed to converge using the [network percentage] and all genes as equal probability restarts. For each submitted user gene set, a random walk that only restarts the the users genes is performed and annotation set nodes are evaluated by their increase from their baseline.
