## Supplementary Notes Document for "Knowledge-guided analysis of ‘omics’ data using the KnowEnG cloud platform"

#### Supplementary Note 1: Infrastructure of the KnowEnG Platform

##### Overview

The KnowEnG platform is a cloud-based web application. A free, public version of the platform is maintained by the Center. Users may also deploy private instances with their own Amazon Web Services (AWS) accounts using a CloudFormation Template [https://aws.amazon.com/cloudformation/aws-cloudformation-templates/], which streamlines setup (Supp. Note SN3). The public and private versions share a common infrastructure, described here.

As shown in Supp. Fig. SN1.SF1, the platform software system consists of containerized components running on a cluster of virtual machines. User-submitted analyses are executed in short-running containers (those labeled Job Worker 1-N in the figure), the same as used in the SB-CGC (Supp. Note SN10) graphical user interfaces. All other containers are long-running, always-on services that implement core functionality such as the API Server and data persistence. Both the short-running and long-running containers are orchestrated by Kubernetes, which automatically adjusts the cluster size, adding and removing nodes, as load changes. This allows the system to remain performant during periods of high demand without incurring costs for excess capacity during periods of low demand.

In the current deployments, AWS provides the cloud computing resources, which include EC2 virtual machines [https://aws.amazon.com/ec2/] as cluster nodes, S3 buckets [https://aws.amazon.com/s3/] for long-term storage, ElastiCache [https://aws.amazon.com/elasticache/] for in-memory caches, and Elastic File System [https://aws.amazon.com/efs/] for shared working storage among cluster nodes. The Center has also deployed the system outside AWS and will endeavor to remain compatible with other cloud providers.

The graphical user interface is single-page application [https://en.wikipedia.org/wiki/Single-page_application] implemented in Angular [https://angular.io/]. Visualizations are rendered using Scalable Vector Graphics [https://www.w3.org/TR/SVG/] and HTML canvas [https://html.spec.whatwg.org/multipage/canvas.html] using data fetched from the Representation State Transfer (REST) [https://www.w3.org/TR/2004/NOTE-ws-arch-20040211/#relwwwrest] API.

The platform adopts best practices and standards-based measures for security. Among these are OAuth 2.0 [https://oauth.net/2/] for single sign-on, JSON Web Tokens [https://jwt.io/] for authentication and authorization, Transport Layer Security [https://tools.ietf.org/html/rfc8446] to encrypt all traffic over the open Internet, and AWS Virtual Private Clouds [https://aws.amazon.com/vpc/] for network security.

#### Cost Analysis

The cloud-computing costs of running the public platform instance can be decomposed into three parts:

1. **Baseline compute**. This is the cost of running the cluster at is smallest size, during periods of low utilization, when it consists of four EC2 nodes. At current AWS on-demand prices as of May 2019, the total baseline compute cost is $0.4904 per hour, or about $364.86 per month. This cost could be reduced by (a) switching from on-demand nodes to Reserved Instances, which require a minimum one-year commitment, and (b) tuning node sizes, possibly reducing the responsiveness of the system during periods of low utilization.
2. **Burst compute**. This is the cost of adding extra nodes to the cluster during periods of high utilization. We currently allow the system to add up to ten m5.4xlarge [https://aws.amazon.com/ec2/instance-types/] nodes, each of which has 16 CPUs and 64 GiB RAM and costs $0.768 per hour.
3. **Storage**. This is the cost of storing cluster data on the Elastic File System. AWS charges $0.30 per gigabyte per month, and total cost to the Center has averaged $32.21 per month for September 2018 through February 2019. This cost could be reduced by (a) migrating more runtime data from Elastic File System to S3 buckets, which are billed at a lower rate, and (b) tightening data-retention policies that govern the persistence of user uploads and analyses.

#### Resources

##### KnowEnG Platform Components

Docker Image for API Server [https://hub.docker.com/r/knowengdev/nest_flask/]
Docker Image for Job Queue [https://hub.docker.com/r/knowengdev/nest_jobs/]
GitHub Repository [https://github.com/KnowEnG/platform]

### Supplementary Note 2: KnowEnG Platform User Experience

#### Overview

KnowEnG is a web-based platform designed to provide biologists with an **intuitive user interface** enabling the execution of complex analytical processes, as well as **interactive visual displays** supporting the study and evaluation of results. The KnowEnG team believed that a carefully considered UI was critical to wide adoption of the platform by people who do not have experience in analytics. Toward that end, we brought together a variety of expertise and perspectives including, biomedical scientists, data scientists, software engineers and designers. Through these collaborations, we identified platform features that (1) addressed the needs of the scientists, (2) harnessed the power of the analytics, and (3) fell within resource and time constraints. Our solutions incorporate information design principles that we have used successfully in other domains (Baker and Bushell, 1995; Tufte, et al., 1998). These include providing (1) qualitative overviews coupled with quantitative details, (2) efficient comparison methods, (3) relevant context and evidence for confirmation, and (4) illustrations of relationships between data types [http://vi-bio.ncsa.illinois.edu/next-gen-sequencing-results-visualization.html].

#### Methods

##### Technology Overview

The KnowEnG Platform interfaces were designed using Sketch App user interface design tool and InVision prototyping and style markup tool. InVision, specifically, provided developers with access to all of the interface mockups and was used during all stages of development to collaborate on design software feature decisions with all members of the development team.

The technology stack includes a single-page web application in the Angular framework, backed by a REST API in Python Flask with Redis, PostgreSQL, and Amazon S3 for data persistence. The compute cluster is managed by Kubernetes and automatically adjusts to user load by adding or removing Amazon EC2 virtual machines. The environment employs best practices for security, including OAuth 2.0 authentication and virtual private clouds. A free, public instance of the platform is administered by KnowEnG staff. Users may easily deploy private instances of the platform to their own AWS accounts using KnowEnG's Cloud Formation Template.

##### Workflow

The KnowEnG platform is geared to the biologist and provides easy access to cutting edge analytical methods required to study genomic data effectively. The platform presents a process that includes easy data upload and selection, effortless algorithm parameter tuning for complex machine learning experiments, and sophisticated visualization tools for evaluating data and studying results. It provides guidance on preparing data, choosing appropriate pipelines, and adjusting parameters. It provides interactive, visual methods for filtering and comparing data. Moreover, as users move through the workflow, contextually relevant help content is available for reference.

##### Navigation

The platform has three conceptual areas: *Analysis Pipelines, Data,* and *Support*. Clicking on *Analysis Pipelines* provides information about each pipeline including a description of the methods, common uses, input requirements, and links to tutorials and video clips. Here, the user can start a pipeline (Supp. Fig. SN2.SF1). Clicking on *Data* provides users with persistent access to their uploaded study data and resulting data produced by the executed pipelines. KnowEnG also provides example data for each pipeline for demonstration. *Support* houses contacts, documentation/training, and links to KnowEnG relevant publications and their associated data.

##### Launching Analysis Pipelines

The KnowEnG platform offers pipelines for common genomics analysis tasks. Current pipelines include Sample Clustering, Feature Prioritization, Gene Set Characterization, Signature Analysis, and Spreadsheet Visualization. Each pipeline provides common algorithmic approaches, and some include novel techniques that incorporate the use of prior knowledge. Selecting “Start Pipeline” for any one of these initiates a step-based setup tool. The setup flow is generally to upload/select data, and then to make parameters choices (Supp. Fig. SN2.SF2). The last step displays all selections for review and allows the user to add notes to keep track of data experiments. The analysis is the launched and runs automatically in the cloud. Results of the pipeline appear in the Data section. Clicking on any completed pipeline run displays reference information about the run and the option to study results in the visualization view or to download a zip file of the results to the local computer. Results of some pipelines can also be used as data inputs to certain other pipelines (Supp. Fig. SN2.SF3).

##### Viewing the Results

All pipelines have interactive visualizations that help the researcher review and study the results. The visuals provide high-level understanding of the data, and multiple features allow the user to filter, sort, and compare. Quantitative details are provided via drill-down methods. Details of each visualization are not discussed in this supplement, but a few selected, novel features are described below:

##### Example Features

<u>Distribution Graphs:</u> Most of the custom visualizations provide descriptive statistics that help users understand the distribution of their data across user-selected variables (often phenotypic). These are available by hovering over or clicking on the elements in the visualization. In the Sample Clustering visualization, the small graph shows the distribution of both continuous and categorical data. Hovering over graph shows the percent of data and quantity of samples at that point (or in the bin). Clicking on the single row heatmap opens a more detailed view of the data distribution, including a breakdown of the distribution based on the “group columns by” variable (Supp. Fig. SN2.SF4). This is recalculated on the fly as the user explores other “group columns by” variables.

<u>Clustering:</u> The visualization for Sample Clustering provides an interactive heatmap displaying the samples (e.g. patients) as columns and the data (e.g. gene expression values) as rows. The columns are grouped by the cluster assignment, and sorted within each grouping by each sample’s silhouette score (the measure of how well the sample fits in that cluster). The rows are sorted by the strength of the statistical association between the row and the clustering assignments. The user can re-sort the rows by variance of data as well. In cases where the researcher included “bootstrapping” in the clustering analysis, a second heatmap is displayed below the other. This heatmap displays the same samples as columns and always stays aligned with the heatmap above. The rows duplicate the same samples providing a matrix that indicates how frequently each pair of samples was assigned to the same cluster, as the algorithm performed extensive clustering operations. This is referred to as the “consensus matrix”.

The single row heatmaps at the top of the screen allow the user to re-group and re-sort the columns by other features of interest, including phenotypic features. Additional single row heatmaps can be displayed at the bottom and their columns are also aligned to the same samples (Supp. Fig. SN2.SF5). This allows for fluid comparison across these different heatmaps on the same screen allowing users to get a sense of the features that are guiding and correlated with the current grouping. For more information about features, see Supp. Note SN8.

Top Features Selector: The Top Features Selector appears in several visualization to help users control the number of features being displayed in the visualization. Rather than arbitrarily selecting top 10 or 100 features, the user can use this tool to select “top feature” thresholds based on seeing the distribution of the scores from highest to lowest. Researchers can see at a glance if there is small set of high scoring features, or if there is a gradual transition between high and low scoring (Supp. Fig. SN2.SF6). This not only allows users to select what is most interesting, but also affords a peek into the distribution of feature scores across all the data.

Filtering and Sorting: The visualizations in KnowEnG incorporate multiple methods for extensive sorting and filtering. This high-level of interactivity supports the efficient study of results. One example is the Gene Set Characterization visualization, which helps researchers see how their genesets overlap with the extensive public geneset collections that make up the Knowledge Network. The results are displayed as a heatmap where the researcher’s gene sets are rows and the public sets are columns. These columns can be grouped by class and sorted alphabetically, or by how well they score across the user gene sets. The cells display the color associated with the score and are clickable to access detailed information. The control panel on the left organizes all the public gene sets and supports refined filtering based on the ontology of each class of gene sets (Supp. Fig. SN2.SF7). Quantitative information about the level of enrichment is integrated in this panel helping the user more quickly find the areas of greatest relevance.

Drill Down: Throughout the platform, multiple drill down methods are provided to gain visibility into the data and help guide investigation. For example, clicking on a cell in the Gene Set Characterization similarity matrix pops up a panel that shows the number and percentage of genes in the user gene set that are also in that public gene set. In the example image, the user gene set, Ovarian Diseases, overlaps with 66.7% of the genes in the selected pathway gene set (Supp. Fig. SN2.SF8).

##### Data Handover between Pipelines

Multiple KnowEnG pipelines can be weaved together into workflows that offer a series of complementary perspectives on the user's data set. For example, a user could begin by partitioning a sample population into subgroups with Sample Clustering, proceed to find the genes most associated with the assigned cluster labels using Feature Prioritization, and conclude by identifying the most significantly enriched Gene Ontology terms with Gene Set Characterization. The KnowEnG platform allows the user to perform a variety of such multi-step analyses entirely inside the application, without having to download intermediate results and transform them for subsequent use. Figure 1D illustrates the supported types of pipeline handover.

To facilitate handover, spreadsheets produced in the execution of a pipeline are automatically saved alongside the user's uploads, grouped with the job that generated them. Later, when configuring another pipeline for execution, the user can simply select one of these output spreadsheets as an input.

##### Other KnowEnG User Experience Options

In addition to the web-based platform, KnowEnG provides two additional user experiences: Seven Bridges Cancer Genomics Cloud (SB-CGC) provides users with a query interface to work on major data sets, such as TCGA, alongside the KnowEnG analytical workflows. The Common Workflow Language is used to describe the sequence of container invocations that constitute the analysis. Such descriptions are then executed by the SB-CGC’s engine, which provisions the necessary computational resources. For more see Supp. Note SN10.
Jupyter Notebooks are a popular tool for data analysis. Accordingly, KnowEnG offers a free, public web server pre-populated with a collection of notebooks that showcase the main analytical pipelines and enable common data manipulations. Users can step through the notebooks as provided, run new analyses by replacing the sample data with their own spreadsheets, and even introduce new functionality by editing the notebook scripts. The web server is built on the open-source JupyterHub project, deployed to AWS via Kubernetes, and offers users the same single-sign on technology as the platform. For more see Supp. Method SM4.

### Supplementary Note 3: Cloud Formation Template

#### Overview

AWS CloudFormation [https://aws.amazon.com/cloudformation/] offers tools to automate the deployment of pre-specified cloud infrastructures. The central concept of CloudFormation is the *template*, which is a description of the required AWS resources, any dependencies between them, and the runtime configuration parameters required to build a particular application stack. These templates are easily shared and version-controlled, allowing customized infrastructures to be precisely encapsulated as code to create replicate cloud instances in different regions and by different user accounts.

KnowEnG has published an AWS CloudFormation template that allows users to create their own instances of the KnowEnG analysis platform using their personal AWS accounts. This capability was developed for researchers whose data or analysis profiles were not conducive to the free use public KnowEnG instance. For example, researchers with sensitive data may want to create their own copy of the KnowEnG infrastructure with additional security protocols in place. Additionally, if a user wanted to run an analysis on a very large omics spreadsheet (>200 MB) or perform intensive computation with thousands of bootstrap iterations or exhaustive parameter sweeps (prohibited on the free use public instance), they could use the template to create a replicate of the KnowEnG platform with appropriately sized compute instances and without the restrictions of the public version.

The AWS CloudFormation template that creates new deployments of the KnowEnG platform, painlessly instantiating and integrating several components:
Container pods for platform services

- A Kubernetes [https://kubernetes.io/] cluster, consisting of several virtual machines:
  - One master node
  - Compute nodes whose number and size are specified by the user
- Virtual Private Cloud isolating the Kubernetes cluster from the public Internet
- Bastion virtual machine providing secure access to the Kubernetes cluster
- Elastic File System [https://aws.amazon.com/efs/] pre-loaded with the Knowledge Network, furnishing data to all pipeline containers and storing results from all analyses
- Resilient backend deployment of KnowEnG platform using containerized pods as a Kubernetes service
- Load-balancer exposing the web interface via a URL

Researchers who use the AWS CloudFormation template to deploy their own instance of the KnowEnG platform will bear the cost of the required infrastructure and of the resources necessary for any analysis runs performed on that instance. The major cloud-computing costs of running a personal instance can be decomposed into two parts:

1. **Virtual Machines**. This is the cost of running the virtual machines in the deployment. There is one t3.micro virtual machine that serves as the “bastion” for system administration, and there is one m5.large virtual machine that serves as the Kubernetes master. Together, these two virtual machines cost $0.1064 per hour. Additionally, the Kubernetes cluster includes compute nodes, the number and size of which can be adjusted by the user. For the default configuration of two m5.large virtual machines, the total compute-node cost is $0.192 per hour, bringing the total virtual machines cost of a default deployment to $0.2984 per hour. These prices are current as of May 2019, for the us-east-1 AWS region. The AWS EC2 price tables [https://aws.amazon.com/ec2/pricing/on-demand/] provide up-to-date information, including how changes to the number and type of the compute nodes and changes of region affect the cost of the virtual machines.
2. **Storage**. This is the cost of storing cluster data on the Elastic File System. As of May 2019, AWS charges $0.30 per gigabyte per month in the us-east-1 region. Users should expect at least 40 GB of storage, which would cost less than $0.018 per hour. The AWS EFS price tables [https://aws.amazon.com/efs/pricing/] provide up-to-date information, including how changes to the amount of data stored and changes of region will affect the cost of storage.

Finally, we note that the existence of the CloudFormation template allows researchers to lower their costs by deleting their KnowEnG platform instance during periods of inactivity and redeploying it as needed.

#### Resources

KnowEnG CloudFormation Deployment GitHub repository
[https://github.com/KnowEnG/Kubernetes_AWS/tree/master/cloudformation]
KnowEnG CloudFormation template: [https://s3.amazonaws.com/knowscripts/knoweng-platform-simple.template]

### Supplementary Note 4: Knowledge Network Selection Guidance

#### Overview

The KnowEnG Analysis Platform offers users the opportunity to perform machine learning analysis on their omics datasets while making use of prior knowledge on gene interactions. This type of analysis can provide alternative, interesting results, but knowing which gene/protein interactions to include can be a daunting task. The knowledge-guided methods benefit when there are sufficient relationships between the gene entities of interest. In this supplement, we explore what types of interaction relationships provide the greatest benefit to a gene ranking task for many different types of gene sets.

The overall approach to this analysis relies on the gene ranking task performed by the DRaWR algorithm (Blatti and Sinha, 2016). The DRaWR algorithm takes an input gene set and a heterogeneous network and attempts to rank all genes in the network for their relatedness to the input gene set. The network is formed with nodes that represent genes (and proteins) and edges that represent different types of relationships between those entities. The DRaWR algorithm uses Random Walks with Restart (see Supp. Method SM3) in a manner that is similar to the KnowEnG Gene Set Characterization pipeline (see Supp. Method SM11). The difference in this application is that we are using the method to rank the relatedness of the gene nodes to the input gene set and not ranking nodes that represent known annotations of functions or pathways.

For this analysis, we provide DRaWR a single gene-gene interaction network from the KnowEnG Knowledge Network (see Supp. Method SM1) and a previously characterized set of genes. We divide the gene set into four equal sized partitions, and use 75% of the genes as the restart set in a RWR on the interaction network. We calculate the resulting gene set vector that shows a walker's long-term probability of being at any gene node in the network, and compare those values to a probability vector generated by a walker that restarts in the network randomly at any gene node. We rank genes for similarity to the gene set based on the difference between the two vectors, and find the ranking of the hidden 25% of the gene sets to calculate the Area Under the Receiver Operating Curve (AUROC). We repeat this process for each partition, for each gene set, and for each gene-gene interaction network. We summarize these results below to access which network types are more amenable to this gene ranking task and how the type of input gene set affects ranking outcome.

#### Results

In this analysis, we collected 12 different gene-gene interaction networks for comparison (see Supp. Table SN4.ST1). Each of these networks we categorized into one of five different categories: networks that capture 1) transcriptional co-expression relationships, 2) DNA/protein sequence relationships, 3) physical interactions between proteins, 4) functional relationships within pathways, and 5) co-occurrence/integrated relationships from various literature resources. These 12 networks range in size from just over 100,000 to nearly seven million network edges/relationships in human databases. We also collected six additional interaction networks from the fruitfly to compare the results across species.

The other input to this analysis was nearly 48,298 human gene sets. These gene sets were taken from 30 gene set collections available through the KnowEnG Knowledge Network. The gene set collections were also grouped into six categories: gene sets derived from 1) experimental measurements in disease/drug perturbations, 2) ontologies of gene function, 3) annotations of biological pathways, 4) identification of protein structures, 5) disruptions and measurements of transcriptional regulation, and 6) assays on different tissues and conditions. Several of these gene set collections were divided into subcollections when gene sets came from distinct sources (Supp. Table SN4.ST2). The subcollections range from 12 to 5832 gene sets with average sizes ranging from 19 to 1772 genes that are in the gene set and the interaction network.

We ran the four fold cross validation mode of DRaWR as described above for all 12 single gene-gene interaction networks and all of the input gene sets. Network edges were treated as undirected and weighted and the restart probability was set to 0.5. The second stage of DRaWR was not run, and the AUROC for each run was calculated from the DRaWR ‘diff’ score.

The results show that in this gene ranking task, the Text_Mining/Integrated (average AUROC 0.664) and Pathway (average AUROC 0.624) based interaction networks generally performed better over all gene sets (Supp. Table SN4.ST3 A). The Experimental Interaction based network performed worst (average AUROC 0.583). This trend was also true in results produced from the six fruitfly networks using fruitfly gene sets from Gene Ontology, KEGG, Pfam, and Reactome (Supp. Table SN4.ST3 B). In Supp. Table SN4.ST4, we explore how these interaction network types perform on different gene set collection types. We find that curated Pathway, Protein Domain, and Ontology (average AUROC > 0.736) types of gene sets are easier to recover than other more experimental gene sets from Disease/Drug, Regulation, and Tissue_Expression (average AUROC < 0.610) (Supp. Table SN4.ST4 A). This pattern is reasonable considering that these strong performing types of gene sets can be embedded in the Text_Mining, Pathway_Database, and Conservation network interactions. We see that the CoExpression network type does generally poorer at identifying the missing genes from any type of gene set collection. We also observe a major performance decrease for Experimental_Interaction networks on gene sets that are more experimental and less curated.

In Supp. Table SN4.ST5, we further divide the results to examine the effect of each single gene-gene network. One general trend shows that for the same category of interaction network, the more complete networks provide the greatest benefit. For example, STRING Function Database (391,679 edges) is the best Pathway_Database network and STRING Co-expression (1,400,944 edges) is the best Experimental_Interaction network. When considering the subcollections of gene sets, we can see that general trends do not always predict performance, but can provide valuable guidance (Supp. Table SN4.ST6). A Disease/Drug gene set collection like OMIM (Hamosh, et al., 2005) gene sets is extremely well captured by Text_Mining/Integrated networks (AUROC 0.904) and very poorly captured by CoExpression networks (AUROC 0.581). On the other hand the MSigDB Cancer Gene Neighborhood gene sets (Liberzon, et al., 2011) are captured by both network types equally well (AUROC ~0.72) and better than any other type. Co-expression networks (and Pathway networks) also seem to perform best for many different types of gene sets derived from the Gene Expression Omnibus (Barrett, et al., 2012), which perhaps represent the common case of experimentally defined differential expression gene sets.

Overall, when choosing an interaction network, if your gene set is defined by careful selection or annotated genes, it may be best to consider a Text_Mining/Integrated Network. In general, the more complete the network (in terms of number of edges) given its category, the more likely it will perform well in analysis tasks. Finally, co-expression networks seem to have a strong value specifically for differential expression gene sets derived from experimental assays.

#### Resources

DRaWR GitHub Repo [https://github.com/KnowEnG/DRaWR ]
Knowledge Network Content Summary [https://knoweng.org/kn-data-references/ ]
Knowledge Network Downloadable Content Links
[https://github.com/KnowEnG/KN_Fetcher/blob/master/Contents.md ]

### Supplementary Note 5: Knowledge Network Retrieval and Mapping

#### Overview

The KnowEnG Center has developed two valuable utilities in order to provide access and interoperability for users to our Knowledge Network of prior knowledge about genes and proteins. Both of these utilities, the KN_Fetcher and KN_Mapper, are developed in Python, and made available as auto-building Docker images on DockerHub. Both also contain a Common Workflow Language (CWL) description, making them executable on many standardized workflow runners and amenable for workflow visualization tools and user interfaces. These utilities with their GitHub, Docker, and CWL references are registered in the Dockstore developed by the Cancer Genome Collaboratory [https://dockstore.org/ ].

##### KN_Fetcher Utility

The first utility, the KN_Fetcher, is a tool that allows users to download a specific subnetwork of the Knowledge Network that relates to a single species and single edge type. In the version of the Knowledge Network used in this publication, there are 185 different subnetworks of this type. This tool can be helpful when examining the downloadable results from the KnowEnG platform, retrieving the networks used in a pipeline run for analysis and visualization. The utility is also valuable for users who wish to run the KnowEnG Analysis Pipelines outside the platform because it retrieves the network data already transformed into the appropriate format. Being able to run the KnowEnG prior knowledge-guided analysis pipelines in the SevenBridges Cancer Genomics Cloud is possible in part because the KN_Fetcher is able to pull the relevant subsections of the Knowledge Network onto the SevenBridges cloud. The KN_Fetcher asks the user to supply a Knowledge Network version, species taxonomic identifier (Supp. Table SM1.ST1) and a Knowledge Network edge type [https://knoweng.org/kn-data-references/#kn_contents_by_gene-gene_edge_type ]. It then uses Amazon Web Services (AWS) tools to retrieve three files that relate to that subnetwork from AWS Simple Storage Service (S3) [https://aws.amazon.com/s3/ ]. The returned files are the file listing the subnetwork edges, a file that contains mapping information for all of the subnetwork nodes, and finally a file that contains the metadata of the subnetwork including the provenance of the original external data files processed, information about the build command and date, and network statistics about the subnetwork. For users who are willing to manually download subnetwork files one at a time for the Knowledge Network build used in this publication, direct links can be accessed from our contents summary page [https://github.com/KnowEnG/KN_Fetcher/blob/master/Contents.md ]

##### KN_Mapper Utility

The second utility, the KN_Mapper, is a tool that allows users to provide a list of gene names or identifiers, and retrieve the Ensembl stable identifiers that are used internally in the KnowEnG network-guided analysis pipelines. In the current version, this tool enables the mapping of over 48 million gene, transcript, and protein names and identifiers to stable identifiers for 583,933 genes across 20 species. The KN_Mapper tool is integrated into the Data Cleanup Pipeline and run before any knowledge-guided analysis in the KnowEnG Platform or on the SevenBridges Cancer Genomics Cloud. It can be used by any user who wishes to harmonize the entities of their dataset with the entities recorded in the KnowEnG Knowledge Networks. The inputs to this tool are a file that contains one original gene name on each line, as well as optional hints about the mapping, such as species taxonomic identifier. The tool then queries an in-memory Redis key-value database [https://redis.io/ ]. It outputs a file with the same number of lines where each line corresponds to the mapped Ensembl stable identifier, gene symbol, and full gene description. If no stable identifiers are found for the original query, the flag “unmapped-none” is returned. If multiple, different stable identifiers are found for the same query string, the flag “unmapped-many” is returned. The tool queries the submitted gene names from the current Knowledge Network Redis database, available at port 6379 at knowredis.knoweng.org. Instructions for setting up a local Redis database can be found at the GitHub readme [https://github.com/KnowEnG/KN_Mapper ].

#### Resources

KnowEnG KN Tools Page [https://knoweng.org/kn-tools/ ]

##### Knowledge Network Fetcher Tool

GitHub Repository [https://github.com/KnowEnG/KN_Fetcher ]
Docker Image [https://hub.docker.com/r/knoweng/kn_fetcher ]
Dockstore Reference [https://dockstore.org/containers/quay.io/cblatti3/kn_fetcher ]

##### Knowledge Network Mapper Tool

GitHub Repository [https://github.com/KnowEnG/KN_Mapper ]
Docker Image [https://hub.docker.com/r/knoweng/kn_mapper ]
Dockstore Reference [https://dockstore.org/containers/quay.io/cblatti3/kn_mapper ]

### Supplementary Note 6: Reproducing Major Analyses in KnowEnG Platform

#### Overview

In this paper, we have demonstrated several bioinformatics analyses such as patient stratification, gene prioritization, gene set characterization and signature analysis on a few major data sets in cancer genomics (TCGA and METABRIC). In doing so, we have reproduced key results from the original studies (Emad, et al., 2017; Hoadley, et al., 2014; The Cancer Genome Atlas Research, et al., 2017) as well as gleaning new biological insights. In this Supplemental Note, we provide resources for users who wish to reproduce the majority of these analyses for themselves in the KnowEnG platform. In doing so, we highlighted both the sophisticated level of analysis possible and the ease-of-use with which multiple pipelines can be invoked, individually as well as in combination, to generate a multi-faceted narrative of the insights that the data have to offer.

We chose eight analyses (Supp. Table SN6.ST1) to highlight that could be recreated in the KnowEnG Analysis Platform (or the SevenBridges Cancer Genomics Cloud). Instructions for running these analyses are found in the README of the repository with the input data [https://github.com/KnowEnG/quickstart-demos/tree/master/publication_data/blatti_et_al_2019].

For each analysis, we first let the user know which pipeline is applicable. We provide the input spreadsheets(s) for that pipeline which must be uploaded into the user's account on the platform. When there are multiple input files for a step, the "File Type" provided in the repository data table indicate for which step of pipeline configuration each input is intended. All the files can be downloaded from a single zipped archive [https://s3.amazonaws.com/knoweng-publication-data/blatti_et_al_2019.tar.gz], individually from the spreadsheets directory in the repository [https://github.com/KnowEnG/quickstart-demos/blob/master/publication_data/blatti_et_al_2019/spreadsheets], or from the links in the repository's input data tables. Files larger than 1 MB will need to be unzipped before loading them into the platform.

In the README for each analysis, we provide the parameters of the pipeline that are needed to recreate the highlighted analysis in the paper. These parameters appear in the order and with the vocabulary found in the KnowEnG Analysis Platform. (The parameters names and requirements will likely differ in the SevenBridges Cancer Genomics Cloud depending on the pipeline). Once an analysis is submitted and finished in the platform, the pipeline visualization and several downloadable outputs will be produced. We have also highlighted the primary downloadable output of interest as part of the README. Often, this primary downloadable output is also the input to the next analysis example. Besides the eight exemplary analyses provided, most of the baseline and alternative analysis described in the paper can be recreated with these resources by modifying the Knowledge Network related parameters.

#### Resources

Data for Recreating Analyses [https://s3.amazonaws.com/knoweng-publication-data/blatti_et_al_2019.tar.gz]
Instructions for Recreating Analyses [https://github.com/KnowEnG/quickstart-demos/tree/master/publication_data/blatti_et_al_2019].

### Supplementary Note 7: Characterization of Knowledge Guided Mutation Subtypes

#### Background

In our first case study, we presented the clustering of pan-cancer patient mutation profiles using the network based stratification (NBS) (Hofree, et al., 2013) method (see Supp. Method SM3) and found that this yields better clustering of mutation data, with more size-balanced clusters and with survival analysis yielding stronger separations. We also noted that these clusters do not simply recapitulate the tumor types of patients, suggesting that the discovered clusters may reveal new structures in the data.

To explore this further and determine the molecular and functional characteristics of these clusters, if any, we identified the genes and pathways whose mutation status/counts are most discriminative of each cluster. We began by extracting the genes by patient matrix that had been subjected to clustering whose columns of mutation data have been transformed with into network-smoothed feature vectors using random walks with restart (RWR). For each of the 14 clusters in our 'hnInt' knowledge-guided clustering using the HumanNet Integrated network (Lee, et al., 2011), we performed t-tests (using KnowEnG's standard Gene Prioritization from Supp. Method SM7) to identify the top 100 genes differentially mutated between the given cluster and all other clusters. We then performed apply the Fisher's exact test to this gene sets with our Gene Set Characterization pipeline (see Supp. Method 11) to identify Gene Ontology (Ashburner, et al., 2000), Reactome (Fabregat, et al., 2018), Pathway Commons, and NCI pathways (downloaded from Enrichr (Chen, et al., 2013)) enriched with these 100 genes. We repeated this for all 14 clusters separately.

#### Results

Supp. Table SN7.ST1 shows basic information about the discovered clusters (rows) — their sizes, and how many patients of each cluster belongs to which tumor type. We note that clusters vary in size, between 3 and 993, with mean, median, and standard deviation of sizes being 234, 60, and 324 respectively. We also note that most clusters are mixed in terms of tumor types, but some are more dominated by a few tumor types, e.g., c2 is almost entirely AML, c3 is dominated by lung cancers (LUAD, LUSC), c4 is mostly breast and uterine. The two largest clusters are c0 (enriched in ovarian cancers) and c1 (enriched in kidney cancers).

Supp. Table SN7.ST2 and Supp. Table SN7.ST3 show genes that are most discriminative of each cluster and their enriched properties respectively. For example,

A. c0 and c1, the largest clusters, are enriched for general tumorigenic properties and pathways related to p53 and apoptosis.
B. Among the several genes whose mutation status is uniquely discriminative of c0 was SOCS1, an important tumor suppressor that attenuates cytokine signaling (Sutherland, et al., 2004). This is particularly interesting because the two most over-represented tumor types in c0 are ovarian and breast cancers, where prolactin and IL-6 secretion from the ovary and mammary gland induce growth by stimulating JAK/STAT signaling. The third largest cancer type in cluster c0 is head & neck cancer, where a high level of JAK/STAT signaling is also known to be important. Cluster c0 is also discriminated from other clusters by ARID1A mutation status. ARID1A is an important SWI/SNF tumor suppressor (Guan, et al., 2011).
C. Cluster c1, which is highly overrepresented in kidney renal clear cell carcinoma, was uniquely marked by S100A16, S100A6, S100A5, and S100A3, among other genes. The finding of S100A as a marker of this cluster is potentially significant, given its tumor type composition (kidney and breast) (Takashi, et al., 1988).
D. Cluster c2 is marked by mutation status of genes in the HIF-1-alpha and HIPK20-related pathways, which is consistent with its AML-dominated composition (Velasco-Hernandez, et al., 2014).
E. Cluster c4 includes mainly breast and uterine carcinoma patients, and is associated with PIP3, PI3K/AKT and GAB1-related pathways, in agreement with prior knowledge about those tumor types (Myers, 2013).
F. Cluster c5 is associated with titin binding, muscle-related pathways.
G. Cluster c7 is associated with sarcoplasmic reticulum.
H. Cluster c8 is associated with protein autophosphorylation.
I. Cluster c9 is associated with neuronal action potential, KEAP1-NFE2L2.
J. Cluster c10 is associated with histone methyltransferase activity (H3-K36 specific).

In other words, the discovered clusters are characterized by mutations in genes from specific and distinct pathways, even when they are mixed in terms of tumor type representation.

We also paid special attention to the characteristics of the two largest clusters c0 and c1, both of which include a number of breast cancer patients but also include a large number of ovarian and kidney cancer patients respectively. In particular, we examined genes whose mutation status is discriminative between these clusters, and found that the genes marking cluster c1 are strongly enriched in calcium ion binding.

### Supplementary Note 8: Capabilities of the Spreadsheet Visualizer

#### Overview

Often, the first objective when beginning data analysis is to get a sense of what the data look like – *are there any striking imbalances between cohorts? any striking correlations with certain phenotypes? any missing data? any skewed data?* It is not easy to grasp this information efficiently, and it is equally difficult for investigators to refresh their memories as they come back to prior studies. We developed a tool, Spreadsheet Visualizer, to address these challenges using descriptive statistics and interactive visuals (Supp. Fig. SN8.SF1).

The Spreadsheet Visualizer provides a flexible method for comparing multiple data types from the same patient samples, such as comparing between demographic and gene data, or other ‘omics data. We translate the values in these spreadsheets to colored “heatmaps” and provide many filtering and display options allowing investigators to check the data for errors and identify any obvious correlations between data types and phenotypes.

This tool is valuable at multiple stages of the data analysis process. As described, an initial understanding of the data ***prior to analysis*** can identify the need for further data cleaning and data transformation, and assist in developing hypotheses to be tested with computational analysis. The tool can also be helpful in studying the ***results of analysis*** by selecting spreadsheets derived from other pipelines such as Gene Prioritization. The tool is also helpful to convey ***reportable observations*** for publications and presentations.

#### Methods

The Spreadsheet Visualizer allows users to select multiple spreadsheets of related data that have been uploaded to the KnowEnG platform. See Supp. Note SN1 for technical description. The pipeline executes standard variation and correlation statistics, distribution analysis and Kaplan-Meier time-to-event curves. An interactive visualization provides fluid exploration of the loaded data.

##### Data Input

When using multiple spreadsheets, each spreadsheet must represent the same sample population. Spreadsheet orientation does not matter—that is, samples can correspond to rows or to columns—as long as the sample identifiers are consistent between files and all files have at least 50% of their samples in common. It is best to separate different data types into their own spreadsheets. For example, if a study includes gene expression data, somatic mutation data, and clinical data, the spreadsheet visualizer will be most effective if the data are organized as three spreadsheets, one containing the gene-expression data, one containing the somatic mutation data, and one containing the clinical data. When studying a single spreadsheet, it is best if the spreadsheet is oriented such that columns correspond to samples. Launching the pipeline is simple and requires the user to select the spreadsheets of interest (Supp. Fig. SN8.SF2).

#### User Interface Highlights

The tool provides both a qualitative overview of the data, as well as quantitative details, and supports the ability to fluidly shift focus and reorganize data, based on selected phenotypes and/or categorical features. This interface is rich with information design techniques including the following highlights:

##### Data Comparison

Central to this visualization is the ability to compare multiple spreadsheets for the same study samples. For example, one of the heatmaps can display expression values of top genes, and another can show DNA methylation. All heatmaps align so that the columns across all visuals represent the same sample (e.g. patient). The biologist can choose a specific phenotype from their data, such as “Cancer Stage”, that is displayed as a single row heatmap at the top of the view. This “Group By” phenotype defines the grouping of all the samples (columns) in all the heatmaps. A secondary phenotype can be selected for sub-sorting the columns within each group. Additional single row heatmaps, or *“data-strips”,* can be displayed for a single variable and appear below the heatmaps in the same alignment for easy comparison of data. These variables are selected from any of the loaded spreadsheets and can be either categorical data such as “survival status”, or continuous variables such as “blood pressure” (Supp. Fig. SN8.SF3). The score beside each of these single-row heatmaps reflects the strength of the statistical association between the row and the “Group By” category that has been selected at the top of the visualization. For rows of continuous data, we apply an ANOVA test, and for rows of categorical data, we apply a Chi-squared test. In both cases, the displayed score is the negative log10(pvalue), capped at 200.

The rows of the multi-row heatmaps represent the data values for each sample. For example, the rows may represent genes and their expression value per sample. These displayed rows can be sorted by row variance or by significance of correlation to the displayed *Group by* phenotype. This correlation score is determined by an ANOVA test. The quantity of rows displayed can be determined by these same measures. The biologist can choose to display the top *N* rows, or view a graph depicting the score for each variable to better determine a meaningful threshold for display (Supp. Fig. SN8.SF4).

##### Data Distribution

Understanding how the data are distributed, and the relationships between distributions based on various phenotypes, provides a richer understanding of the data. Small graphs are included adjacent to each *data-strip* (Supp. Fig. SN8.SF5). providing a quick reference to the distribution of values for that variable. Quantitative details are provided by rolling over segments of the graph (Supp. Fig. SN8.SF6). These small yet powerful graphs facilitate visual skimming of the data – a task that is not often addressed by other tools. Clicking on the *data-strips* displays statistical details of the distribution, including the distribution based on the *Grouping Phenotype* displayed at the very top. This is recomputed each time the user re-sorts using other phenotypes of interest for grouping (Supp. Fig. SN8.SF7).

##### Heatmaps

Heatmaps for large spreadsheets of data provide a way to comprehend the numerical values across many variables and samples. The visualization allows users to explore the data at multiple resolutions: (1) all samples at once in a "*Birdseye View",* (2) only the samples in a selected group, or (3) "pages" of 250 samples at a time. Whenever the number of displayed samples exceeds the number of horizontal pixels available, cell colors will be blended to provide a sense of the data values on a single screen; i.e., without horizontal scrolling. The researcher can then toggle between the zoomed-out view and a zoomed-in view using the navigator (Supp. Fig. SN8.SF8). Interactive rollovers for each cell of the heatmap help researchers find specific detail about the data, such as sample ID, gene name and expression value (Supp. Fig. SN8.SF9). Color map choices are provided so the user can choose a color system that works best with their data (Supp. Fig. SN8.SF10).

##### Time-to-Event Curves

For spreadsheets that contain time-to-event measures, such as years of survival, the researcher can generate a Kaplan-Meier analysis for any categorical phenotype of interest. To generate curves, the researcher clicks on the clock icon adjacent to that single-row heatmap. This opens a popup where the time variable and event variable are selected and the graph is created (Supp. Fig. SN8.SF5). For example, a graph is easily computed to compare survival between "Anatomic Neoplasm Subdivision" groupings. The graph includes a log-rank-p-value that is a summary statistic capturing the extent to which the differences between the groups' curves are significant (Supp. Fig. SN8.SF11).

### Supplementary Note 9: Pan-Cancer Signature from Prioritized Genes

#### Background

The second case study was about using knowledge-guided Gene Prioritization in order to identify lists of 100 genes that were specific to each pancan12 cancer disease type (Supp. Method SM7). We then sought to identify whether these sets of prioritized genes were able to form a 'signature' for tumor type, i.e., a representative collection of genes that captures much of the diagnostic or prognostic value of the entire expression profile. In order to perform this analysis, we created a union gene set from the twelve tumor type specific top100 gene lists. We repeated this process for all three gene prioritization analyses each run with different prior knowledge,

1) no network, 'noNet' (Supp. Table SM7.ST1),
2) HumanNet Integrated network, 'hnInt' (Supp. Table SM7.ST2).
3) STRING TextMining, 'SText' (Supp. Table SM7.ST3)

These three union gene sets resulted in 993, 985, and 963 genes respectively. We then used our standard hierarchical-based Sample Clustering (Supp. Method SM3) with 'Euclidean' affinity and 'Ward' linkage separately to find 16 clusters (as done in original pancan12 analysis (Hoadley, et al., 2014)) using the full pancan12 14373 gene expression matrix (expr_All), and that expression matrix subsetted for each of our three union gene sets (expr_gp100_noNet, expr_gp100_hnInt, expr_gp100_sText).

#### Results

Supp. Table SN9.ST1 shows the details and the Kaplan-Meier survival analysis p-values for the clusters resulting from these four sample clusterings. Indeed, we observed that pan-cancer subtypes obtained from clustering only the expression of the HumanNet Integrated network informed tumor-associated genes were just as predictive of survival (Kaplan Meier p-value 3.8E-175, Supp Fig SN9.SF1) as the above-mentioned clusters based on entire expression profiles (p-value 1.2E-169). This result held whether we used a different network, STRING TextMining, or no network in the Gene Prioritization step. This means we can reduce from ~14K to 1K genes without losing the ability to identify meaningful clusters that relate to overall survival. Interestingly, although the clusters identified by this method had slightly improved survival outcome predictive ability, they were not as strongly related to tumor type as the clusters identified with the expression data and the Cluster-Of-Cluster-Assignments (COCA) in the original paper (Hoadley, et al., 2014). Neither were these new clusters strongly similar to each other, suggesting that each different modality of Gene Prioritization enable the discovery of a different by equally good clustering that relates to survival separation (Supp Table SN9.ST2).

### Supplementary Note 10: KnowEnG Analysis on SB-CGC using Docker and CWL

#### Overview

##### Seven Bridges Cancer Genomics Cloud (SB-CGC)

The Cancer Genomics Cloud (CGC) [https://cgc.sbgenomics.com/], built by Seven Bridges Genomics as part of the National Cancer Institute's Cancer Genomics Cloud pilot program, is a cloud-based environment that hosts large genomic data sets (e.g., TCGA) as well as genomic analysis applications. These apps can be executed on the CGC, which uses Amazon's AWS behind the scenes.

Apps are either **command line tools** or **workflows** built up from other apps. They can be developed on, or imported to, the CGC site. KnowEnG [https://knoweng.org/] has created apps on the CGC for several of the pipelines we have developed.

Supp. Fig. SN10.SF1 shows one tool imported from KnowEnG, Gene Set Characterization (GSC), being edited directly on the CGC Tool Editor web interface. The execution of this pipeline tool is specified with an appropriate command line call and the associated GSC Docker image developed by KnowEnG. Besides the Tool Editor interface, CGC also provides the Rabix Composer IDE that allows editing and testing CGC apps on one's own local machine.

In general, for the KnowEnG pipelines, both a command line tool and a workflow version are necessary. The latter involves collecting all the inputs for the pipeline and modifying and distributing them as necessary to the different components of the workflow, running all the components of the workflow, and collecting and modifying the outputs.

Figures Supp. Fig. SN10.SF2 and Supp. Fig. SN10.SF3 show the workflow version of the GSC pipeline. Supp. Fig. SN10.SF2 shows the app's description, and Supp. Fig. SN10.SF3 shows creating/editing the app on the CGC. For a workflow, this mostly involves dragging and dropping icons representing apps (command line tools or workflows), connecting them as needed, and configuring the inputs and outputs of the overall workflow and the internal components.

The CGC also provides some public analysis tools of their own, and allows apps developed on their site to be published and made available to others. KnowEnG has published several of their analysis pipelines on the CGC. Supp. Fig. SN10.SF4 shows the KnowEnG apps on the CGC.

##### Technologies for Importing Workflows

Two of the main technologies that allow for tools to be developed on, imported to, and run on the CGC are Docker and the Common Workflow Language (CWL).

Docker [https://www.docker.com/] provides containerization, a form of virtualization that allows for capturing all of the system, environment, library, and code dependencies an application needs in a standard, lightweight object. Docker containers can be executed in any computation environment.

When a command line tool is developed on the CGC, a Docker container is specified. This can be seen in Supp. Fig. SN10.SF1, where the container is specified in the field **Docker Repository[:Tag]**. This container is used when the tool is executed.

CWL [https://www.commonwl.org/] is a language for describing analysis tools and workflows. It specifies the details of the inputs, outputs, and execution steps of the tools. This allows for making the tools portable and scalable across of variety of software and hardware environments. There are many execution engines for CWL, taking a tool's CWL description and executing it.

Normally, apps on the CGC are developed interactively and graphically, and the CWL is created behind the scenes. But it is also possible to import an app already described in CWL. Similarly, it is possible to export an app from the CGC into a pure text CWL format. Supp. Fig. SN10.SF5 shows a portion of the CWL description of the GSC command line tool app.

#### Methods

##### Run Combined, Multi-pipeline Workflow

As stated earlier, KnowEnG has made available several of our pipelines as workflows on the CGC. These pipelines are useful by themselves, but additionally they can be used in conjunction to create more complex workflows.

To achieve this, one possibility is to run each workflow individually, using outputs from some workflow runs as inputs to others. With just a little bit of work on the CGC, however, it is possible to relatively quickly create such larger workflows as apps themselves from the individual lower level workflows.

Supp. Fig. SN10.SF6 shows an example of this, where the KnowEnG Analysis of LUSC Subtypes (KALS) workflow is built up from several of the other KnowEnG workflows.

Supp. Fig. SN10.SF7 shows editing the KALS workflow, with the node for the GSC workflow highlighted. When integrating an app into a workflow, one can choose whether to hardcode values for that app's input parameters, or to "port" them and make them available as inputs in the new, larger workflow. Similarly, one can choose which of the app's outputs to make available.

This document [https://github.com/KnowEnG/KnowEnG_CWL/tree/master/CGC] describes how to run the workflows on the CGC, both individually and the combined workflow directly.

##### Sharing and Reproducibility

In publishing these tools on the CGC, KnowEnG is working towards the "FAIR" principles: Findable, Accessible, Interoperable, and Reusable. They are findable through the CGC, as well as Docker Hub and GitHub. The versions on the CGC are accessible to researchers, ready to use with the public data sets available there or with their own data. They are interoperable with other tools and workflows available on the CGC and they are reusable in a reproducible manner with the other CGC datasets and collections in the KnowEnG Knowledge Network.

Users are able to run KnowEnG tools singly or in combination. As an example, we published a multi-tool workflow, the aforementioned KALS workflow that performs knowledge-guided enrichment analysis on sets of genes that are differentially expressed between TCGA samples that map to different genomic subtypes.

One could easily envision bringing additional custom collections of knowledge bases into this analysis framework using the Docker and CWL paradigm demonstrated here. By seamlessly linking petabyte-scale data access to the full suite of bioinformatics tools and advanced analytics available on the CGC and KnowEnG platforms, an even more valuable data analysis ecosystem is made available to researchers worldwide.

#### Resources

KnowEnG KnowEnG_CWL GitHub repo
[https://github.com/KnowEnG/KnowEnG_CWL/tree/master/CGC]
KnowEnG Public Apps on the CGC
[https://cgc.sbgenomics.com/public/apps#q?search=knoweng]

#### Supplementary Note 11: Consistency Analysis of DRaWR

### Supplementary Notes Tables

**Supplementary Table SN4.ST1. Gene-Gene Interaction Networks.**
Listed in each row are the different gene-gene interaction network that were assessed in this analysis. Additional information about each network is reported including its category, identifier, and the number of network edges in the human and fruitfly.

**Supplementary Table SN4.ST2. Gene Set Collections.**
Listed in each row are the different gene set collections that were assessed in this analysis. Additional information about each collection includes the collection category, identifier, and the aliases of possible subcollections. For each collection, we list the number of gene sets and the average size of those sets after mapping them to the gene-gene interaction network.

**Supplementary Table SN4.ST3. AUROC by Interaction Network.**
The average AUROC across four folds for every gene set is listed for each interaction network as well as the number of folds averaged. The interaction networks are grouped and summarized by network category. Results are shown for A) human and B) fruitfly.

**Supplementary Table SN4.ST4. AUROC by Gene Set Collection Type and Network Type.**
The average AUROC across four folds for every gene set is listed for each gene set collection type (rows) and interaction network type (columns) for the human only runs.

**Supplementary Table SN4.ST5. AUROC by Interaction Network and Gene Set Collection Type.**
The average AUROC across four folds for every gene set is listed for each gene set collection type (columns) and interaction network (rows) for the human only runs. The interaction networks are grouped and summarized by network category.

**Supplementary Table SN4.ST6. AUROC by Gene Set Collection and Network Type.**
The average AUROC across four folds for every gene set is listed for each each gene set subcollection (rows) and interaction network type (columns) for the human only runs. The gene set subcollections are grouped by gene set collection type.

**Supplementary Table SN6.ST1. Analyses for Reproduction in the KnowEnG Platform.**
We highlight eight primary analyses whose datasets and parameters necessary for reproduction in the KnowEnG platform are provided. For each analyses, we list its name, the pipeline it requires, the source of its input data, and the supplementary methods section where more details can be found.

**Supplementary Table SN7.ST1. 'hnInt' Clusters by Disease Type.**
For each of the 14 clusters (rows) from our knowledge guided clustering of the somatic mutation data using the HumanNet Integrated network, we show the number of samples from each tumor disease type (columns).

**Supplementary Table SN7.ST2. 'hnInt' Cluster Specific Genes.**
For each of the 14 clusters from our knowledge guided clustering using the HumanNet Integrated network, we found the set of top 100 genes (rows) according to the a t-test on the network smoothed mutation values between the samples within each cluster (columns) vs all others. Membership in each of these genes sets is indicated with a table value of 1.

**Supplementary Table SN7.ST1. Top Pathways Enriched in Genes Specific for 'hnInt' Clusters.**
For each top 100 cluster specific gene list from our 'hnInt' clustering, we return the top 5 enriched pathways using the one-sided Fisher's exact test (reported as the negative log10 p-value (neglog pval)). The number of genes annotated for these pathways (pathway size) and the source of the information is also reported (collection).

**Supplementary Table SN9.ST1. Analysis of Clustering with TCGA Expression Data.**
Shows statistics for various methods for clustering the gene expression data. The first column ('Alias') shows the name given to each clustering run. The first three rows show clusterings from the original PANCAN12 paper analysis and the last four rows show the results of our General Clustering pipeline using Ward linkage with Euclidean affinity on the PANCAN12 gene expression data. The final three rows only use genes that were in the top 100 prioritized gene lists for disease type. We report the number of clusters ('nClust'), the number of gene features used in the clustering ('nFeatures') and the Kaplan-Meier p-value of the significance of the relationship between the clustering and survival outcome.

**Supplementary Table SN9.ST2. Similarity Between Expression Clusterings.**
Shows the adjusted Rand Index for each pairing of the seven selected clusterings in this analysis: “disease” - grouping by TGCA primary disease, and “tcga_expr” and “tcga_coca”, expression-only and COCA clustering analysis in original TCGA paper, and “expr_gp100_hnInt”, “expr_gp100_sText”, and “expr_gp100_noNet”- clustering of expression data using genes from gene prioritization for disease type with HumanNet Integrated, STRING Textmining, and no prior knowledge networks respectively.

### Supplementary Notes References

Ashburner, M., *et al.* Gene ontology: tool for the unification of biology. The Gene Ontology Consortium. *Nat Genet* 2000;25(1):25-29.

Baker, M.P. and Bushell, C. After the storm: Considerations for information visualization. *IEEE Computer Graphics and Applications* 1995;15(3):12-15.

Barrett, T., *et al.* NCBI GEO: archive for functional genomics data sets—update. *Nucleic acids research* 2012;41(D1):D991-D995.

Blatti, C. and Sinha, S. Characterizing gene sets using discriminative random walks with restart on heterogeneous biological networks. *Bioinformatics* 2016;32(14):2167-2175.

Chen, E.Y., *et al.* Enrichr: interactive and collaborative HTML5 gene list enrichment analysis tool. *BMC Bioinformatics* 2013;14:128.

Emad, A., *et al.* An epithelial-mesenchymal-amoeboid transition gene signature reveals molecular subtypes of breast cancer progression and metastasis. *bioRxiv* 2017:219410.

Fabregat, A., *et al.* The Reactome Pathway Knowledgebase. *Nucleic Acids Res* 2018;46(D1):D649-D655.

Guan, B., Wang, T.-L. and Shih, I.-M. ARID1A, a factor that promotes formation of SWI/SNF-mediated chromatin remodeling, is a tumor suppressor in gynecologic cancers. *Cancer research* 2011;71(21):6718-6727.

Hamosh, A., *et al.* Online Mendelian Inheritance in Man (OMIM), a knowledgebase of human genes and genetic disorders. *Nucleic acids research* 2005;33(suppl_1):D514-D517.

Hoadley, K.A., *et al.* Multiplatform analysis of 12 cancer types reveals molecular classification within and across tissues of origin. *Cell* 2014;158(4):929-944.

Hofree, M., *et al.* Network-based stratification of tumor mutations. *Nat Methods* 2013;10(11):1108-1115.

Lee, I., *et al.* Prioritizing candidate disease genes by network-based boosting of genome-wide association data. *Genome Res* 2011;21(7):1109-1121.

Liberzon, A., *et al.* Molecular signatures database (MSigDB) 3.0. *Bioinformatics* 2011;27(12):1739-1740.

Myers, A.P. New strategies in endometrial cancer: targeting the PI3K/mTOR pathway—the devil is in the details. *Clinical Cancer Research* 2013;19(19):5264-5274.

Sutherland, K.D., *et al.* Differential hypermethylation of SOCS genes in ovarian and breast carcinomas. *Oncogene* 2004;23(46):7726.

Takashi, M., *et al.* An immunochemical and immunohistochemical study of S100 protein in renal cell carcinoma. *Cancer* 1988;61(5):889-895.

The Cancer Genome Atlas Research, N., *et al.* Integrated genomic characterization of oesophageal carcinoma. *Nature* 2017;541:169.

Tufte, E.R., *et al.* Visual explanations: images and quantities, evidence and narrative. In.: AIP; 1998.

Velasco-Hernandez, T., *et al.* HIF-1α can act as a tumor suppressor gene in murine acute myeloid leukemia. *Blood* 2014;124(24):3597-3607.
