## Supplementary Notes Figures for "Knowledge-guided analysis of ‘omics’ data using the KnowEnG cloud platform"

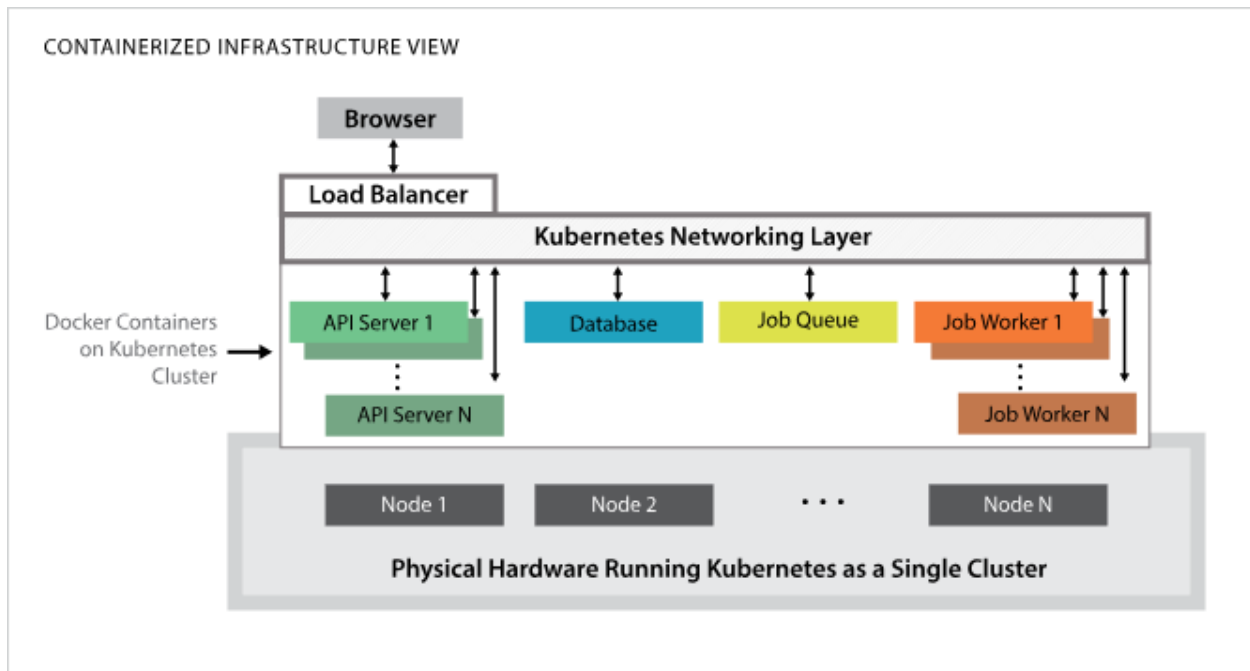

**Supplementary Figure SN1.SF1. KnowEnG Platform Components.**

Components are isolated in Docker containers, indicated by the colored boxes. The containers are run by a Kubernetes orchestration system over multiple nodes. The number of cluster nodes automatically adjusts in response to user traffic.

**knoweng** Analysis Pipelines Data Support Lisa Gatzke

### SELECT A PIPELINE

- **Sample Clustering** Start Pipeline
- Feature Prioritization
- Gene Set Characterization
- Signature Analysis
- Spreadsheet Visualization

**About Sample Clustering**

You have a spreadsheet that contains "omics" data signatures of multiple biological samples. You want to find clusters of the samples that have specific "omics" signatures (e.g. subtypes of cancer patients). You may also have phenotypic measurements (e.g., drug response, patient survival, etc.) for each sample. In this case, you want to find "omics" subtypes related to the important phenotypic measurements. This pipeline uses different clustering algorithms to group the samples and scores the clusters in relation to the provided phenotypes. This pipeline also provides methods for finding robust clusters and/or using the Knowledge Network to transform the genomic signatures of the samples, if the features are genes.

How does this pipeline work?

How does this pipeline differ from standard clustering?

What input does the pipeline need?

What output does the pipeline produce?

Info and training

**Supplementary Figure SN2.SF1. Pipeline Home.**

KnowEnG offers five pipelines that are commonly used in "omics" analyses. Frequently asked questions about each pipeline are addressed in the Info Panel on the right.

**knoweng** Analysis Pipelines Data Support Lisa Gatzke

Pipeline: **sample\_clustering-2018-11-2** Features File Response File **Network** Parameters Bootstrapping Review & Submission Cancel

Reset Defaults

#### Do you want to use the Knowledge Network?

**Yes** **No**

- Select species**
 default
- Select Interaction Network for analysis**
 default
- Choose the amount of network smoothing**
 % default

**What does it mean to use the Knowledge Network?**

Briefly, if the features are genes, the network makes use of known relationships such as protein-protein interactions, genetic interactions and homology relationships among genes. These relationships enable the pipeline to transform the genomic signatures of each sample by integrating each gene's signal with the signals from its interacting neighbors. This transformation can potentially aid in the correct clustering of samples by propagating weak individual gene signals to higher levels (pathways and modules) where the sample similarity is stronger.

**Advantages**

**Disadvantages**

**What if my data's species isn't an option in the list?**

**What is an Interaction Network?**

**What is Network Smoothing?**

**Info and training**

**Previous** **Next**

#### Supplementary Figure SN2.SF2. Pipeline Setup Workflow.

Wizard-like interface that steps users through pipeline setup, including contextually relevant information provided in the Info Panel for each step. The above image shows the Network selections step in the Sample Clustering Pipeline.

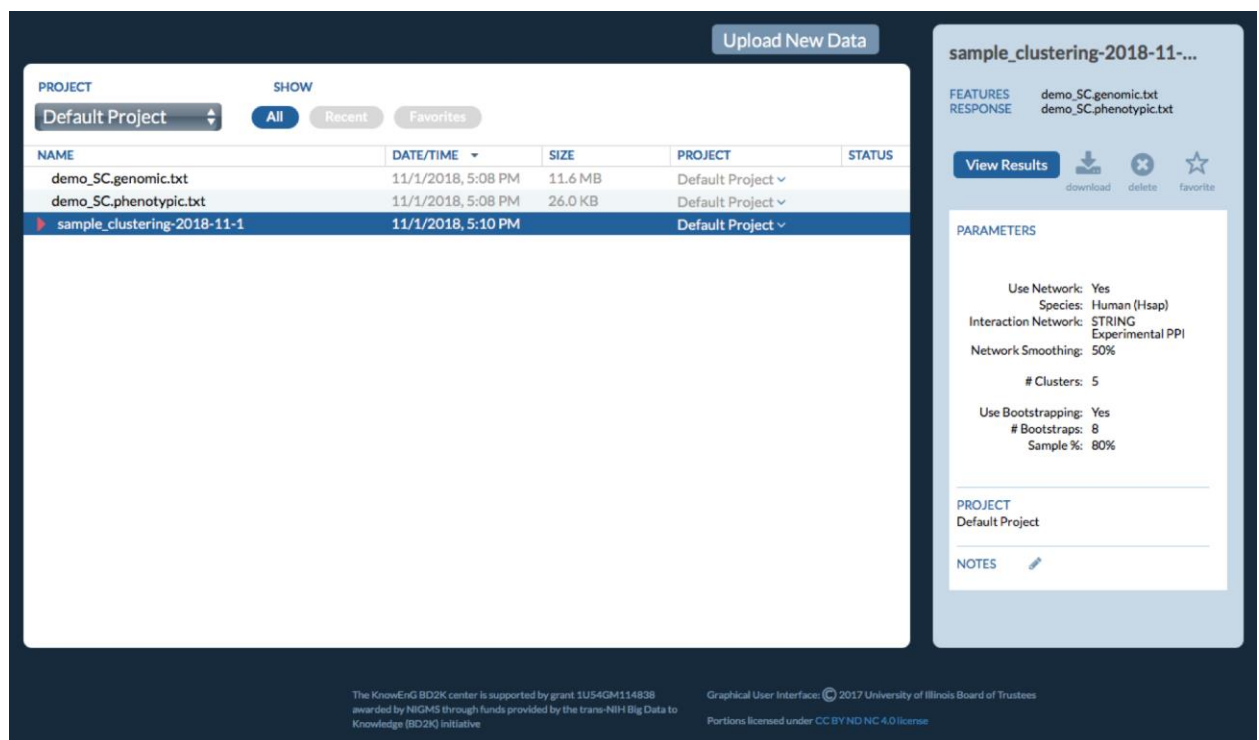

**Supplementary Figure SN2.SF3. The Data Table.**

Appears when clicking on “Data” in the top navigation bar. The table provides persistent access to all pipeline run results as well as user uploaded data. Metadata about each file in the table is provided in the Info Panel on the right. Tools are also available to tag and add notes to data to facilitate data management.

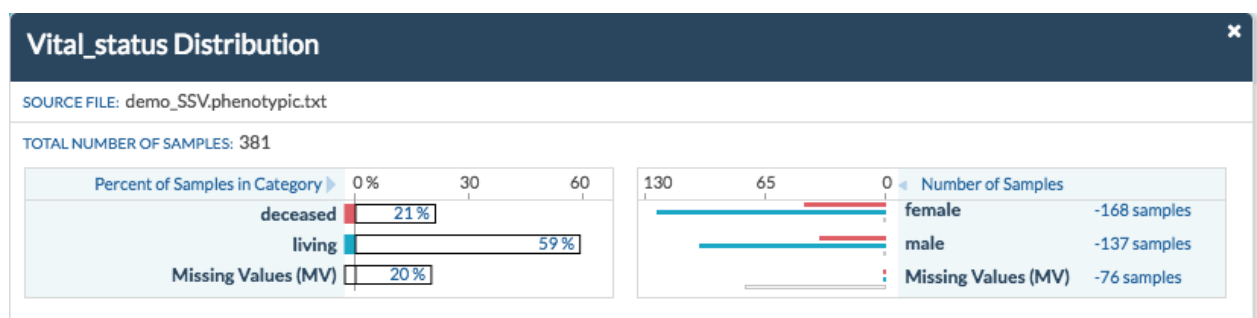

**Supplementary Figure SN2.SF4. Descriptive Statistics Panel.**

Available by clicking on elements in the Spreadsheet Visualization interface (as seen in Figure 2E). Allows users to explore the distribution of data across all of the variables available.

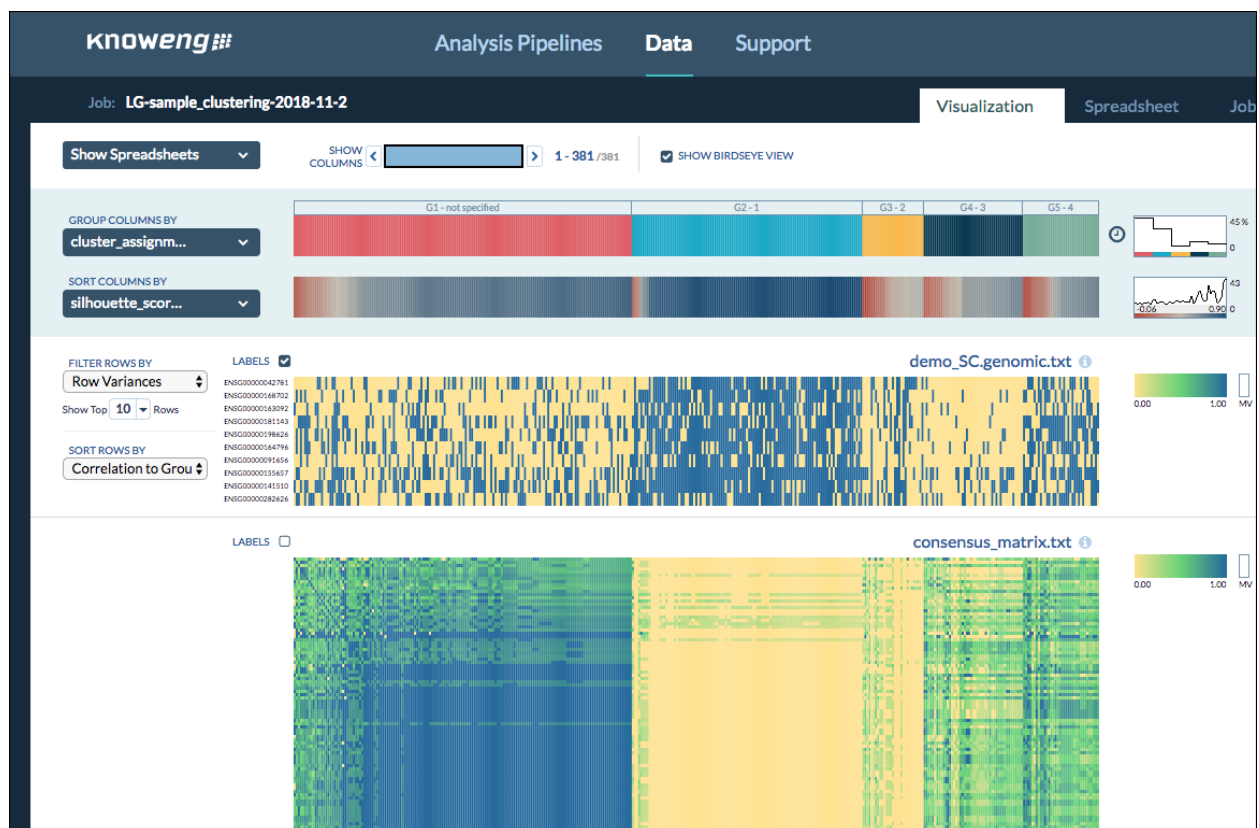

#### Supplementary Figure SN2.SF5. Sample Clustering Visualization.

Offers several different heat maps that can be compared across columns for possible correlations among omics features and clinical features. The top two bars are “data-strips” that organize the columns by user-selected features.

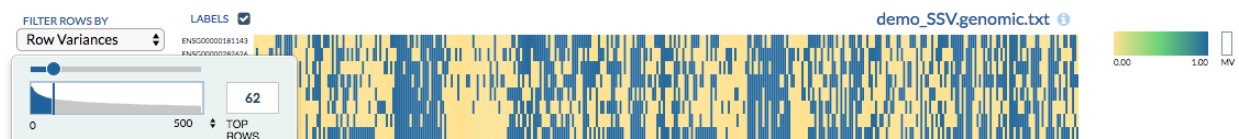

#### Supplementary Figure SN2.SF6. Top Feature Selector.

Helps user make an informed decision about the number of omics features to display in the visualization. It is used in the Spreadsheet, Sample Clustering, and Gene Prioritization Visualizations.

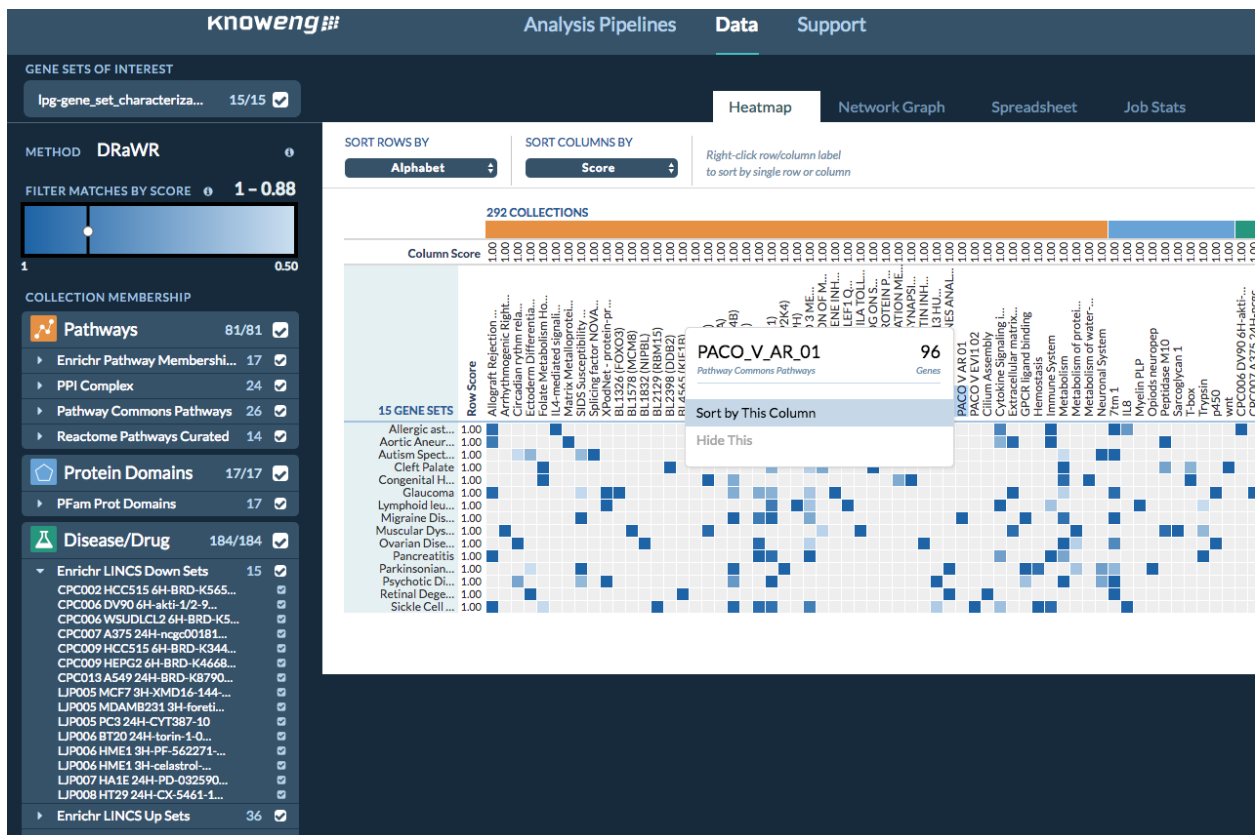

**Supplementary Figure SN2.SF7. Gene Set Characterization Visualization Tool.**

Visualizes degree of overlap of user gene set(s) with public gene sets that are incorporated into the platform.

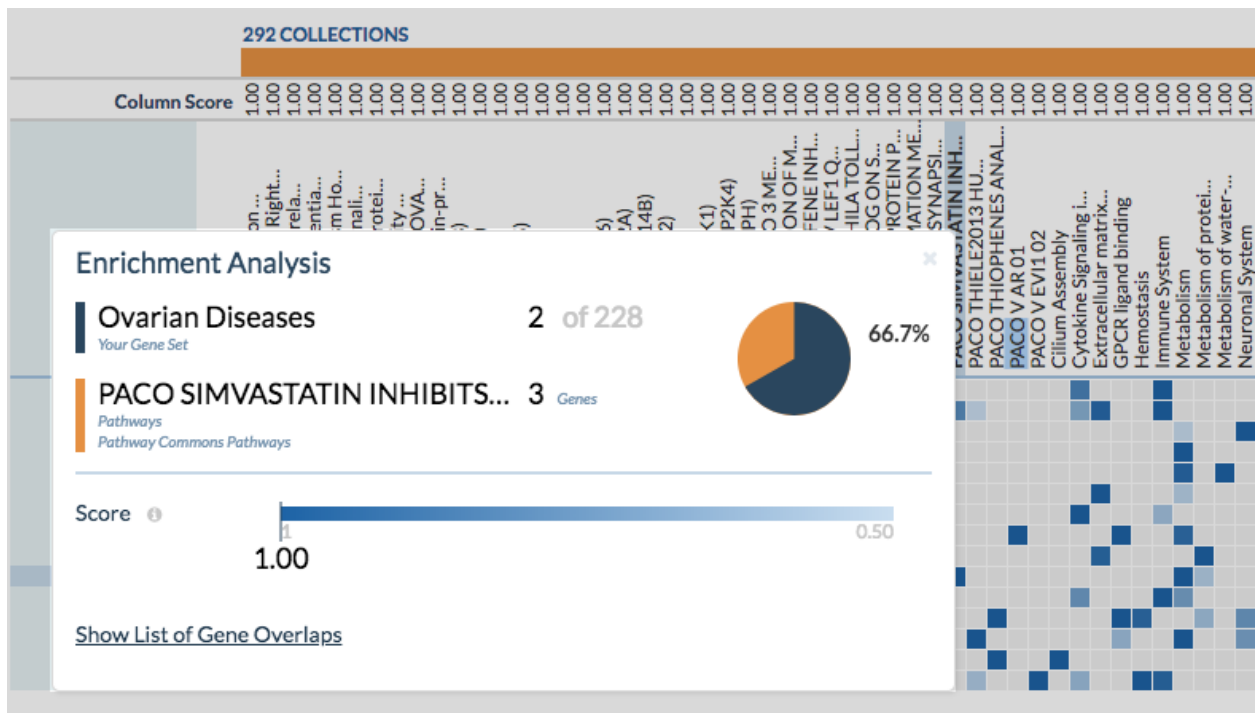

**Supplementary Figure SN2.SF8. Enrichment Analysis Drill Down Panel.**

Calculates and visualizes the degree of overlap between the user gene set and the selected public gene set.

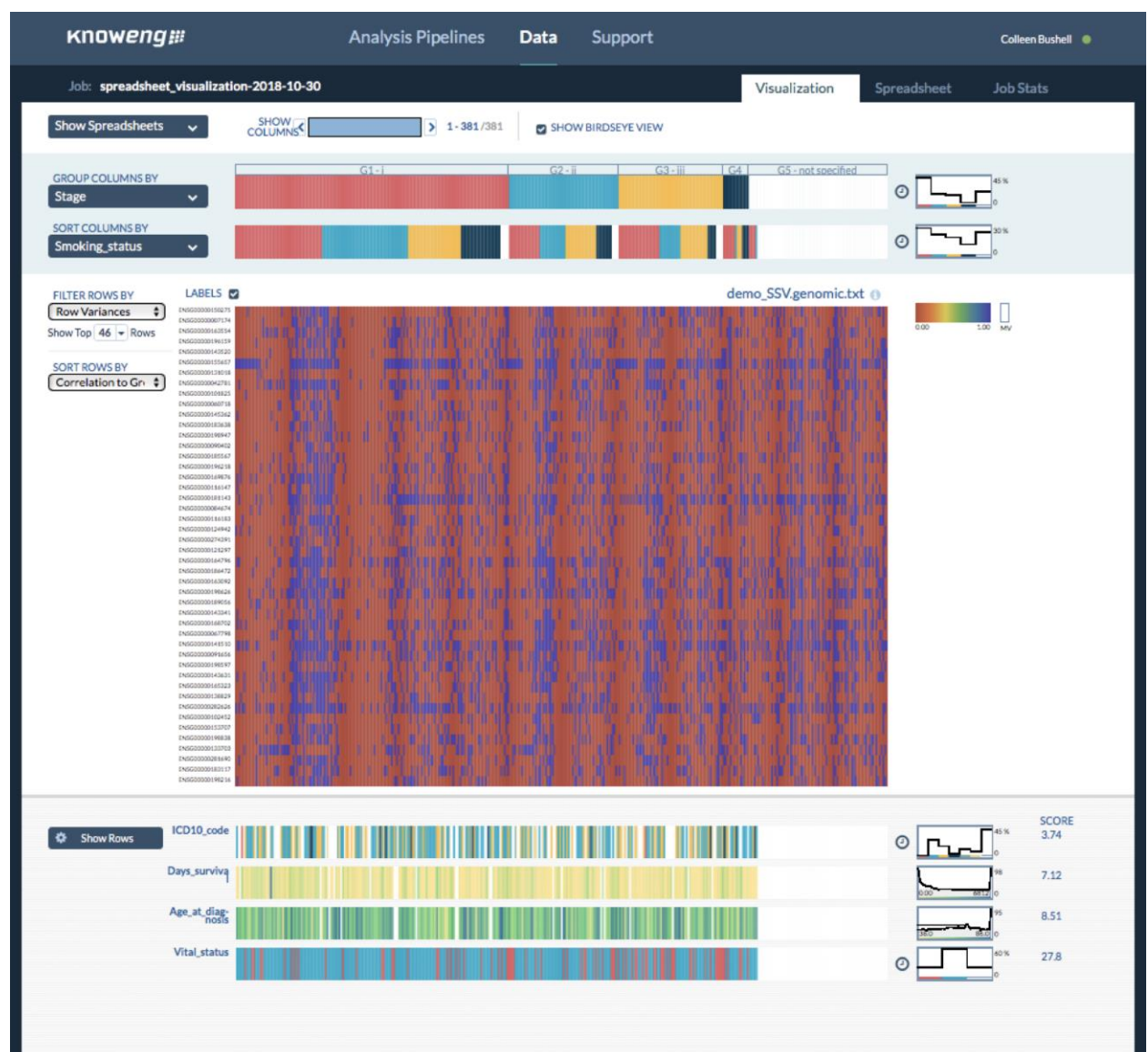

Supplementary Figure SN8.SF1. Spreadsheet Visualizer Interface.

**knoweng** Analysis Pipelines Data Support Lisa Gatzke

Pipeline: **spreadsheet\_visualization-2018-12-12** Data Review & Submission Cancel

Reset Defaults

Select one or more spreadsheets. Use Demo Data Upload New Data

**PROJECT** **FILTER BY**

Default Project All Recent Favorites

| SELECT | NAME | DATE/TIME | SIZE | PROJECT | STATUS |
| --- | --- | --- | --- | --- | --- |
|  | Demo2_Clinical_pancan12_30.txt | 11/2/2018, 1:37 PM | 26.4 KB | Default Project |  |
|  | Demo2_Mutation_pancan12_30.tsv | 11/2/2018, 1:37 PM | 20.6 MB | Default Project |  |
|  | demo_FPgenomic.txt | 11/12/2018, 11:22 AM | 71.0 MB | Default Project |  |
|  | demo_FPphenotypic.txt | 11/12/2018, 11:23 AM | 112.3 KB | Default Project |  |
|  | demo_GSC.spreadsheet.txt | 11/2/2018, 2:59 PM | 635.3 KB | Default Project |  |
|  | demo_SA.samples.txt | 11/12/2018, 1:37 PM | 281.5 MB | Default Project |  |
|  | demo_SA.signatures.mapped.txt | 11/12/2018, 1:37 PM | 17.8 KB | Default Project |  |
|  | demo_SC.genomic.txt | 11/1/2018, 5:08 PM | 11.6 MB | Default Project |  |
|  | demo_SC.phenotypic.txt | 11/1/2018, 5:08 PM | 26.0 KB | Default Project |  |
|  | demo_SSV.genomic.txt | 11/2/2018, 11:01 AM | 11.6 MB | Demo File |  |
|  | demo_SSV.phenotypic.txt | 11/2/2018, 11:01 AM | 26.0 KB | Demo File |  |
|  | HiSeqV2_normalized_xx.tsv | 11/8/2018, 2:12 PM | 44.2 MB | Default Project |  |
|  | HiSeqV2_percentile_xx.tsv | 11/8/2018, 2:12 PM | 25.4 MB | Default Project |  |
|  | HiSeqV2_xx.tsv | 11/8/2018, 2:12 PM | 23.4 MB | Default Project |  |
|  | PANCAN_phenotype_limited_xx.ts... | 11/8/2018, 2:12 PM | 30.4 KB | Default Project |  |
|  | lpg_signature_analysis-2018-11... | 11/12/2018, 1:38 PM |  | Default Project |  |

**What should I use as spreadsheets?**

The spreadsheet visualizer can be run with multiple spreadsheets or with a single spreadsheet.

**Multiple Spreadsheets (recommended)**

When using multiple spreadsheets, each spreadsheet must represent the same sample population. Spreadsheet orientation does not matter—that is, samples can correspond to rows or to columns—as long as the sample identifiers are consistent between files and all files have at least 50% of their samples in common. It is best to separate different data types into their own spreadsheets. For example, if you have gene-expression data, somatic mutation data, and clinical data, the spreadsheet visualizer will be most effective if the data are organized as three spreadsheets, one containing the gene-expression data, one containing the somatic mutation data, and one containing the clinical data.

**Single Spreadsheet**

When using a single spreadsheet, it is best if the spreadsheet is oriented such that columns correspond to samples.

Next

#### Supplementary Figure SN8.SF2. Spreadsheet Visualizer Data Upload.

Used for selection of spreadsheets for visualization.

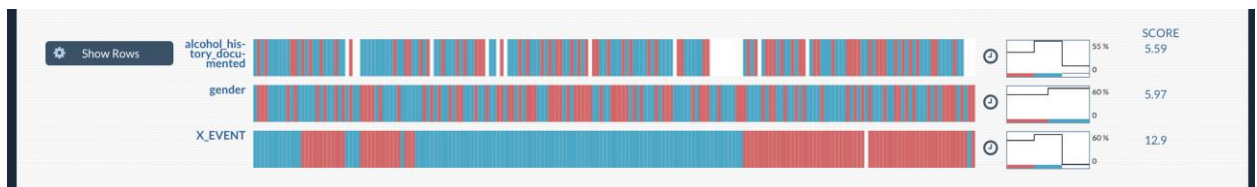

#### Supplementary Figure SN8.SF3. Spreadsheet Visualizer “Data-Strips”.

Use these to visually scan for correlations across the different spreadsheets.

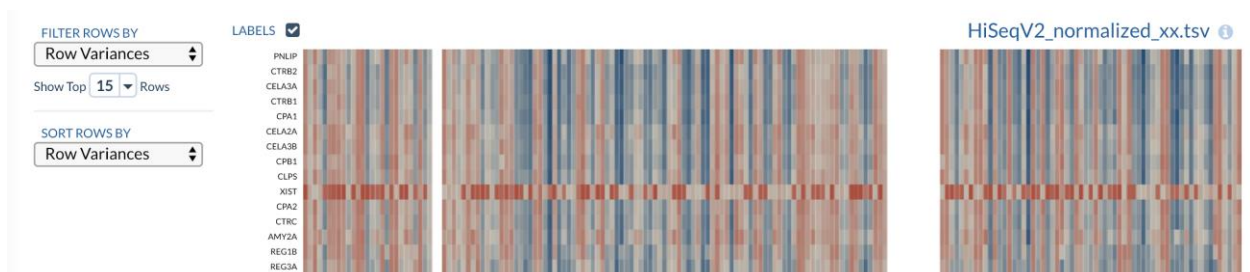

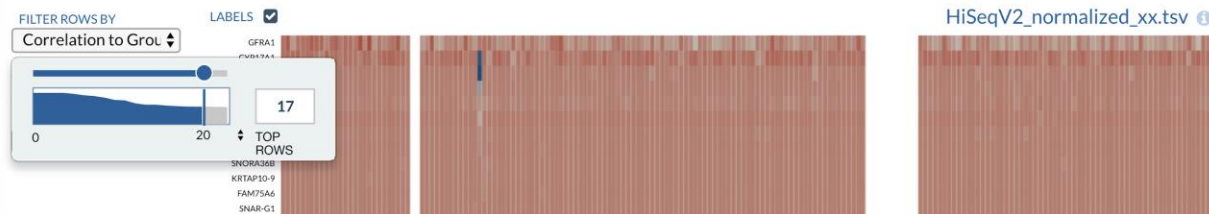

**Supplementary Figure SN8.SF4. Row Filtering and Sorting Controls.**

Rows can be filtered and sorted by Row Variance score as well as Correlation to the Group By category selected at the top.

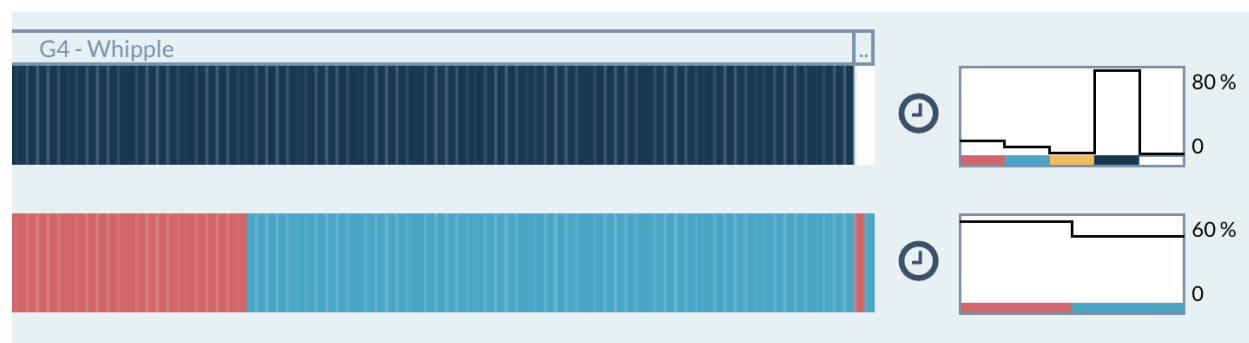

**Supplementary Figure SN8.SF5. Spreadsheet Visualizer Distribution Graphs.**

Spreadsheet Visualizer “data strips” distribution graphs shown on the right.

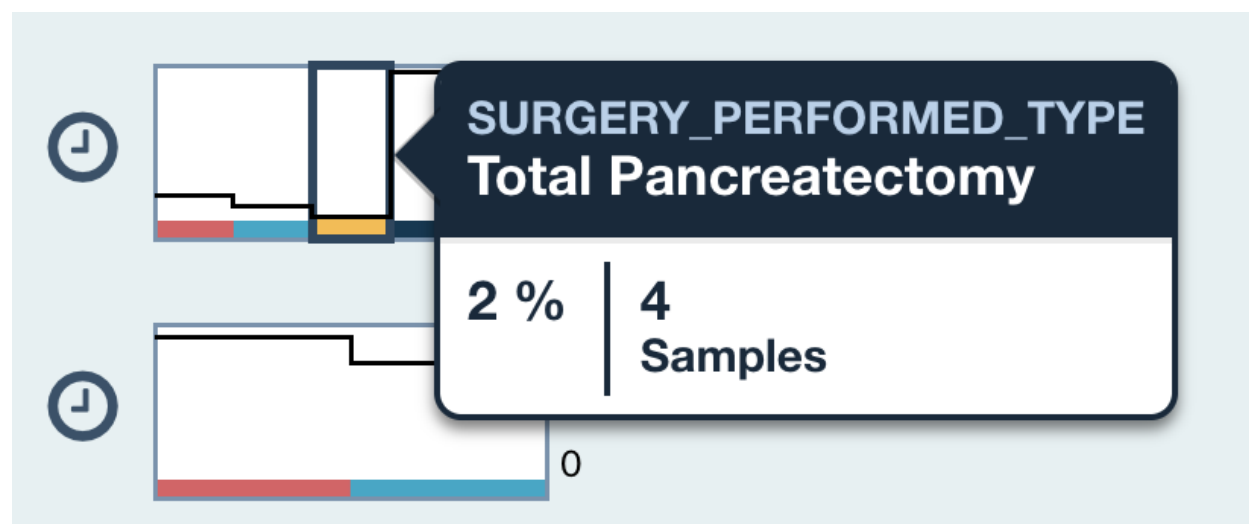

**Supplementary Figure SN8.SF6. Distribution Graph Hover Information.**

Spreadsheet Visualizer “data strips” distribution graphs provide additional statistical information on hover.

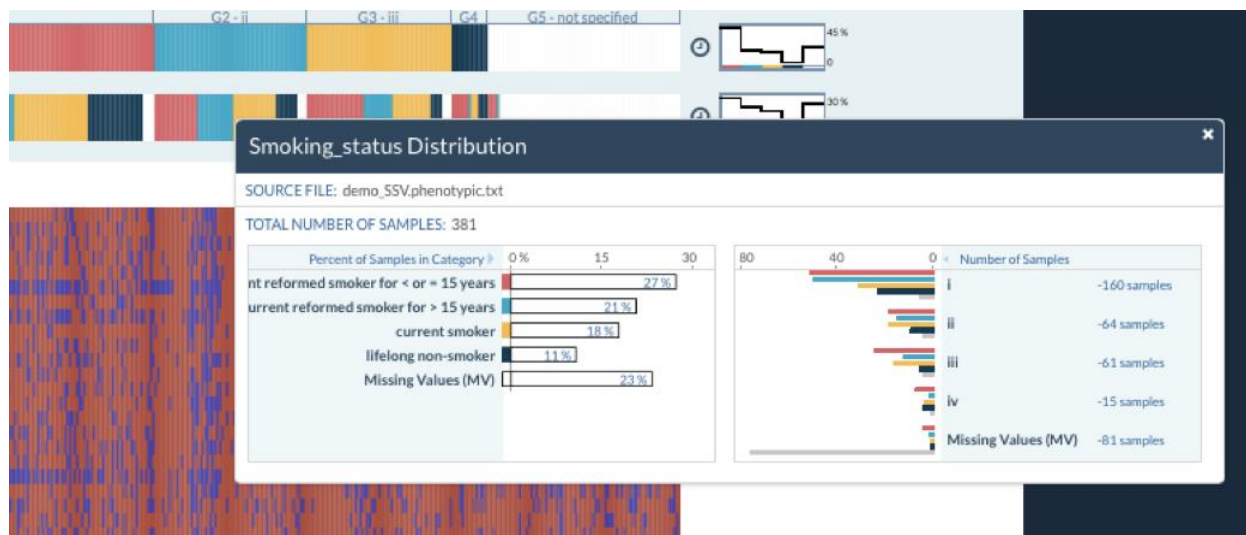

#### Supplementary Figure SN8.SF7. By Group Data Distribution.

Clicking on Spreadsheet Visualizer “Group by” and “Sort by” data strips triggers a popup showing the distribution of the data across the category bins as well as showing how these distribute across the “Group By” category selected at the top.

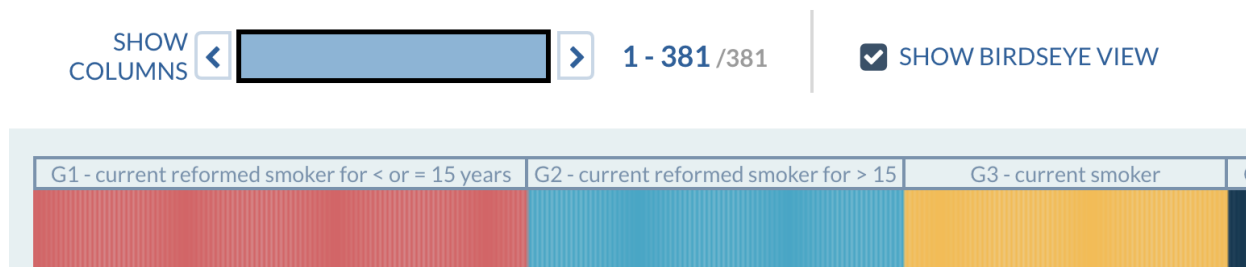

#### Supplementary Figure SN8.SF8. Birdseye View.

The Navigator controls help users control how many columns are in the display at a time. When there are many more columns than can be displayed on screen, “Show Birdseye View” can be selected. Cell colors will be blended to provide a sense of the data values on a single screen the user a sense of how all of the data looks without horizontal scrolling.

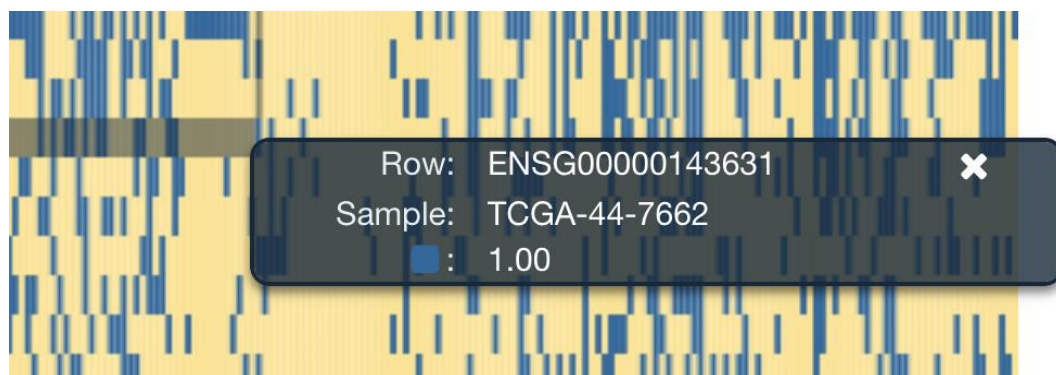

#### Supplementary Figure SN8.SF9. Data Cell Hover Over.

The Navigator controls help users control how many columns are in the display at a time. When there are many more columns than can be displayed on screen, “Show Birdseye View” can be selected. Cell colors will be blended to provide a sense of the data values on a single screen the user a sense of how all of the data looks without horizontal scrolling.

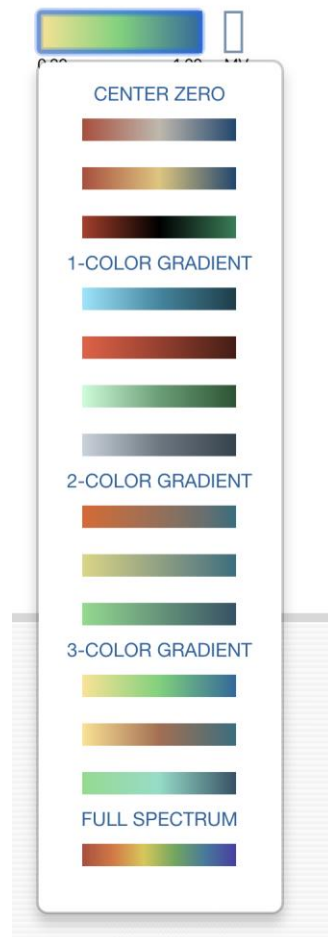

#### Supplementary Figure SN8.SF10. Color Scale Map.

A variety of color map choices are available and can be switched out on the fly depending on the needs of the data being displayed.

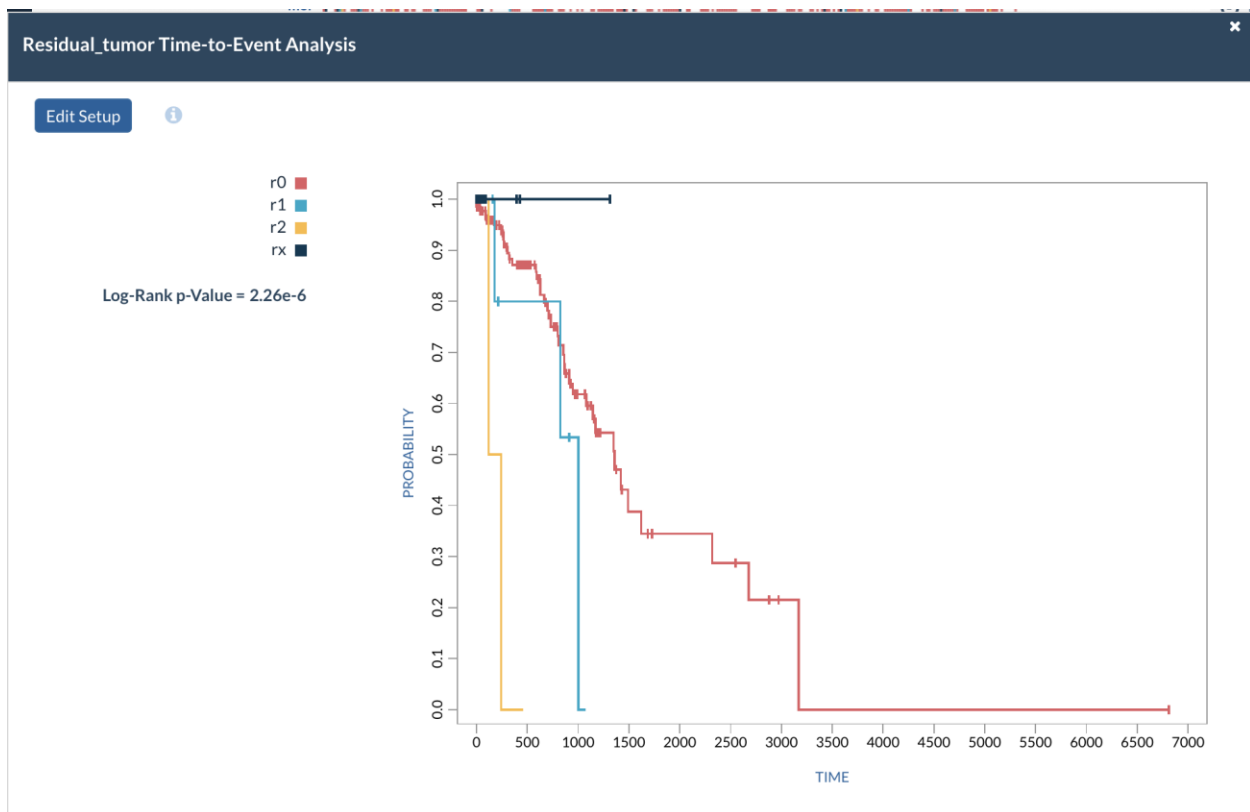

**Supplementary Figure SN8.SF11. Survival Curve.**

For spreadsheets that contain time-to-event measures, such as years of survival, the researcher can generate a Kaplan-Meier analysis for any categorical phenotype of interest.

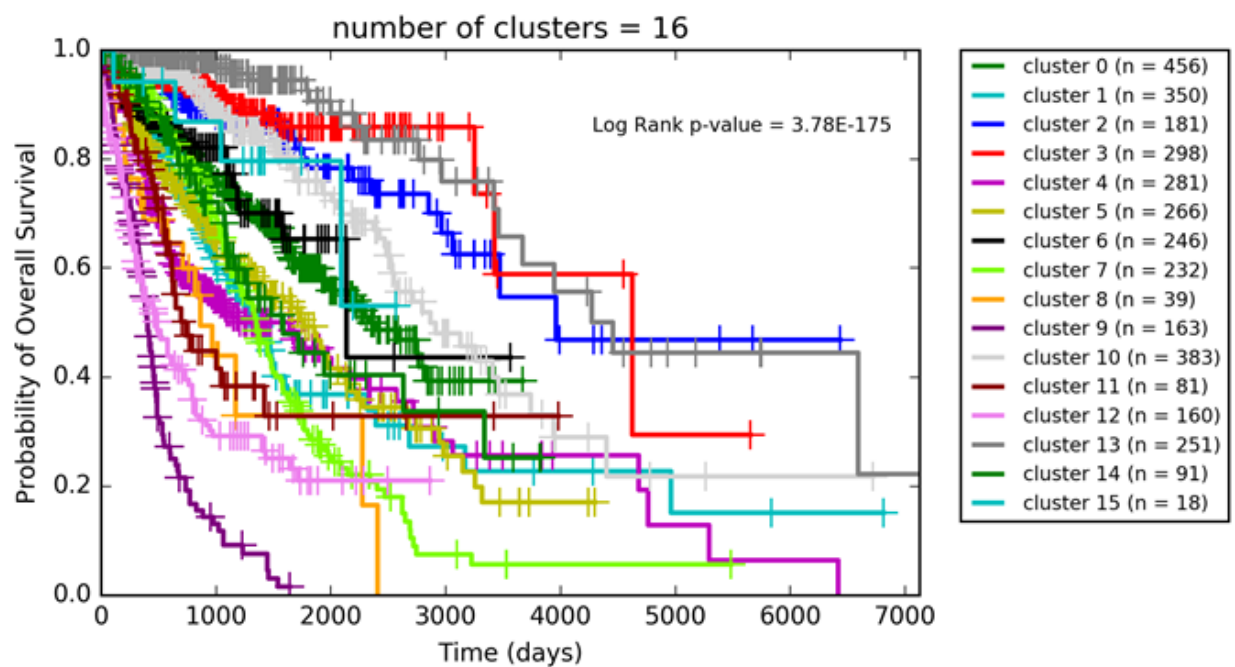

**Supplementary Figure SN9.SF1. Survival Analysis of “expr\_gp100\_hnInt” Clustering.**
Each cluster is plotted as a separate survival curve in the Kaplan-Meier plot and the p-value of the multivariate log rank test is reported.

#### Supplementary Figure SN10.SF1. Editing Tool on the SB-CGC.

This page shows the editing of the Gene Set Characterization command line tool on the SB-CGC. There are several tabs available to edit different aspects of a tool; this figure shows the **GENERAL** tab, which includes things like the command being run, its arguments, the docker container, the compute resources required, and associated files; other tabs include **INPUTS** and **OUTPUTS**.

Projects Data Public Apps Public projects Developer

mepstein

Dashboard Files Apps Tasks

KnowEnG\_GeneSetCharacterization\_Dev

Interactive Analysis Settings Notes

Revision 55 Edit with Rabin Composer Run

#### Gene Set Characterization Workflow

Created by mepstein on Sept. 14, 2017 13:50 • Last edited by mepstein on Mar. 23, 2018 16:29
Revision note: "Updated version of DCP."

##### Description

This KnowEnG Gene Set Characterization workflow tests a gene set for enrichment against a large compendium of annotations. This workflow starts with a user-submitted gene set (or multiple gene sets) and determines if each gene set is enriched for a pathway, a Gene Ontology term, or other types of annotations. This pipeline tests your gene set for enrichment against a large compendium of annotations. Gene Set Characterization can be done using a standard statistical test or in a Knowledge Network-guided mode (using DRaWR).

A network-guided analysis can offer various benefits over a standard one, including considering not just significant genes but also their network neighbors, and inferring properties of poorly annotated genes.

##### Required inputs

This workflow has one required input file:

1. Genomic Spreadsheet File (ID: *genomic\_spreadsheet\_file*). This currently must be a TSV file (a spreadsheet with tab-separated values). The first row (header) should contain the names of the gene sets in the corresponding columns. The first column of the spreadsheet should be the gene identifiers corresponding to each row. For each entry in the spreadsheet table, a "1" indicates that the corresponding row gene is part of the corresponding column gene set, a "0" means it is not. There should be no NA values/empty cells.

A sample input file, demo\_GSC.spreadsheet.txt, as described in the quickstart guide for this workflow, is available.

Example of Genomic Spreadsheet File Format:

|  | GeneSet1 | GeneSet2 | GeneSet3 |
| --- | --- | --- | --- |
| Gene1 | 0 | 1 | 1 |
| Gene2 | 0 | 0 | 0 |
| Gene3 | 1 | 0 | 1 |
| Gene4 | 0 | 1 | 0 |
| Gene5 | 1 | 0 | 0 |

##### Basic Information

CWL Version: sbg:draft-2

Contributors: mepstein

Toolkit: KnowEnG\_CGC v1.0

License: Copyright (c) 2017, University of Illinois Board of Trustees; All rights reserved.

Category: Analysis, Characterization, Enrichment

App ID: mepstein/genesetcharacterization/gsc-workflow

Links: KnowEnG Main Website, KnowEnG Analytics, Knowledge Network Overview, Knowledge-Guided Pipelines, GSC Pipeline, Pipeline Quickstart Guides, GSC Pipeline Quickstart, CGC GSC Pipeline Quickstart, KnowEnG YouTube Channel

##### Workflow steps

Join Names >

Knowledge Network Fetcher >

Knowledge Network Fetcher >

Gene Set Characterization Parameters >

Gene Set Characterization >

### Supplementary Figure SN10.SF2. The GSC Workflow Description.

This page can be seen at [https://cgsc.sbggenomics.com/public/apps#mepstein/knoweng-genesetcharacterization-public/gene-set-characterization/]. The Description is the documentation for an app; conventionally, it includes a variety of information about the app, including a general description of the app, how it is used, and its inputs and outputs. It also includes a specification of a sample command line (for command line tools) or an image of the workflow.

#### Supplementary Figure SN10.SF3. Editing the GSC Workflow.

Editing a workflow on the SB-CCG primarily consists of creating a graphical image of the workflow (using drag-and-drop); the nodes to the left are the inputs to the workflow, those to the right are the outputs, and those in the middle are the component apps of the workflow. In this figure, where one of the apps is selected, the fields to the right show the inputs for that app; hard-coded values can be set there, or inputs can be ported and made inputs for the overall workflow.

Public Apps Developer Login

Public apps

Category
Toolkit
Reset search

**KnowEnG Analysis of LUSC Subtypes**

KnowEnG\_CGC v1.0

This workflow is a compilation of other KnowEnG workflows. It takes some input gene expression...

Run

**Gene Prioritization Workflow**
KnowEnG\_CGC v1.0

This KnowEnG Gene Prioritization workflow identifies genes whose genomic measurements are most...

ANALYSIS PRIORITIZATION Run

**Gene Set Characterization Workflow**
KnowEnG\_CGC v1.0

This KnowEnG Gene Set Characterization workflow tests a gene set for enrichment against a larg...

ANALYSIS CHARACTERIZATION ENRICHMENT Run

**Samples Clustering Workflow**
KnowEnG\_CGC v1.0

This KnowEnG workflow performs sample clustering on a spreadsheet of "omics" data. With this ...

ANALYSIS Run

**Signature Analysis Workflow**
KnowEnG\_CGC v1.0

This KnowEnG workflow matches genomic profiles from a user-supplied query spreadsheet to a lib...

ANALYSIS Run

**Spreadsheet Builder**
KnowEnG\_CGC v1.0

This KnowEnG workflow processes CGC input files with genomic data and generates genomic and ph...

CONVERTERS Run

Showing 6 of 6

#### Supplementary Figure SN10.SF4. KnowEnG Public Apps on the SB-CGC.

This page can be seen at [https://cgc.sbgenomics.com/public/apps#q?search=knoweng]. It is easy to search through the many public apps at the SB-CGC. Currently, KnowEnG has published six apps there.

**Supplementary Figure SN10.SF5. CWL for the GSC Tool.**
CWL can be in JSON or YAML format, and includes description of the inputs and outputs of the app, and the specification of the command line (e.g., the base command and arguments) for command line tools or the steps in the workflow.

Projects

Data

Public Apps

Public projects

Developer

Dashboard

Files

Apps

Tasks

Lung\_Signatures\_Public

Interactive Analysis

Settings

Notes

KnowEnG Analysis of LUSC Subtypes

KNOWENG

Created by mepstein on July 10, 2018 15:05 • Last edited by mepstein on Sept. 4, 2018 14:25

Revision note: "Updated documentation on some inputs."

Revision 7

Edit with Rabin Composer

Run

Basic Information

CWL Version

sbg:draft-2

Contributors:

mepstein

Toolkit:

KnowEnG\_CGC v1.0

App Id:

mepstein/lung/knoweng-analysis-of-lusc-subtypes

Workflow steps >

Spreadsheet Builder >

Signature Analysis Workflow >

Gene Prioritization Workflow >

Gene Set Characterization Workflow >

Gene Set Characterization Workflow >

Description

This workflow is a compilation of other KnowEnG workflows. It takes some input gene expressions files (typically TCGA), and uses the Spreadsheet Builder workflow to create genomic and phenotypic spreadsheet files. The genomic file, along with an input signatures file, is passed to the Signature Analysis workflow, which outputs a binary similarity matrix. This binary matrix, along with the genomic file, is passed to the Gene Prioritization workflow, to create a top genes file output. This top genes file, along with property gene set network information, is then passed to two Gene Set Characterization workflows, one running in DRaWR mode (with an associated Interaction Knowledge Network) and one in Fisher mode (no knowledge network).

Ports

Inputs

App Settings

Outputs

### Supplementary Figure SN10.SF6. Workflow for the Analysis of LUSC Subtypes.

This page can be seen at

https://cgc.sbggenomics.com/public/apps#mepstein/lung/knoweng-analysis-of-lusc-subtypes/.

The documentation for the KnowEnG Analysis of LUSC Subtypes (KALS) workflow app on the SB-CGC.

20

Projects

Data

Public Apps

Public projects

Developer

Dashboard

Files

Apps

Tasks

Lung\_Signatures\_Public

Interactive Analysis

Settings

Notes

KnowEnG Analysis of LUSC Subtypes

Revision: 7

Want to try our new desktop editor, the Rabix Composer?

Save

Run

Input Files

Filter Minimum Percen.

Filter Threshold

Metadata Sample ID

Normalize Flag

Signatures File

Similarity Measure

Number of Top Genes

Gene Set Property Network Edge Type

Knowledge Network I

Amount of Network Influence

Spreadsheet Builder

Signature Analysis Workflow

Gene Prioritization Workflow

Gene Set Characterization Workflow

Gene Set Characterization Workflow

Clean Genomic Matrix

Clean Phenotypic Matrix

GSF DBaWR Results

Top Genes File

GSF Fisher Results

APPS

PARAMS

Gene Set Characterization Wo...

Uncategorized

taxonid

9606

pg\_edge\_type

network\_smoothing\_percent

gg\_edge\_type

+

-

Forum

Terms

Policies

Data Use

© 2018 Seven Bridges Genomics

### Supplementary Figure SN10.SF7. Editing the KALS Workflow.

The KALS workflow combines the Spreadsheet Builder workflow, the Signature Analysis workflow, the Gene Prioritization workflow, and two instances of the Gene Set Characterization workflow.
